# CheckM2: a rapid, scalable and accurate tool for assessing microbial genome quality using machine learning

**DOI:** 10.1101/2022.07.11.499243

**Authors:** Alex Chklovski, Donovan H. Parks, Ben J. Woodcroft, Gene W. Tyson

**Affiliations:** Centre for Microbiome Research, School of Biomedical Sciences, Queensland University of Technology (QUT), Translational Research Institute, Woolloongabba, Queensland, Australia; Donovan Parks, Bioinformatic Consultant, Castlegar, British Columbia, Canada

## Abstract

Advances in DNA sequencing and bioinformatics have dramatically increased the rate of recovery of microbial genomes from metagenomic data. Assessing the quality of metagenome-assembled genomes (MAGs) is a critical step prior to downstream analysis. Here, we present CheckM2, an improved method of predicting the completeness and contamination of MAGs using machine learning. We demonstrate the effectiveness of CheckM2 on synthetic and experimental data, and show that it outperforms the original version of CheckM in predicting MAG quality. CheckM2 is substantially faster than CheckM and its database can be rapidly updated with new high-quality reference genomes. We show that CheckM2 accurately predicts genome quality for MAGs from novel lineages, even those with sparse genomic representation, or reduced genome size (e.g. symbionts) such as those found in the Patescibacteria and the DPANN superphylum. CheckM2 provides accurate genome quality predictions across the microbial tree of life, giving increased confidence when inferring novel biological conclusions from MAGs.

---

Large-scale sequencing and assembly of genomes directly from environmental samples has led to the recovery of hundreds of thousands of highly diverse metagenome-assembled genomes (MAGs) from metagenomic data^1–3^, making it impractical to manually assess the quality of these genomes. The original approach to this problem used by CheckM (hereafter CheckM1)^4^ and other similar tools (such as BUSCO^7^) is to identify single-copy, near-universal marker genes associated with specific lineages to predict genome completeness and contamination. However, this approach has a number of limitations.

The single-copy marker gene approach used by CheckM1 relies on comparative genomics to identify lineage-specific marker gene sets to predict the completeness and contamination of a recovered MAG based on their presence, absence and copy number. Well-studied lineages with many high-quality genomes usually have more robust marker sets, which allows for higher accuracy and confidence in genome quality predictions. For novel lineages that lack high-quality genomic representation, only the most general marker sets (e.g. domain-level) can be used for genome quality estimates, resulting in reduced accuracy and sensitivity. In addition, this approach typically performs poorly on MAGs from microorganisms with reduced genomes, which lack some ‘universal’ marker genes^6^, and in many instances do not have many high-quality genomic representatives to derive robust marker sets.

An alternative approach to this problem is to use more complex mathematical techniques such as machine learning to link a wider range of genomic inputs to predict genome quality. Machine learning (ML) algorithms can generate insights into complex data and have been used for important biological challenges such as protein folding^7^ and metagenomic binning^8^. The application of ML to estimating genome quality has a number of advantages as it allows the incorporation of additional genomic information such as multi-copy genes, biological pathways and modules, and other genomic features such as amino acid counts and number of coding sequences. Furthermore, it allows for automatic selection of relevant genomic features to use for genome quality predictions without relying on a pre-defined lineage-specific marker sets.

Here we introduce CheckM2, a machine learning-based tool for predicting isolate, single-cell and MAG quality. CheckM2 builds models suitable for predicting bacterial and archaeal genome completeness and contamination without explicitly considering taxonomic information. CheckM2 was trained on simulated genomes with known levels of completeness and contamination, benchmarked, and subsequently applied to MAGs from a range of different environments. Overall, CheckM2 outperformed CheckM1, and performed substantially better on MAGs from novel lineages, such as the Candidate Phyla Radiation (Patescibacteria) and DPANN superphylum, as well as other lineages with sparse or no genomic representation.

## Results

### CheckM2 genome simulation, training and benchmarking

To demonstrate that machine learning (ML) can be applied to accurately predict genome quality, synthetic metagenome-assembled genomes (MAGs) with known quality were constructed for ML training. A ‘random-protein-sampling’ method was used to build training MAG sets, where predicted proteins from a subset of 4,978 bacterial and 322 archaeal complete isolate genomes selected from NCBI RefSeq^9^ Release 89 were randomly sampled to build ∼700,000 synthetic MAGs at pre-determined completeness and contamination percentages (see Methods; **Figure 1a)**. To validate the performance of ML models, two separate MAG simulation approaches were used: i) a ‘20kb-nucleotide-fragmentation’ method where the full-length genomes were sheared into ∼20 kb-long pieces, and ii) a ‘MAG-derived-fragmentation’ model where full-length genomes were sheared into contig distributions representative of MAGs in the Genome Taxonomy Database (GTDB^10^; **Figure 1b)**. In both simulation models used for validation, contigs were randomly sampled to build MAGs with a range of simulated completeness (5%-100%) and contamination (0% - 100%) values.

**Figure 1:**
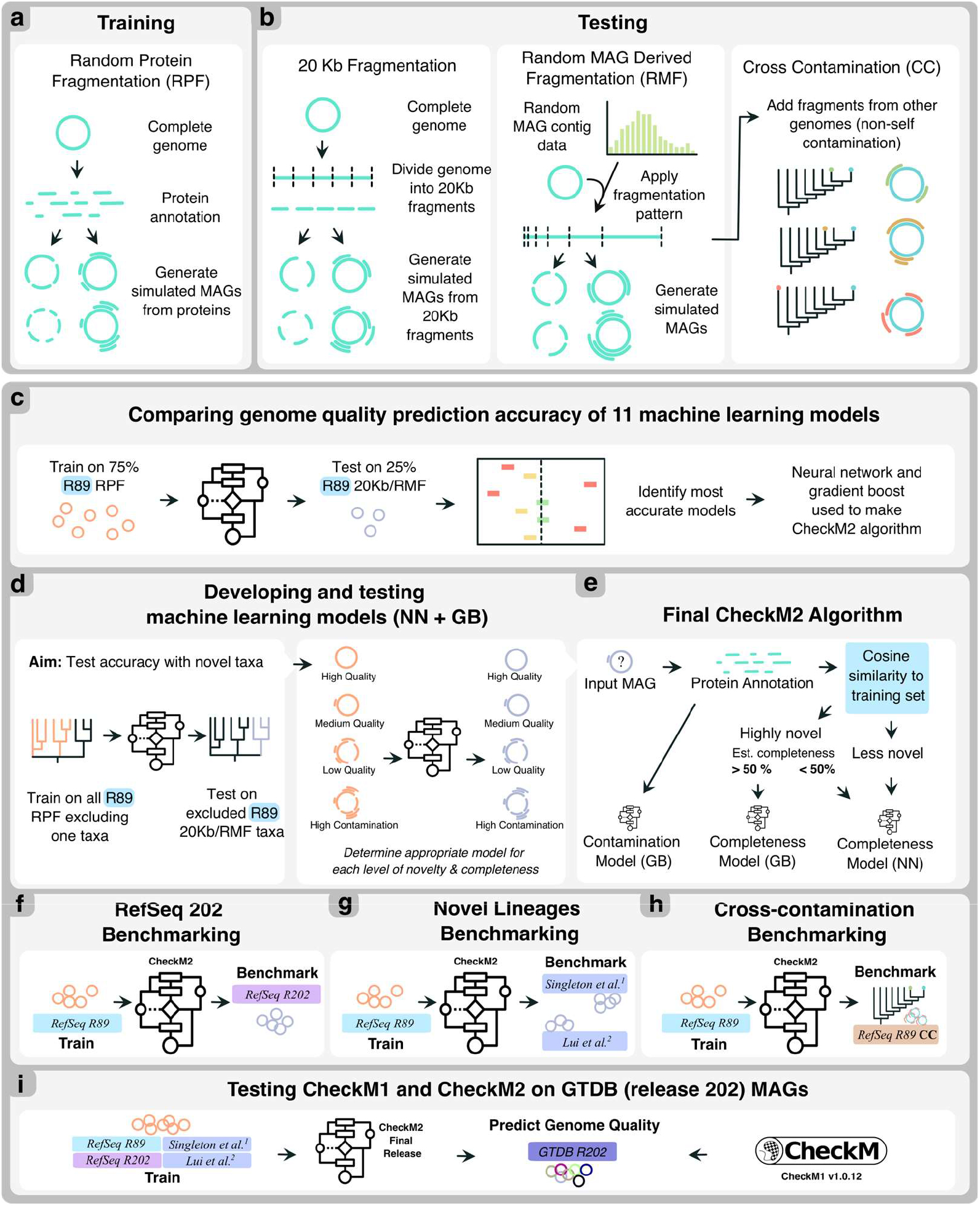
Overview of CheckM2 development, benchmarking and validation. **a)** Simulation of synthetic genomes for training and **b)** for testing, **c)** benchmarking the performance of 11 machine learning models on predicting genome quality in synthetic RefSeq r89 genomes, **d)** selection of neural network and gradient boost models and further testing and refinement, **e)** final algorithm used by CheckM2 to decide between gradient boost and neural network models, **f)** benchmarking of CheckM2 on RefSeq202 synthetic genomes, **g)** benchmarking of CheckM2 on novel and unusual synthetic genomes derived from circular MAGs including Patescibacteria, **h)** benchmarking CheckM2 on synthetic genomes with non-self contamination derived from RefSeq r89 genomes, and **i)** comparing CheckM1 and CheckM2 genome quality predictions for all GTDB r202 MAGs.

To train and test different ML models for predicting genome quality, the genome properties of synthetic MAGs were calculated as feature vectors for the ML models, including the genome length, number of coding sequences, and individual amino acid counts, as well as annotation of predicted proteins using KEGG^11^. In total, eleven machine learning methods (see Methods) were trained on randomly selected subsets of the simulated MAGs (75% of all MAGs; ‘random-protein-sampling’) and subsequently validated on the remainder of the MAGs (25%; for both ‘20kb-nucleotide-fragmentation’ and ‘MAG-derived-fragmentation’) for an initial assessment of quality prediction performance across diverse phyla (**Figure 1c**). To assess performance of ML models, predictions on simulated genomes were divided into four groups based on MIMAG completeness and contamination standards^12^ (*high quality*: >90% complete, <5% contaminated; *medium quality*: 50%-90% complete, <10% contaminated; *low quality*: <50% complete, <10% contaminated), as well as a separate group for *high contamination* (>10% contaminated); **Supplementary Table 1**.

Artificial neural networks^13^ (NN) and gradient boosted decision trees^14^ (GB) had the best overall performance **(Supplementary Table 2)** and were used for further optimisation and testing (**Figure 1c**). Both the NN and GB models exhibited higher accuracy when KEGG annotations were considered in the context of their pathways/modules (see Methods). In addition, the NN included convolutional layers for feature extraction, leading to an improvement in accuracy **(Supplementary Table 3, Supplementary Note 1)**. These optimisations to both models were used in subsequent testing.

### Assessment of ML models for predicting genome quality using simulated genomes

To assess the effect of taxonomic novelty on the accuracy of the optimized NN and GB models, an iterative leave-one-out approach was used on the synthetic genome set, where a specific taxonomic lineage was excluded from the training set and then each model tested for its performance on that lineage. The mean average error of both models for predicting completeness and contamination was systematically assessed from phylum to species level **(Figure 1d)**.

As expected, removing lineages from the training set with increasing taxonomic level proportionally affects the genome quality estimates (i.e. removing all genus-level representation has a substantially lower impact on accurately predicting genome quality than removing class-level or phylum-level representation of query genomes). Overall bacterial and archaeal genome quality estimates improve if the training set contains a genome that is more taxonomically related to the query genome **(Figure 2a)**. However, the two models have different strengths relative to genome novelty and genome completeness. Completeness quality estimates for query genomes representing novel phyla, classes, and orders were more accurate with the GB model, while the NN model was on average more accurate for genomes representing novel families, genera and species **(Figure 2a)**. Additionally, for low-quality (<50% complete) genomes the NN model was more accurate at all taxonomic levels, while the GB model accuracy declined with lower MAG quality **(Figure 2a)**.

**Figure 2.**
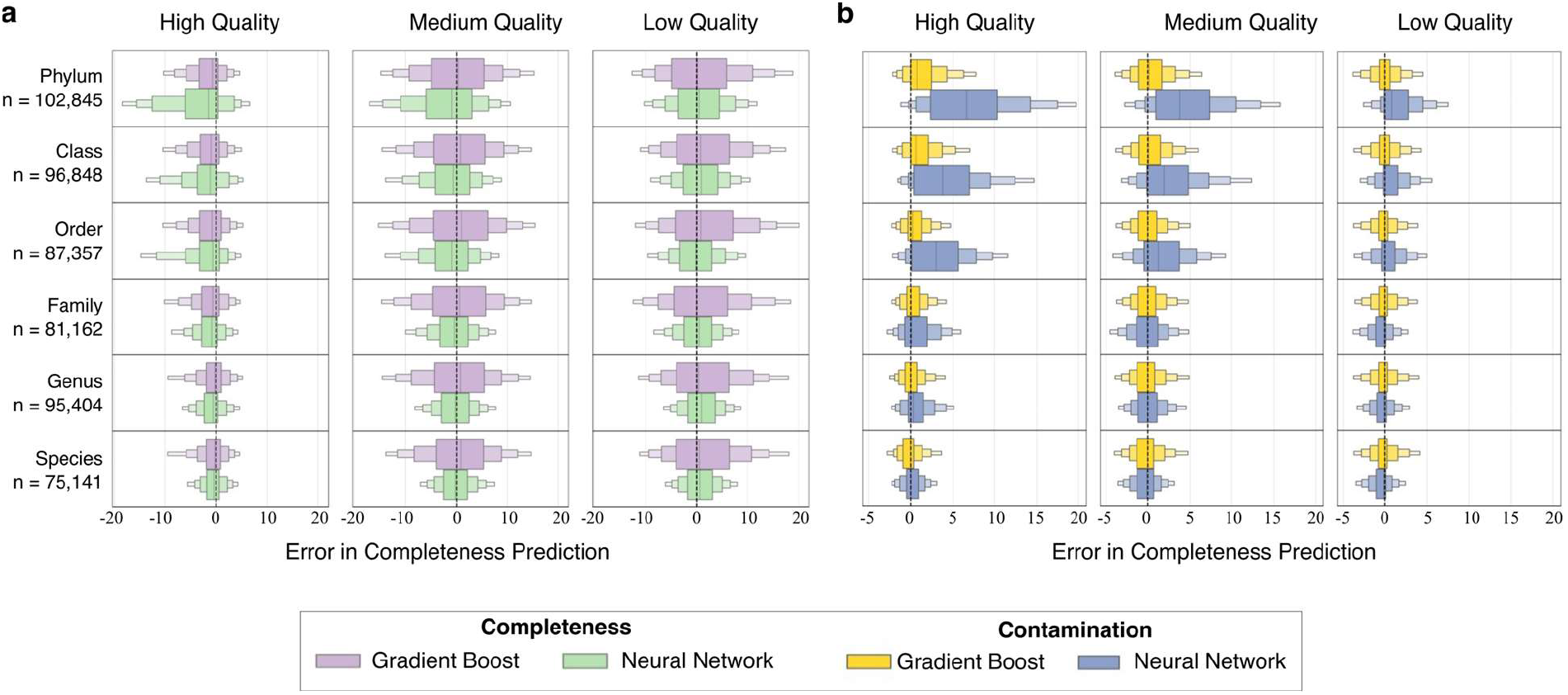
Comparison between neural network and gradient boost models on synthetic genomes derived from RefSeq release 89 to assess the effect of taxonomic novelty on accuracy. Using a leave-one-out approach, genomes from specific taxa were removed from the training set from phyla to species, models were trained on the remaining genomes, and prediction accuracy tested on the left-out group. **a)** Error in predicting completeness and **b)** contamination at varying taxonomic levels of novelty. Each taxonomic novelty level is broken into separate error margins across different MIMAG quality cut-offs (high quality: 90-100% completeness, 0-5% contamination; medium quality: 50-90% completeness and 0-10% contamination; and low quality: < 50% completeness, 0-10% contamination). Positive values indicate overestimation, while negative values indicate underestimation of true values. The size of each error box in the letter-value plot shows half the remaining data, starting with 50% for the first box, 25% for the second box, etc.

The most difficult completeness prediction scenario is likely to be genomes belonging to a new phylum (i.e. a phylum without a complete isolate genome). For near-complete genomes from a novel phylum, the mean average error (MAE) for completeness predictions using the GB model is **3.1±3.9%** and **5.2±5.7%** for the NN model. For medium quality genomes, the GB model had a MAE of **4.6±4.4%**, while the NN model had a MAE of **5.9±5.3%**. These results suggest the models have an ability to generalize to phylum-level novelty with relatively good accuracy even as genome quality declines. While it is impossible to reproduce this test for CheckM1, using CheckM1’s domain-level bacterial or archaeal marker sets consisting of ∼120 universal marker genes resulted in roughly equivalent MAEs of **3.4±4.4%** for high quality and **7.2±5.8%** for medium quality genomes.

For genome contamination predictions at all taxonomic levels, the gradient boost model substantially outperformed the neural network and was chosen as the model for predicting contamination **(Figure 2b; Figure 1e)**. For genomes belonging to a novel phylum, the predicted contamination MAE of the GB model is **2.0±2.2%** (HQ), contrasting with a NN MAE of **7.3±5.5%** (HQ) and a CheckM1 domain-level marker set comparison of **1.9±2.2%** (HQ).

The results of the simulation led to both models being implemented in the final version of CheckM2 for completeness prediction, while only the GB model was implemented to predict contamination. For completeness predictions on novel and more complete genomes, CheckM2 uses a ‘general’ model based on gradient boost decision tree algorithms, while for genomes more closely related to those in its reference set or less complete genomes, it uses a ‘specific’ model based on artificial neural networks **(Figure 1e)**. A cosine similarity measure was found to correlate well with input genome taxonomic novelty, with a linear relationship between squared cosine similarity and taxonomic distance **(Supplementary Figure 1)**, enabling CheckM2 to use this measure to select between the ‘general’ and ‘specific’ model for each input genome based on pre-defined cosine similarity cut-offs derived from the leave-one-out approach (See Methods; **Figure 1e; Supplementary Table 4)**.

### Benchmarking CheckM2 performance on new RefSeq genomes using simulated genomes

The initial CheckM2 ML models were built on genomes from RefSeq Release 89, allowing new complete genomes from RefSeq Release 202 to be used to test CheckM2’s performance, as they were not part of the original training and validation sets **(Figure 1f)**. In total, this included 2,864 new complete microbial isolate genomes representing 6 novel phyla, 13 novel classes, 43 novel orders, 87 new families, 439 novel genera and 1,554 novel species according to their GTDB classifications. As these genomes represent the range and types of genomes added to public databases over the course of ∼2 years, they provide a reasonable indication of how CheckM2 performs when tested against new genomes of varying taxonomic novelty. They also provide suitable complete genomes for simulating new genomes of known completeness and contamination (as in **Figure 1b)**, allowing benchmarking of CheckM2 against CheckM1.

When predicting the completeness of 712,880 simulated RefSeq 202-based genomes, CheckM2 was substantially more accurate than CheckM1 with a lower mean average error across all genomes **(Figure 3a; Supplementary Note 2)**. Overall, there was similar performance between CheckM2 and CheckM1 on high quality genomes (CheckM2 MAE: **2.1±2.9%**, CheckM1 MAE: **2.0±3.2%**), but CheckM2 was far more accurate for medium, low-quality and highly contaminated genomes (**Figure 3a**; CheckM2 MAE: **3.1±3.3%**, CheckM1 MAE: **4.7±5.4%**). However, as some phyla within RefSeq 202 are highly oversampled, bulk genome mean average error underestimates performance across broad taxonomic ranks. When using a phylum-weighted mean average error (PW-MAE), CheckM2 outperformed CheckM1 with both substantially higher accuracy and much lower error variance for both high quality genomes (CheckM2 PW-MAE: **2.5±1.4%**, CheckM1 PW-MAE: **5.7±8.2%**) as well as medium and low-quality genomes (CheckM2 PW-MAE: **3.9±1.1%**, CheckM1 PW-MAE: **7.1±4.8%**). CheckM2 exhibited comparable performance across both the ‘20-kb-fragmentation’ and the ‘MAG-derived random fragmentation’ simulations, suggesting that there is little effect of simulation method on resulting predictions **(Supplementary Note 2)**. The most significant increase in performance of CheckM2 was seen in predicting completeness of genomes from the phyla with very few high-quality genomic representatives such as Iainarchaeota, Nanohaloarchaeota, Dependentiae, Bipolaricaulota and Patescibacteria (high quality MAE: **3.7±3.1**) when compared to CheckM1 (high quality MAE: **26.3±10.7**) **(Figure 3b, 3c)**. Notably, there was only a single reference genome for Nanohaloarchaeota, Dependentiae, Bipolaricaulota and Iainarchaeota in the training set for CheckM2, indicating that a single genomic representative of a lineage provides sufficient information for an accurate prediction of genome quality.

**Figure 3.**
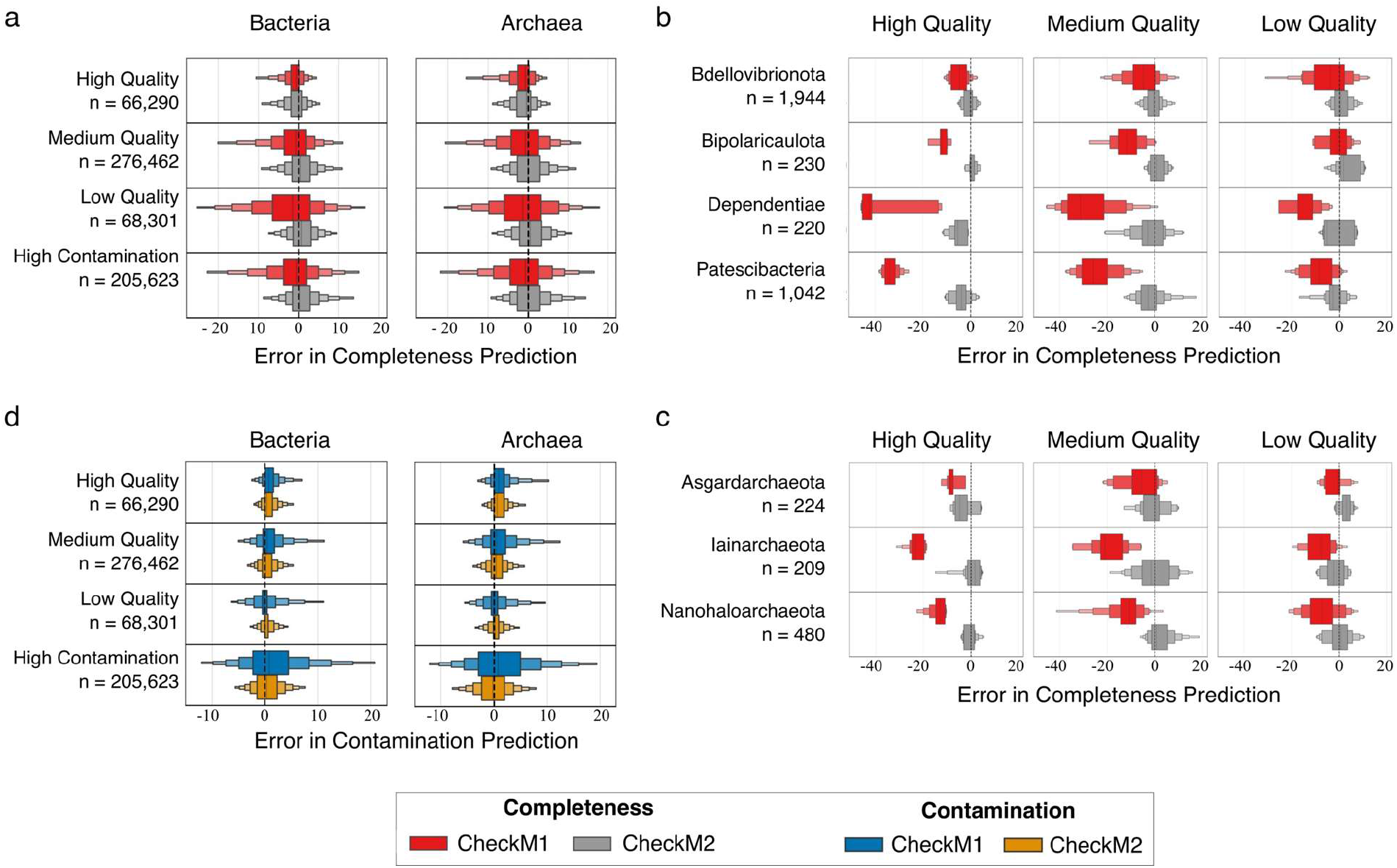
CheckM1 and CheckM2 completeness and contamination error on synthetic genomes derived from RefSeq r202 genomes. **a)** Error in predicting completeness on all bacterial and archaeal genomes, **b)** Error in predicting completeness on specific bacterial phyla and **c)** Error in predicting completeness on specific archaeal phyla with substantial differences between CheckM1 and CheckM2. **d)** Error in predicting contamination on all bacterial and archaeal genomes. Results are broken into separate error margins across different MIMAG quality cut-offs (high quality: 90-100% completeness, 0-5% contamination; medium quality: 50-90% completeness and 0-10% contamination; low quality: < 50% completeness, 0-10% contamination; and high contamination: > 10% completeness). Positive values indicate overestimation while negative values indicate underestimation of true values. The size of each error box in the letter-value plot shows half the remaining data, starting with 50% for the first box, 25% for the second box, etc.

When predicting contamination, CheckM2’s mean average error (MAE: **1.2±1.3**) was comparable to CheckM1 (MAE: **1.5±1.8**) on high quality genomes, and was substantially more accurate for medium and low quality genomes (CheckM2: **1.7±1.7%**, CheckM1: **3.0±4.0%**). It was also substantially better at predicting contamination in highly contaminated genomes **(Figure 3d)**.

### Benchmarking CheckM2 performance on DPANN and Patescibacteria

One of the weaknesses of CheckM1 was its poor performance on highly novel genomes, particularly those from the DPANN and Patescibacteria. The archaeal DPANN superphylum and the bacterial Patescibacteria are large radiations of microorganisms comprising a significant fraction of the tree of life^15^. Their high diversity, unusual biology, absence of key genes and often small genomes make predicting their genome quality particularly challenging^16–18^. To assess CheckM2’s ability to predict the quality of genomes from these microorganisms, 57 closed circular genomes were obtained from wastewater (Singleton et al., 2021)^19^, including 30 genomes from the Patescibacteria, as well as other highly novel and often small genomes from phyla such as Dependentiae, Iainarchaeota and UBA10199. Additionally, 36 other circularized Patescibacteria genomes were obtained from Lui et al., (2021)^20^. These lineages are poorly represented in the RefSeq Release 202, which mostly cover phyla and classes with existing isolate representatives. Together, this dataset represents 25 unique classes and 45 unique orders of curated and circular Patescibacteria genomes along with a number of other circularized MAGs representing novel phyla and classes, providing an excellent opportunity to test CheckM2’s performance on novel genomes (**Figure 1g**). From these complete genomes, simulated genomes of varying completeness and contamination were created as above **(Figure 1b)** to enable tool benchmarking across different levels of genome quality.

Across novel lineages CheckM2 is more accurate for both high-quality genomes (MAE: **2.9±2.6**) and medium/low quality genomes (MAE: **4.7±3.8**) than CheckM1 (high quality MAE: **4.7±6.0**; medium/low quality MAE: **6.1±5.7**), showing an ability to generalize well across a range of novel phyla and classes (**Figure 4a**). For novel lineages with reduced genomes such as the phyla Patescibacteria, Dependentiae or Iainarchaeota, where a specific marker set is not available for CheckM1, CheckM2 completeness predictions are far more accurate (MAE: **6.4±4.5**) than CheckM1 (MAE: **19.7±12.1**; **Figure 4b**). As with RefSeq 202 benchmarking, CheckM2 showed similar performance on genomes simulated by both test simulation methods and consistently outperformed CheckM1 (**Supplementary Note 3**).

**Figure 4.**
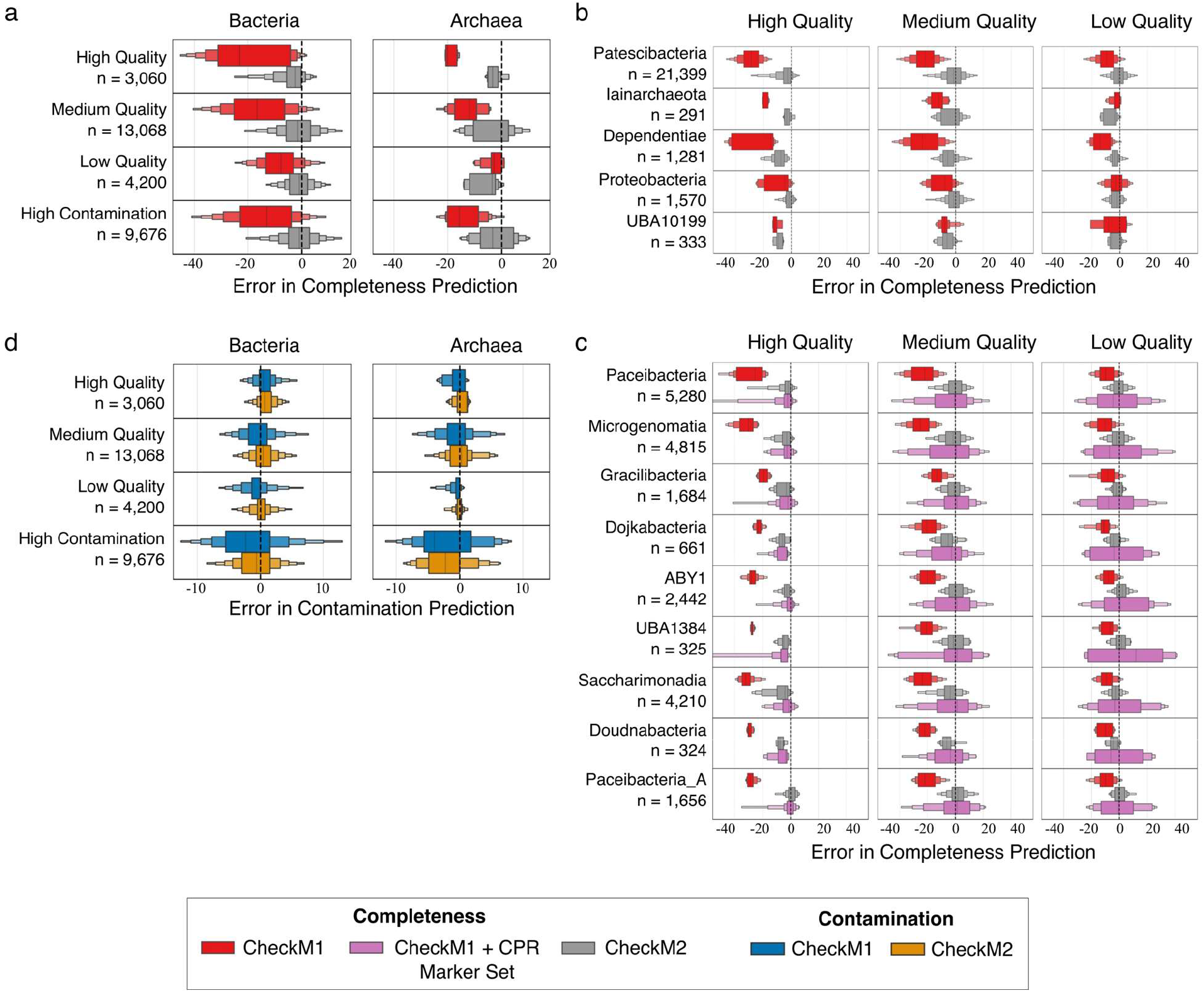
CheckM1 and CheckM2 completeness and contamination error on synthetic genomes derived from novel circular MAGs assembled by Singleton et al.^19^, and Lui et al.^20^ **a)** Error in predicting completeness on all bacterial and archaeal genomes, **b)** Error in predicting completeness on specific bacterial and archaeal phyla with substantial differences between CheckM1 and CheckM2, **c)** Error in predicting completeness on all classes of Patescibacteria, including predictions by CheckM1 using the CPR marker set, and **d)** Error in predicting contamination on all bacterial and archaeal genomes. Results are broken into separate error margins across different MIMAG quality cut-offs (high quality: 90-100% completeness, 0-5% contamination; medium quality: 50-90% completeness and 0-10% contamination; low quality: < 50% completeness, 0-10% contamination; and high contamination: > 10% completeness). Positive values indicate overestimation while negative values indicate underestimation of true values. The size of each error box in the letter-value plot shows half the remaining data, starting with 50% for the first box, 25% for the second box, etc.

Across all classes of Patescibacteria, CheckM2 was far more accurate than CheckM1, with CheckM1’s performance only being improved by using a custom CheckM1 Patescibacteria marker set based on 43 ribosomal genes^6^ **(Figure 4c)**. However, the accuracy of CheckM1 using the custom CPR marker set substantially declined on medium and low-quality Patescibacteria genomes with completeness error rates as high as 30-40%, making the method unreliable. The superior performance of CheckM2 extends to all Patescibacteria classes represented in the Singleton et al., and Lui et al.^20^, indicating that CheckM2 is able to robustly and accurately predict genome quality from highly diverse lineages such as the Patescibacteria, despite only having a few genomic representatives in the training set.

CheckM2 also outperformed CheckM1 on most cases of contamination (**Figure 4d**). The only partial exception are some high-quality Patescibacteria genomes where CheckM1’s lineage-specific marker sets provide slightly better accuracy (CheckM1 MAE: 1.4**±**1.1, CheckM2 MAE: 1.7**±**1.3; **Supplementary Note 3)**. In part this may be due to CheckM2’s conservative nature when approaching contamination predictions. However, it is likely that the addition of all new circularized Patescibacteria genomes to CheckM2’s final reference set (**Figure 1i**) will increase its accuracy.

### CheckM2 cross-contamination benchmarking using simulated genomes

Contamination in MAGs may come from the binning together of closely-related strain or species, but may potentially also contain divergent DNA from other lineages or even domains. CheckM1 uses duplicated single-copy marker gene counts to infer contamination, on the assumption that contamination will come from closely related genomes being binned together, and thus will contain duplicated single-copy genes. This is likely to work better when the sources of contamination are highly related and thus likely to share the same distribution of single-copy markers.

However, it is unclear how accurate CheckM1 is when assessing contamination from a different source than the same strain or species, and whether CheckM2’s weighted combination of feature vectors is better able to identify foreign contamination compared to only using duplicate single-copy marker genes. Therefore, both CheckM1 and CheckM2 were tested on simulated genomes with contamination originating from increasingly divergent sources, from same species to same domain (**Figure 1h**).

Results show that CheckM2 is accurate at identifying foreign contamination, particularly for high-quality genomes **(Figure 5)**. CheckM2 is highly reliable when predicting the contamination of highly complete genomes, though may somewhat underestimate such higher-taxa contamination for medium-quality genomes, as does CheckM1. Importantly, it is far less likely to overestimate contamination compared to CheckM1, likely due the fact it does not rely as strongly on single-copy marker genes and does not use small marker sets (**Supplementary Note 4**). CheckM2 should do particularly well on contamination from the same species, genus, or family which is considerably more difficult to detect with taxonomy-based detection tools such as GUNC^21^.

**Figure 5.**
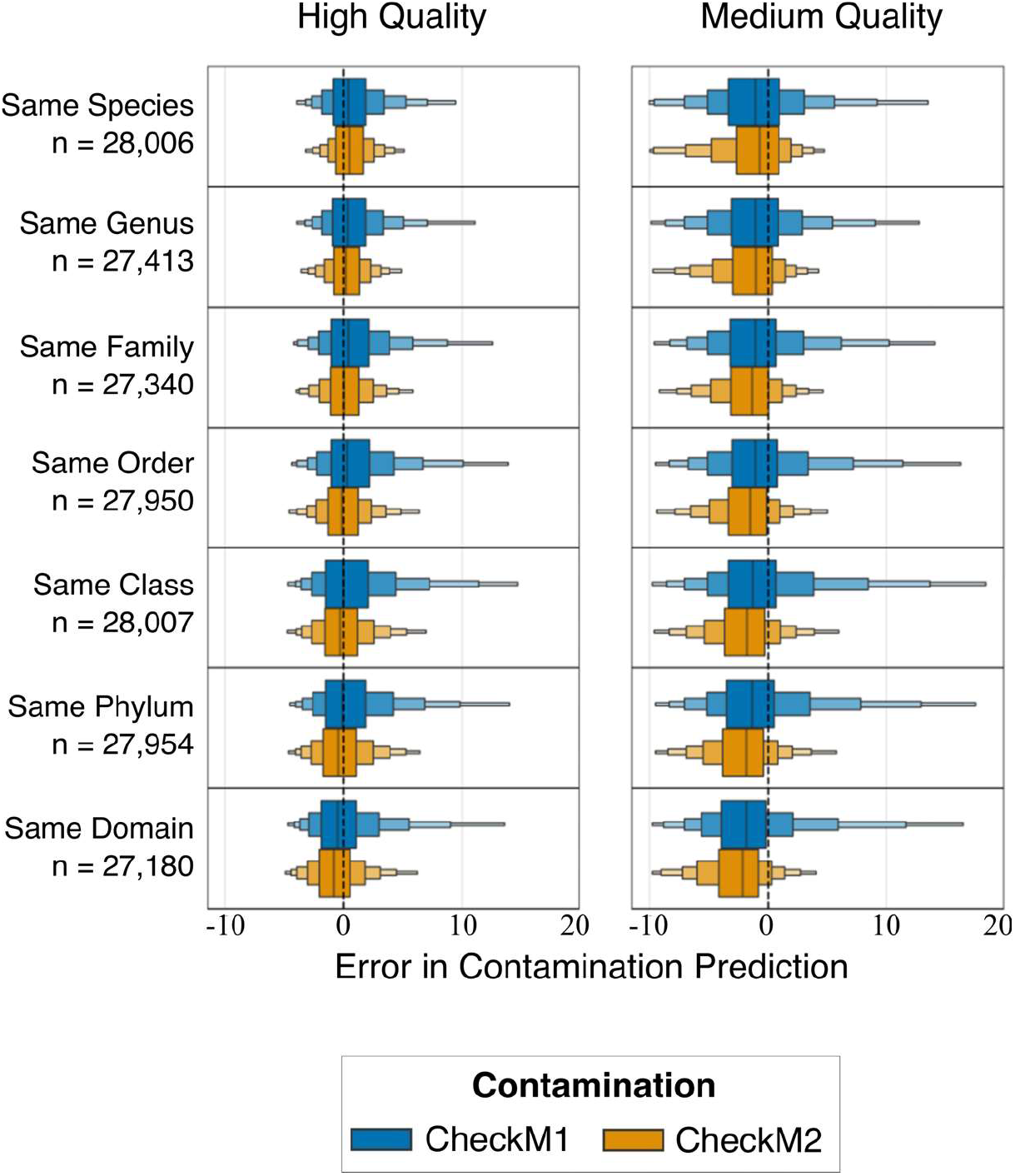
Contamination predictions by CheckM1 and CheckM2 on synthetic genomes derived from RefSeq r89 containing contamination from different taxonomic levels. Error in contamination prediction is broken by the taxonomic source of the contaminant relative to contaminated genome. Results are broken into separate error margins across different MIMAG quality cut-offs (high quality: 90-100% completeness, 0-5% contamination; medium quality: 50-90% completeness and 0-10% contamination. Positive values indicate overestimation while negative values indicate underestimation of true values. Each box shows half the remaining data, starting with 50% for the first and 25% for the second, etc.

### Application of CheckM2 to environmental MAGs

#### Comparison of CheckM1 vs CheckM2 predictions across all bacteria and archaea

Following benchmarking on synthetic genomes, CheckM2 was retrained with all complete genomes in RefSeq Release 202 to provide a comprehensive reference database for inclusion with the CheckM2 release version^1^. To systematically compare the genome quality predictions of CheckM1 and CheckM2 across all bacterial and archaeal lineages, both tools were used to predict the completeness and contamination of 224,101 bacterial and 3,881 archaeal genomes in the GTDB Release 202 annotated as ‘incomplete’ – i.e. not isolate genomes or closed circular MAGs **(Figure 1i)**.

Overall, there is good congruence among completeness predictions across most phyla between CheckM2 and CheckM1, with 73% of all completeness predictions being within 1% of each other, and 91% being within 5% of each other **(Figure 6a)**. Similar congruency in results was observed for contamination, with 82% of all genome predictions being within 1% of each other, and 99% being within 5% **(Figure 6b)**. Substantially higher/lower completeness predictions (> 5% difference) using CheckM2 often occurred across entire lineages (phylum through to genera), while discrepancies in contamination were typically restricted to specific genomes within lineages (i.e. were not systematic; **Figure 6; Supplementary Table 5; Supplementary Figures 2 and 3)**. CheckM2 was also able to identify previously undetected contamination in a number of MAGs and isolate genomes and may avoid some potential contamination overestimations by CheckM1 (**Supplementary Note 8**).

**Figure 6:**
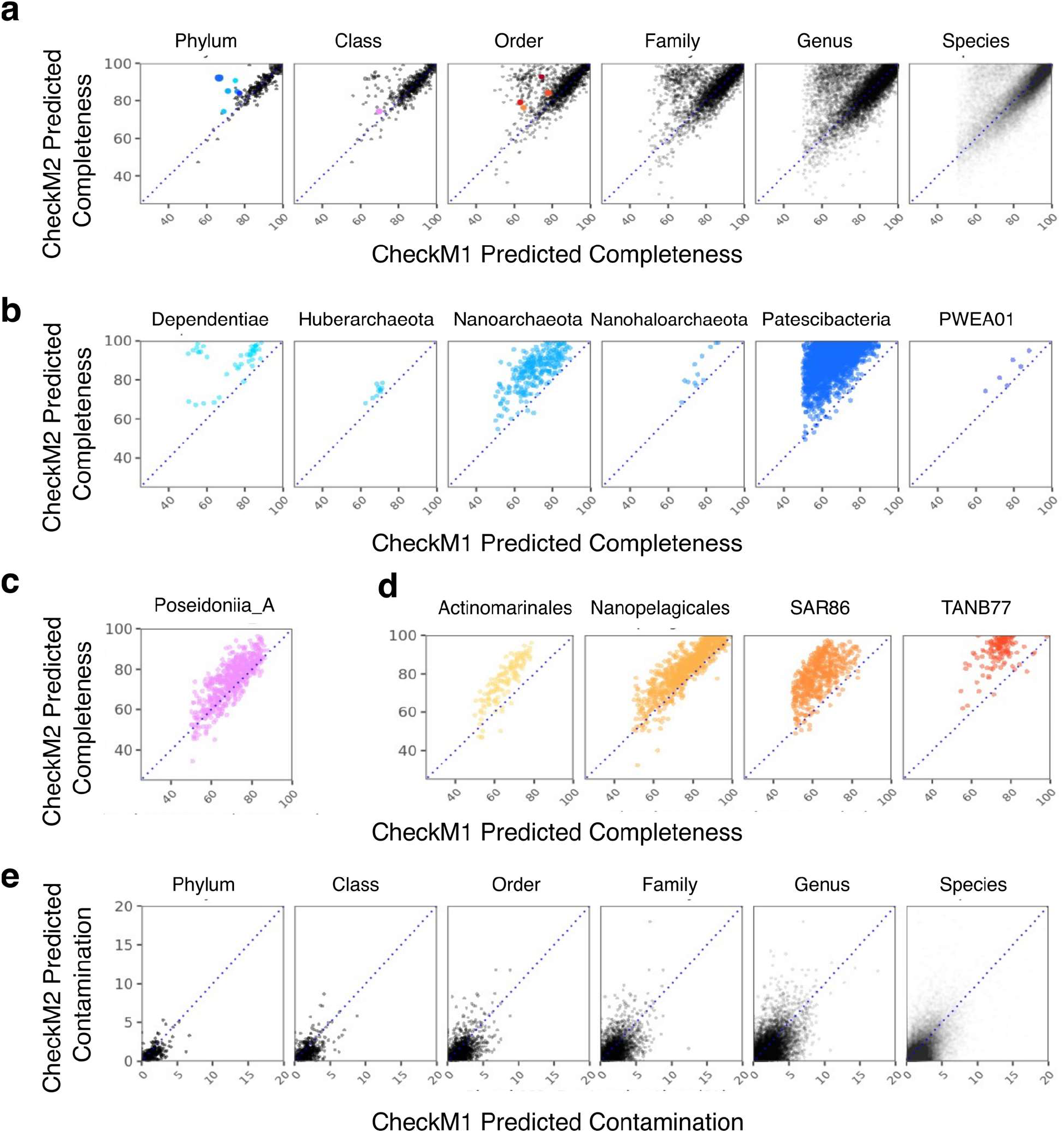
Genome quality predictions of CheckM1 versus CheckM2 for all GTDB r202 MAGs across different taxonomic levels. **a)** Mean completeness prediction from phylum to species, **e)** shows mean contamination prediction from phylum to species. For both, size of dots correspond linearly to number of genomes in taxa. **b)** shows completeness predictions for genomes in the specific phyla; **c)** completeness predictions for each genome in the class shown; and **d)** completeness predictions for each genome in the orders shown.

In the Bacteria, the highest divergence in completeness predictions is within the Patescibacteria phylum, where CheckM2 scores are substantially higher than those predicted by CheckM1 **(Figure 6c)**. Based on benchmarking, the CheckM2 results are likely to be significantly more accurate, enabling much better Patescibacteria MAG curation in the future and giving greater confidence to biological insights derived from these genomes. Other bacterial lineages predicted to be substantially more complete all appear to have common features such as small or reduced genome size, and/or hypothesized endosymbiotic or parasitic lifestyle. This includes the phyla Dependentiae, which are phylogenetically related to the Patescibacteria^22^, as well as the orders RF32 within the Proteobacteria, TANB77 within the Firmicutes_A^23^, and the actinobacterial orders Actinomarinales and Nanopelagicales^24^. Intriguingly, while some families within the Firmicutes_A order Christensenellales have concordant CheckM1 and CheckM2 predictions, other families such as CAG-74 have much higher CheckM2 completeness values. CAG-74 has been hypothesised to lack certain key functions (e.g. amino acid biosynthesis pathways) and may be potential symbionts^23^. Members of the family UBA1242, where the average genome size is ∼1 Mbp also shows substantially higher CheckM2 completeness predictions (on average 11% more complete), indicating that this family may also have a symbiotic or parasitic lifestyle which has not previously been reported (**Figure 6c**).

Analysis of manually curated complete bacterial endosymbiont genomes **(Supplementary Table 6)** demonstrated that CheckM2 markedly outperformed CheckM1 by predicting a much higher completeness, with CheckM2 predicting an average completeness of 71%, compared to CheckM1’s 39% average. Notably, CheckM2 was able to achieve this accuracy with little to no endosymbiont representation in its training database (as they are usually excluded from RefSeq) and incorporating the test genomes into the final models will likely substantially improve its accuracy on future endosymbiont cases. It is likely that use of CheckM2 on assembled metagenomic data will lead to the discovery of novel endosymbiont genomes that are highly complete with a small genome size.

Archaeal lineages with substantially higher CheckM2 completeness scores are primarily in the DPANN superphylum, including members of the Nanoarchaeota, Nanohaloarchaeota, and Micrarchaeota phyla, which have high-quality genomic representatives in CheckM2, as well as the phyla Huberarchaeota, Aenimatarchaeota and PWEA01 (formerly part of Aenigmatarchaeota), which are not represented in the CheckM2 release reference set **(Figure 6c)**. These predictions underscore the effectiveness of CheckM2’s prediction approach which generalizes to novel taxa with biological similarities to genomes CheckM2 was trained on. Other lineages that are predicted to be more complete by CheckM2 include the class Poseidoniia_A within the Thermoplasmatota, which is missing a number of single-copy genes used by CheckM1^25^, and the Asgardarchaeota order CR-4. The recently isolated and sequenced *Prometheoarchaeum syntrophicum*, which belongs to the order CR-4, was included in CheckM2’s release reference set, which likely contributed to a higher completeness score for this lineage. This highlights the power of including a single genome representative in CheckM2’s machine learning predictions.

In a small number of instances, CheckM2 completeness values were substantially lower than CheckM1 (5.4% of all genomes were ≥5% lower, 1.6% of genomes were ≥10% lower or more). The underlying cause for this difference is unclear but is likely due to multiple factors, such as the novelty of genomes, CheckM2’s choice of ML model, or CheckM1’s use of a kingdom-level marker set to assess the completeness of some novel lineages (**Supplementary Note 5**). Indel-dominated genomes were also found to have a particularly low CheckM2 score relative to CheckM1 **(Supplementary Note 6)**. Furthermore, as CheckM1 is often used to select MAGs for submission and publication, this can produce an imbalanced selection effect where genomes with overprediction error are retained in databases at higher rates than genomes with underprediction errors (**Supplementary Note 7**). Given the benchmarking results and careful investigation of example cases it is likely that CheckM2 is more accurate than CheckM1 in the majority of these instances. However, for some lineages with only a few MAGs and no complete genomes it is difficult to assess whether CheckM1 or CheckM2 scores are more accurate. The addition of any single complete representative genome will improve/validate the accuracy of CheckM2’s predictions for these lineages.

Overall, we see good congruence between CheckM2 and CheckM1, and increased CheckM2 completeness scores in lineages where CheckM1 is known to have poor predictive capacity^17,26^. This gives confidence in the robustness and reliability of both estimates, given the different underlying algorithms behind both tools. Based on these results, the benchmarking data sets and investigation of individual cases **(Supplementary Notes 3-8)** we strongly believe that in most cases of incongruent predictions, CheckM2 values are likely to be substantially more accurate.

#### Biological insights into the machine learning models

It is difficult to identify the contribution of specific genomic features to the predictions of machine learning models used by CheckM2. Some interpretable machine learning approaches, such as SHAP^27^, use robust mathematical techniques to approximate feature importance. While imperfect, when applied in the context of CheckM2’s models, these approaches can highlight the importance of specific genes/pathways that can be further investigated and assessed independently.

According to their SHAP values, key pathways contributing to completeness predictions across most lineages are ribosomal proteins, as well as genes in the DNA processing and tRNA biosynthesis pathways. There are also individual pathways with substantially higher predictive values for only certain lineages. For example, the membrane transporters pathway in the Patescibacteria have much higher importance values compared to most other lineages which are not characterized by genome reduction or streamlining **(Supplementary Table 7)**. Transporters are particularly noteworthy as they are likely to be key to an auxotrophic lifestyle, while also presenting an example of a set of genes that would be missed using only a conserved single-copy marker approach. The high importance placed on these pathways are in line with our biological understanding of microorganisms and give confidence that the machine learning models are capturing details of underlying biological reality.

### CheckM2 updates, computational benchmarking and resources

CheckM2 will be updated in line with GTDB releases. Unlike the very computationally heavy simulation method of CheckM1^4^, the simulation and training with new complete genomes takes less than 1 min per genome (per thread), with KEGG annotation of simulated genomes using DIAMOND forming the only computational bottleneck. This means that a new GTDB release can be updated into CheckM2 within 24-48 hours, as the speed of updating CheckM2 with new simulated genomes is generally equivalent to running CheckM2 on the simulated genomes.

During runtime, CheckM2 was consistently faster than CheckM1, processing an average of 1.56±0.83 genomes per minute per thread on an AMD EPYC 7702 64-Core Processor, relative to CheckM1’s 0.57±0.19 genomes per minute per thread. As CheckM2 has no taxonomic determination step, its speed is more variable than CheckM1, being substantially faster when predicting the quality of small or low completeness genomes. CheckM2 is capable of processing hundreds of thousands of genomes at a time with reasonable (< 90 GB for a batch run of 224,000 genomes) RAM usage.

## Discussion

Here we present a novel machine learning approach for predicting completeness and contamination of microbial genomes derived from metagenomic, single-cell and isolate sequence data. When benchmarked against CheckM1, we show congruency in genome quality prediction for lineages with good genomic representation, but demonstrate that CheckM2 has substantially better accuracy on medium and low-quality genomes and genomes from lineages with poor genomic representation. We also demonstrate that in the majority of cases it is able to generate highly accurate predictions for genomes in phyla with only a single genomic representative. Additionally, CheckM2 is substantially more accurate on lineages with small or reduced genomes such as the DPANN, Patescibacteria and Dependentiae, where CheckM1 often produces highly inaccurate predictions. Finally, CheckM2 typically performs better than or equal to CheckM1 on novel lineages with no genomic representation.

The use of genome quality predictions from CheckM2 are likely to have important implications for existing databases and biological interpretations of novel lineages. For example, CheckM2 completeness predictions will allow the inclusion of additional genomes currently excluded from GTDB due to the inaccurate CheckM1-based minimum cutoff (50% completeness), as demonstrated for the Patescibacteria phylum and DPANN.

Improved genome quality predictions by CheckM2 are the result of considering fully annotated genomes in its machine learning models, as opposed to CheckM1’s requirement for single-copy marker gene sets for each lineage. An additional advantage of the CheckM2 approach is that its models can be easily and rapidly updated to incorporate additional high quality genomic representation for novel lineages, further increasing the accuracy of its genome quality predictions. Future version of CheckM2 will be iteratively updated and may also include additional annotation databases (e.g. STRING^28^, EggNOG^29^) if this leads to significant improvements in genome quality predictions. CheckM2 is a major step forward in our ability to rapidly and accurately predict genome quality across the microbial tree of life.

## Methods

### Simulating genome completeness and contamination

To construct a training and validation set of genomes with known ranges of completeness and contamination, all genomes from RefSeq release 89 annotated as ‘complete’ or ‘chromosome’ were downloaded and dereplicated at 99% ANI using the software package Galah^30^. Genes were predicted using Prodigal v.2.6.3^31^, and resulting predicted proteins were randomly sampled using the BBMap^32^ suite v.38.18 (‘reformat.sh’ with the ‘samplerate’ option) to create a range of completeness between 5% to 100% at 5% intervals. To simulate self-contamination, the same proteins were sampled multiple times to generate a range of 0% to 35% contamination. True completeness was calculated as 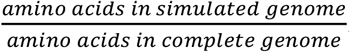. True contamination % was calculated as 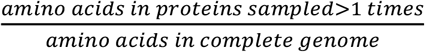.

For testing, two different simulation methods were used. These were purposefully distinct from the training set to minimize any overfitting to the simulation method. In the ‘20-kb-fragmentation’ method, the input genome was fragmented into approximately 20-kb fragments using the ‘shred.sh’ option of the BBMap suite with median length 20,000 and variance 2000. These were sampled using the BBMap suite (‘reformat.sh’ with the ‘samplereadstarget’ option) to create a range of completeness and contamination values as described above. The fragments were not stitched back together to replicate the state of a MAG with multiple contigs. In the ‘MAG-derived-length-fragmentation’, a database of contig size distributions was created using GTDB release 95 as its base, where the contig size distribution of all MAGs with less than 350 and more than 10 contigs was used as a pool to choose from. Each genome had to be at least 65% complete and no more than 5% contaminated as determined using CheckM1. For each test genome, a MAG contig distribution was randomly selected and the full genomic sequence of the genome being simulated was cut into the same length fragments relative to genome size, where the order of contig sizes was randomized. These were then sampled in the same way as in the ‘20-kb-fragmentation’ method. For both, true completeness was calculated as 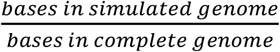. True contamination % was calculated as 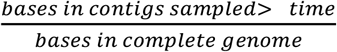.

### Annotation of genomes

For all simulated genomes, genes were predicted using Prodigal v.2.6.3 and annotated with KEGG^33^ ids using diamond v2.0.4 ‘blastp’ command against uniref100 (released 26/11/2018) containing KO annotations. To filter annotations, a query_cover of 80, a subject_cover of 80, an evalue of 1e-05, and a percent_id of 30 was used, taking only the top hit for each gene. Annotations were converted to a frequency matrix containing all existing KEGG ID’s with rows representing a simulated genome and columns representing counts of annotation detected. KO’s found in the same pathway were grouped next to each other in order to allow sliding convolutional windows of the neural network to extract useful information from this grouping. KO’s present in multiple pathways were assigned only to the first pathway based on pathway alphabetical order.

After annotation, KEGG pathway, module and category completeness was calculated based on the definitions for each downloaded from KEGG on 26-11-2018 where completeness was defined as the fraction of genes present in a genome out of all genes defined in a module, pathway or category. Each module, pathway and category completeness feature vector was encoded as an additional column with fractional value between 0 and 1. Nested modules (modules containing other modules) were not used. Only the ‘general’ gradient boost model utilized these additional completeness feature vectors.

Additionally, the frequency counts of each amino acid, number of coding sequences and the total amino acid length of each genome was calculated and added to the protein annotations.

For testing, all genes were predicted and annotated de novo for each simulated genome.

### Selection of additional genomes

To widen the scope of the trained model and reduce the uniformity of the training data set, a small number of potentially complete non-RefSeq genomes were identified using a ‘repeated-quality-metrics’ strategy: CheckM1 and CheckM2 (trained only on RefSeq release 89) were used to assess all GenBank genomes part of release 89. Those that had at least 5 or more members of a species, and had the same completeness and contamination scores from CheckM1 for more than three quarters of them as well as a genome size within 5% of each other were selected as potentially complete. A genome of the same length assembled repeatedly to yield the same completeness and contamination statistics was used as potential evidence for completeness. As these genomes were often in multiple contigs, the contigs were sampled randomly and completeness calculated based as 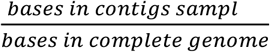. From these generated synthetic genomes, only those with less than 85% or less completeness were used for training in order to avoid incorrect bias for highly-complete genomes in CheckM2. These new genomes were added to the training pool with a 50% lower sample weight relative to known complete RefSeq genomes. Even if some of these genomes are not complete, the addition of potential noise was also a desirable part of this process as a potential regularisation constraint on the neural network model. Moreover, these genomes also introduce a more accurate example of low-completeness genomes compared to genomes generated using the ‘random-protein-sampling’ method used for the bulk of the training set, further increasing the neural network model’s accuracy in these cases. These genomes (**Supplementary Table 9**) were added to the training pool used to train neural network models but not to the gradient boost models (completeness or contamination). Neural network models trained with these additional genomes included those used to benchmark RefSeq release 202, benchmark the novel circular MAGs from Singleton et al., and Lui et al., as well as used in the final release of CheckM2.

### Training machine learning models

To train the ‘general’ gradient boost models, annotations of genomes in the training set were used as feature vectors, with contamination and completeness values being the predictor targets for the ‘general’ completeness and ‘general’ contamination models. The lightgbm^14^ package was used to train a regression model with the following parameters: ‘boosting type’: ‘gbdt’, ‘objective’: regression, ‘num_leaves’: 11, ‘min_data_in_leaf’: 150, ‘learning_rate’: 0.2, ‘feature_fraction’: 0.5, ‘bagging_fraction’: 0.5, ‘baging_freq’: 3, ‘reg_sqrt’: True, ‘min_child_weight’: 180 for completeness and ‘boosting type’: ‘gbdt’, ‘objective’: regression, ‘num_leaves’: 211, ‘learning_rate’: 0.2, ‘feature_fraction’: 0.9, ‘bagging_fraction’: 0.8, ‘baging_freq’: 5, ‘reg_sqrt’: True for contamination. Both models were boosted for 450 iterations.

To train the ‘specific’ neural network model, tensorflow^13^ v2.2.0 was used. The model architecture was encoded using the keras API and consists of a sequential model with three 1D convolutional layers (kernel_size=10, strides=10, activation=‘relu’) with size 180, followed by another with size 100. These are flattened and followed by a dense layer (size=100, activation=‘relu) connected to an output layer (activation ‘sigmoid’ for the completeness model and ‘linear’ for the contamination model). A BatchNormalisation layer was added after each convolutional layer to standardize and normalize network weights and feature vector input. The loss_weight parameter was used at compilation and completeness of samples * 500 was passed as weight to penalize the model errors more for more complete genomes (thus aiming for higher accuracy on higher-quality genomes). The addition of the batch normalisation layers added substantial improvement in accuracy, but also caused validation loss to fluctuate substantially during training. Therefore, the model was trained with checkpoints on complete RefSeq genomes simulated using ‘random-protein-sampling’ as well as non-RefSeq genomes identified using the ‘repeated quality metrics’ strategy outlined below, for 5-15 iterations, while using validation accuracy on a subset of the training set (high, medium and low-quality simulated MAGs, see **Supplementary Table 1**) to identify and select the model iteration with best validation loss across all quality levels. This process was repeated for both the validation and final CheckM2 neural network models. For more details, see **Supplementary Note 1**.

### Filtering out low-quality genomes

While the majority of genomes in RefSeq are likely to be complete if annotated as such, there will always be exceptions. To remove potentially incomplete genomes, an intermediate neural network model was trained for 2 epochs on all training genomes, then used to predict their completeness and contamination. Those with high deviations (more than 10% difference) between predicted and assumed completeness or contamination were removed from the training set. Notably, most of these genomes also had inferior CheckM1 metrics. Generally they were found to belong to species which had at least one other complete genomic representative in RefSeq which did not show high deviation, indicating an issue with the genomes and not with the biology of particular lineages. A complete list is included in **Supplementary Table 10**.

### Benchmarking software

To benchmark CheckM2, CheckM1 were used as key tools of comparison. CheckM1 v 1.0.12 was run with the flag ‘lineage_wf’ for automatic lineage selection. Error in prediction of completeness or contamination was defined *predicted value – true value*, with negative errors representing under-prediction and positive errors representing overestimation of the true value. MIMAG completeness and contamination standards^12^ were used to divide all benchmarking results (**Supplementary Table 1**). MIMAG standards are often used to report the quality of MAGs and make for a useful way to visualize CheckM2 performance across varying genome quality. As CheckM2 does not detect nor rely on other MIMAG factors such as full-length 16S RNA or the presence of tRNAs, only the completeness and contamination information was used. For supplementary analysis, BUSCO^5^ predictions were added. BUSCO v5.0.0 was run offline with reference database downloaded on 21-02-21 with the option ‘--auto-lineage-prok --mode genome’. Results from BUSCO were parsed to select the specific output if it was generated, otherwise use the generic output file. In BUSCO’s output ‘C’ was used as completeness, while ‘D’ was used as contamination for benchmarking against CheckM1 and CheckM2.

### Benchmarking calculations

For all benchmarking on synthetic genomes, completeness error (defined as predicted completeness% - actual completeness %) and contamination error (defined as predicted contamination % - actual contamination %) was calculated for each genome, and the results graphed using a letter value plot in the Seaborn^34^ package. Mean absolute error values were calculated as 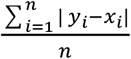 where y is the predicted value and x the true value. Phylum-wide mean absolute error value was calculated as 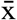 across 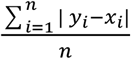 for each phylum where y is the predicted completeness value and x the true completeness value.

### Evaluation of taxonomic novelty effects on accuracy

To test the accuracy of both models on taxonomically novel groups, separate models were trained where one phylum was left out, the models were trained on all remaining phyla, and the models were then tested on the omitted group to determine error when predicting completeness and contamination. This was repeated for all phyla, then repeated for all subsequent taxonomic levels of relatedness (class, order, family, genus, species) with the left-out group doubling in scope every level (1 group left out per phylum, 2 per class, 4 per order, etc.) in order to account for increasing diversity and number of iterations necessary to cover all levels. Lineages representing multiple taxonomic levels of novelty (e.g. one class only containing one family) were tested only at the level of highest taxonomic difference (e.g. in this case at the class level). In all cases, GTDB r89 taxonomy was used to determine leave-out groups.

### Cosine similarity measure and model selection

In order to determine the appropriate model to use, CheckM2 uses the cosine similarity calculation 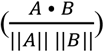 (where A and B are vector arrays) as cosine similarity correlates well with taxonomic novelty of query genome relative to the closest genome in the reference dataset. Rough taxonomic similarity enables selection between the general and specific completeness prediction models without the need to compute taxonomy, which would require substantially more computational time and resources.

As cosine similarity declines with completeness **(Supplementary Figure 1)**, a stable ‘novelty ratio’ was calculated by dividing completeness predicted by the general model by the squared cosine similarity, and subsequently used for selection between the neural network and gradient boost models.

Input for cosine similarity calculations is identical to the input into the neural network model. Based on the results from novelty testing, the median cosine similarity for novel phyla, classes orders and some families were assigned to be predicted with the ‘general’ model and all other with ‘specific’ model, including all genomes with mean completeness prediction below 50% (see **Supplementary Figure 1**). As completeness declines, a slightly higher share novel genomes is assigned to the neural network model to take advantage of its superior performance at lower completeness levels (see **Supplementary Table 4**).

### Evaluation of non-redundant contamination effects on accuracy

To simulate non-redundant contamination, simulated genomes were created from RefSeq release 89. Different levels of completeness were generated by removing a random subsection of the genome to generate completeness between 50% and 100%. A contaminant fragment was chosen from a taxonomic source defined by GTDB Release 89 taxonomic assignment. A randomly chosen isolate genome was chosen from the same species, same genus, etc. up to same domain, and a random subsection of the genome was used as the contaminating contig. The contaminating fraction did not exceed a maximum of 10% contamination, where contamination was calculated as 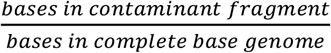. For each synthetic cross-contaminated species, only one other species was used, after which the same species could not be used as a source for contamination at that taxonomic level.

### Benchmarking speed of CheckM1 and CheckM2

Five replicate metagenomic sets were used to benchmark the speed of CheckM1 and CheckM2, consisting of an average of 450 genomes per set. Three sets consisted of MAGs randomly sampled from publicly available metagenomes, one set comprised a random selection of RefSeq r89 genomes, and one set comprised a random sample of synthetic MAGs belonging to the DPANN superphylum or Patescibacteria phylum derived from MAGs in RefSeq r202. CheckM1 was run in the ‘lineage_wf’ mode with 45 threads and 45 pplacer_threads, while CheckM2 was run in the ‘predict’ mode with 45 threads. All benchmarking was done on an AMD EPYC 7702 64-Core Processor, and time determined using the ‘time’ bash command, where the ‘real’ time was for comparison. During runtime, threads were not shared with any other processes. Time taken per minute per thread was calculated as 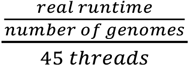. Peak RAM usage for a large batch job (225,000 GTDB Release 202 genomes in a single folder) was determined from ‘maximum resident set size’ using the command */usr/bin/time -v* and visual verification using the *htop* command.

### SHAP value calculations

SHAP values for the gradient boost models were calculated using the SHAP package (0.39.0). To calculate the ten feature vectors contributing most towards completeness predictions by phylum (**Supplementary Table 7**), a TreeExplainer was used to generate SHAP values for the gradient boost completeness model in the CheckM2 release version. Aggregate sums across all 21241 feature vectors were calculated, and the top ten were included in the Supplementary Table with a mean per phylum.

## Code availability

CheckM2 is available on GitHub (https://github.com/chklovski/CheckM2) and is released under the GNU General Public License Version 3.

## Acknowledgements

The authors thank E. McMaster for her help in refining the Figures. This work was supported by the National Science Foundation (NSF) Biology Integration Institute - EMERGE (DBI#2022070). A.C. is supported by Australian Government Research Training Program (RTP) Scholarships. G.W.T. is supported by Australian Research Council (ARC; Grants FT170100070). B.J.W. is supported by Australian Research Council Discovery Early Career Research (DE160100248).

## Author contributions

A.C. and G.W.T. designed the overall workflow and planned the key steps. A.C. generated the synthetic genomes, trained machine learning models, performed benchmarking and wrote the final code base of CheckM2. B.J.W. and D.H.P. guided code improvements and optimisations. G.W.T., D.H.P and B.J.W. helped interpret the result and made further suggestions for future directions and improvements. A.C. and G.W.T. drafted and wrote the manuscript. All authors edited the manuscript before submission.

## Footnotes

1 The most recent available at the time; GTDB r207 was released while manuscript was in preparation.

