## Supplementary Figures 1-3 for "CheckM2: a rapid, scalable and accurate tool for assessing microbial genome quality using machine learning"

**CheckM2 Supplementary Figures**

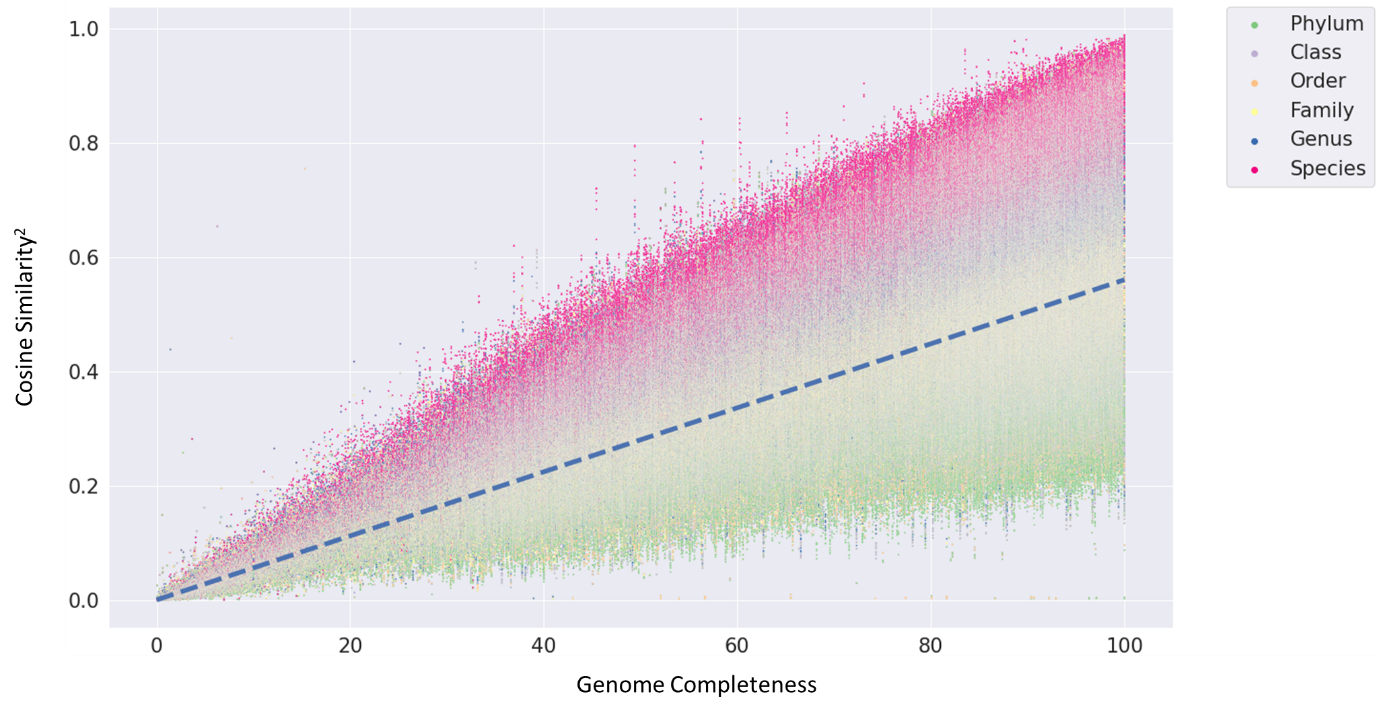

**Supplementary Figure 1**: Cosine similarity versus genome completeness. Legend/colours indicate taxonomic similarity of genome relative to reference, while regression line represents the switch from using the gradient boost model (below the line) to the neural network model (above the line).

**a)**

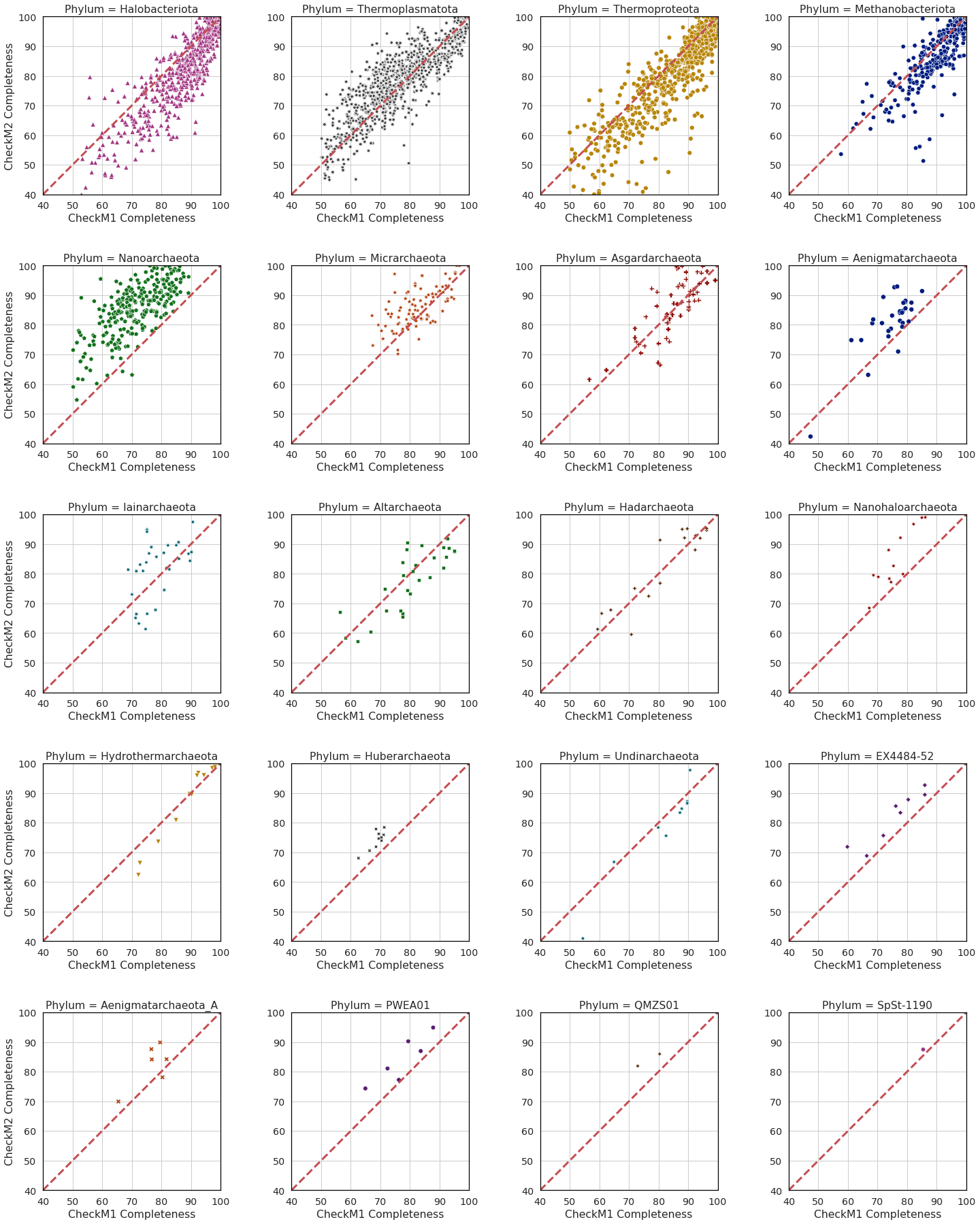

**b)**

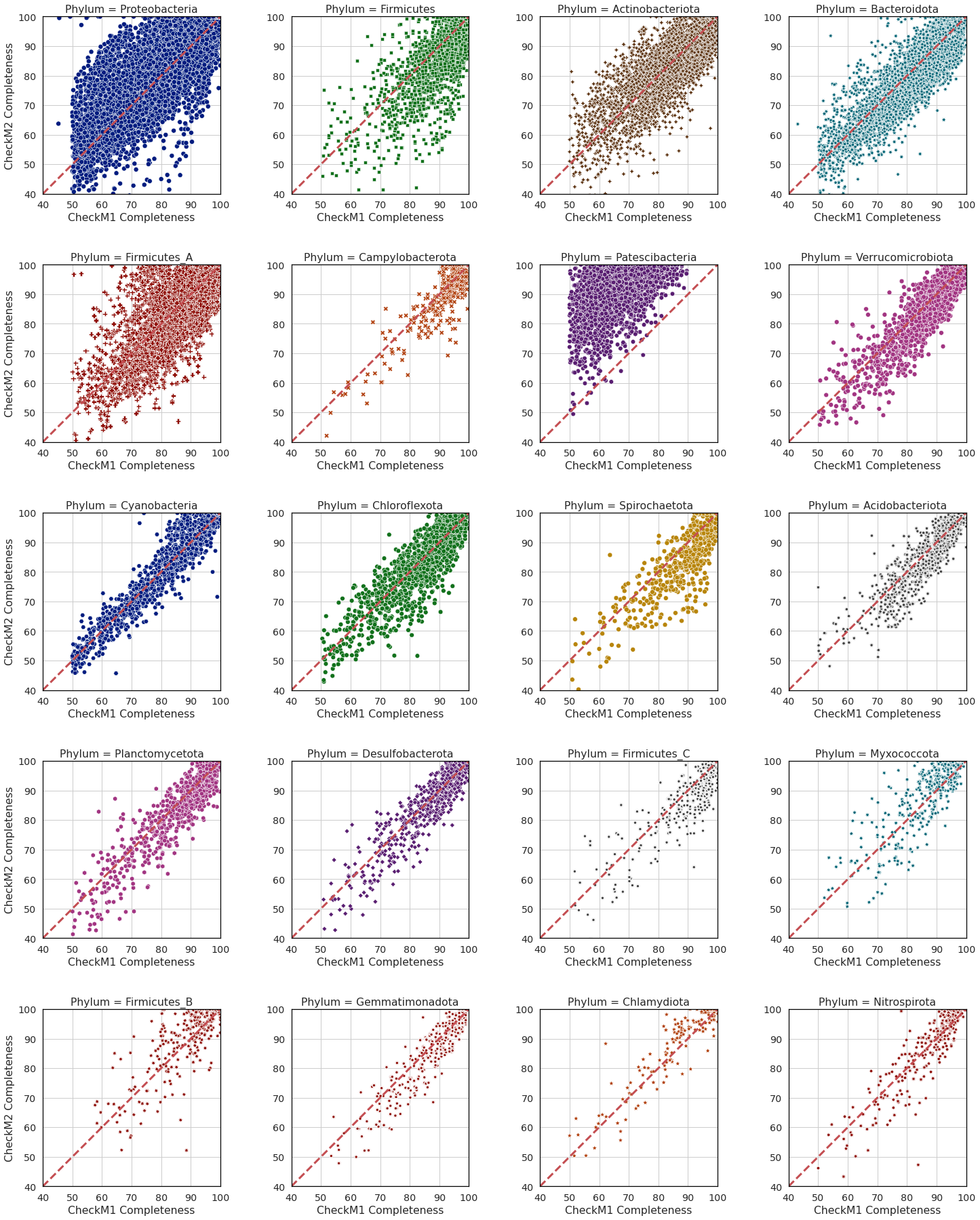

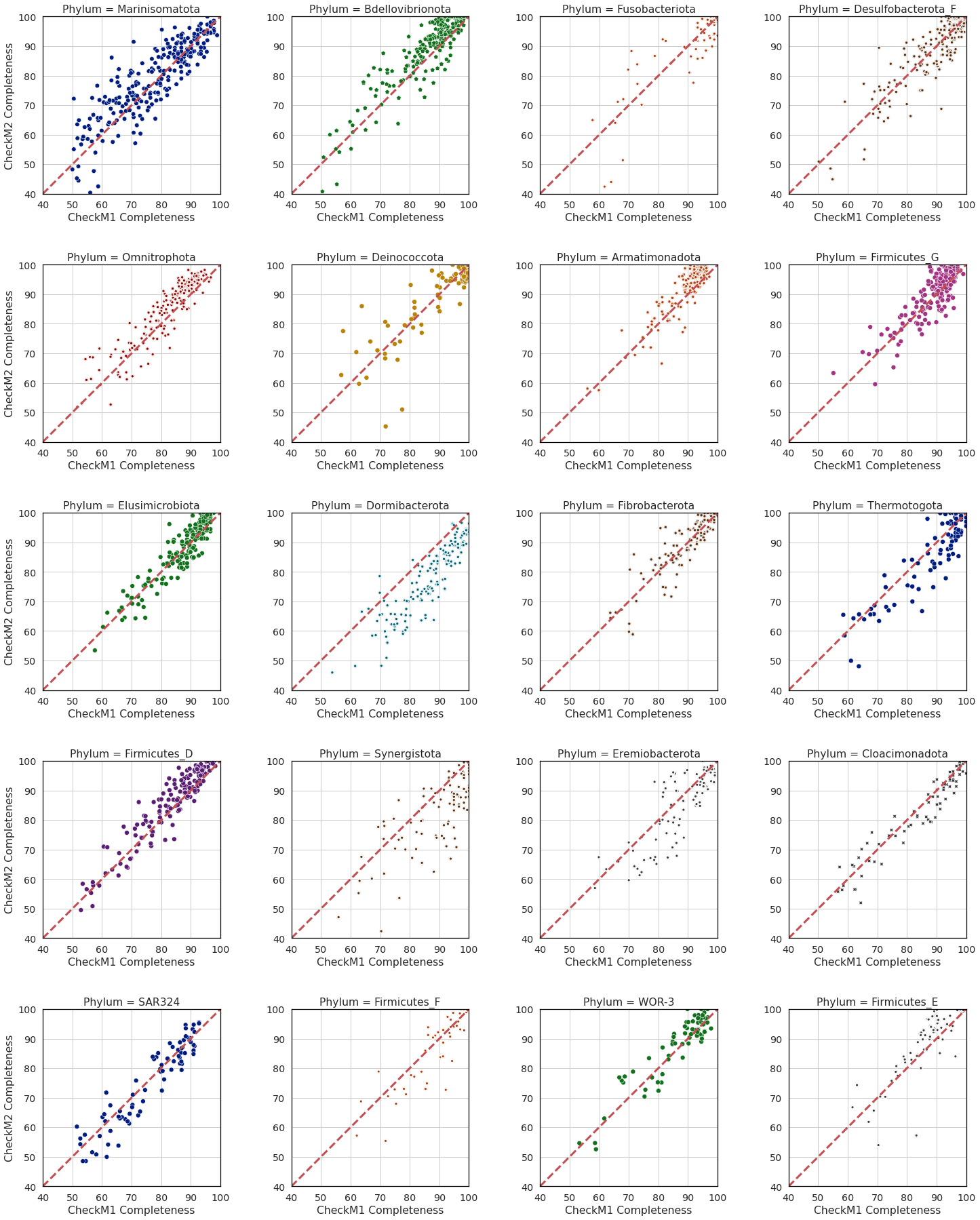

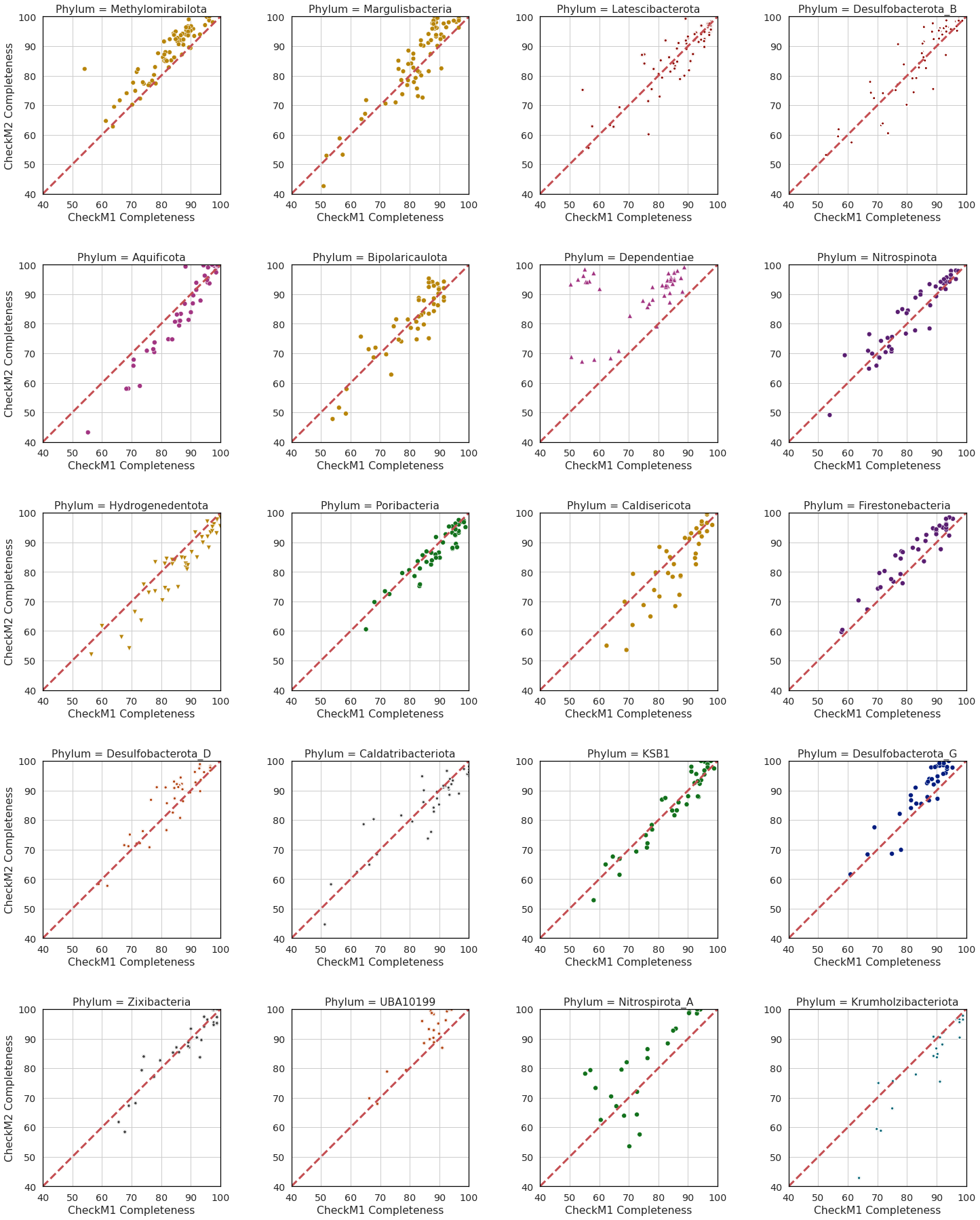

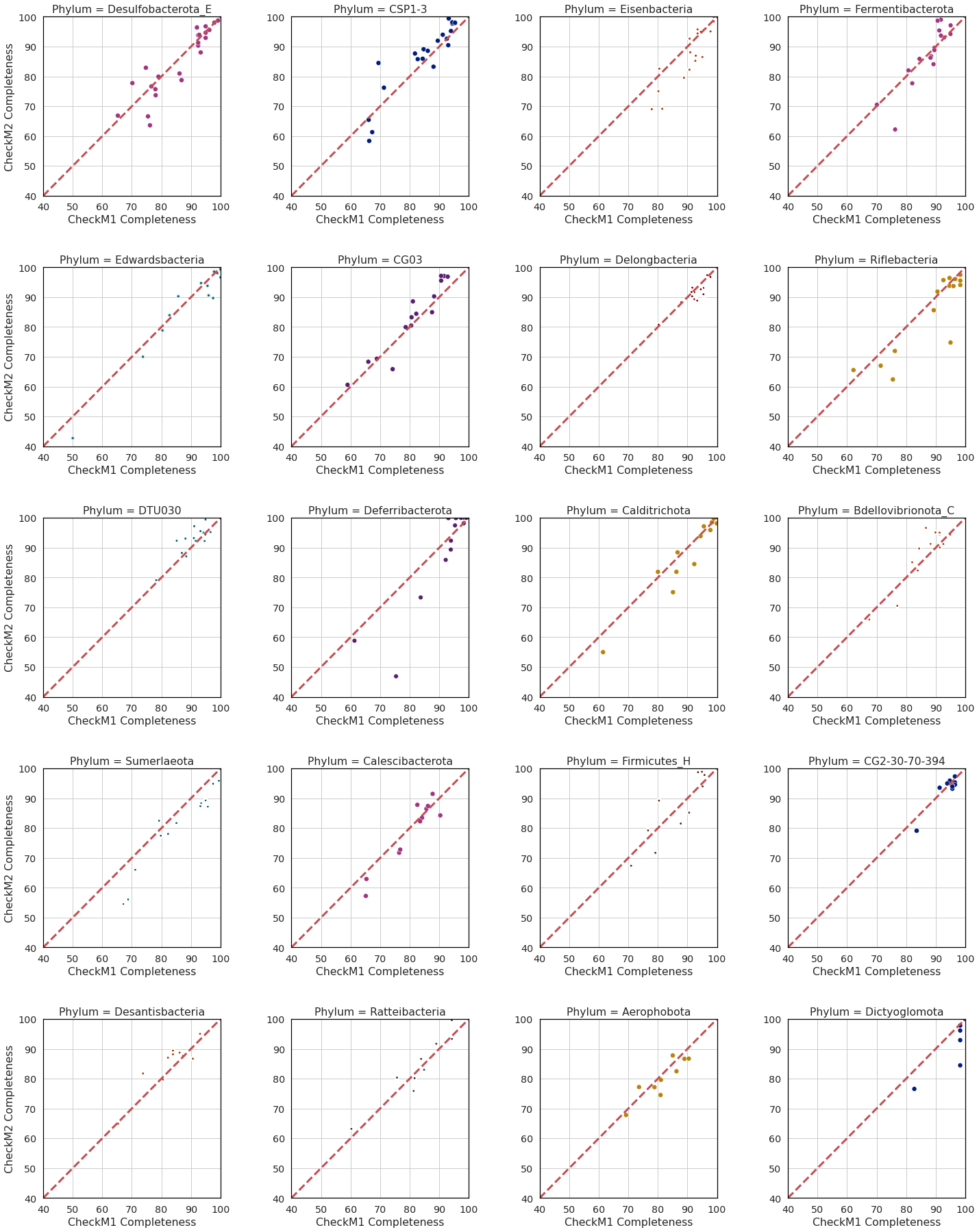

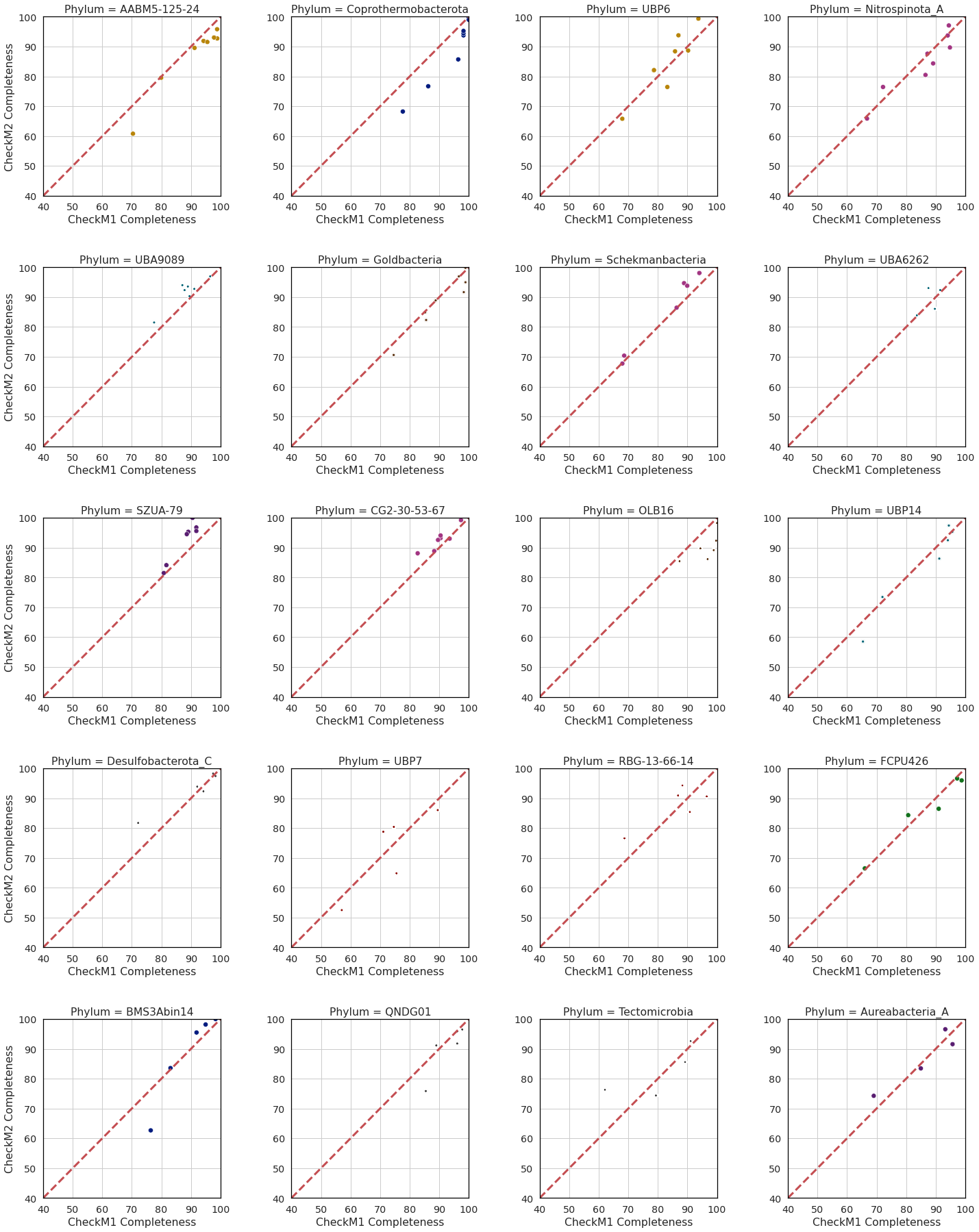

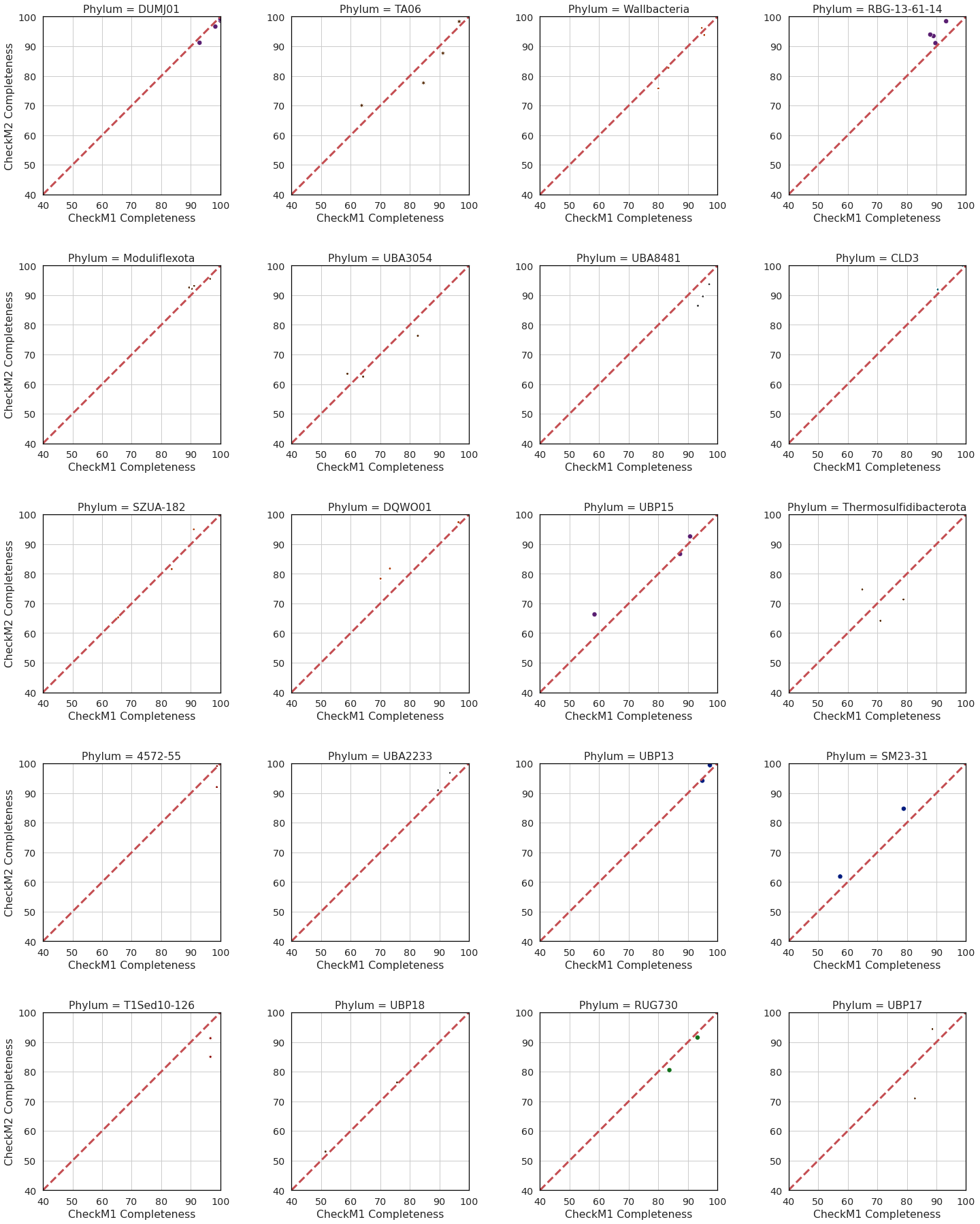

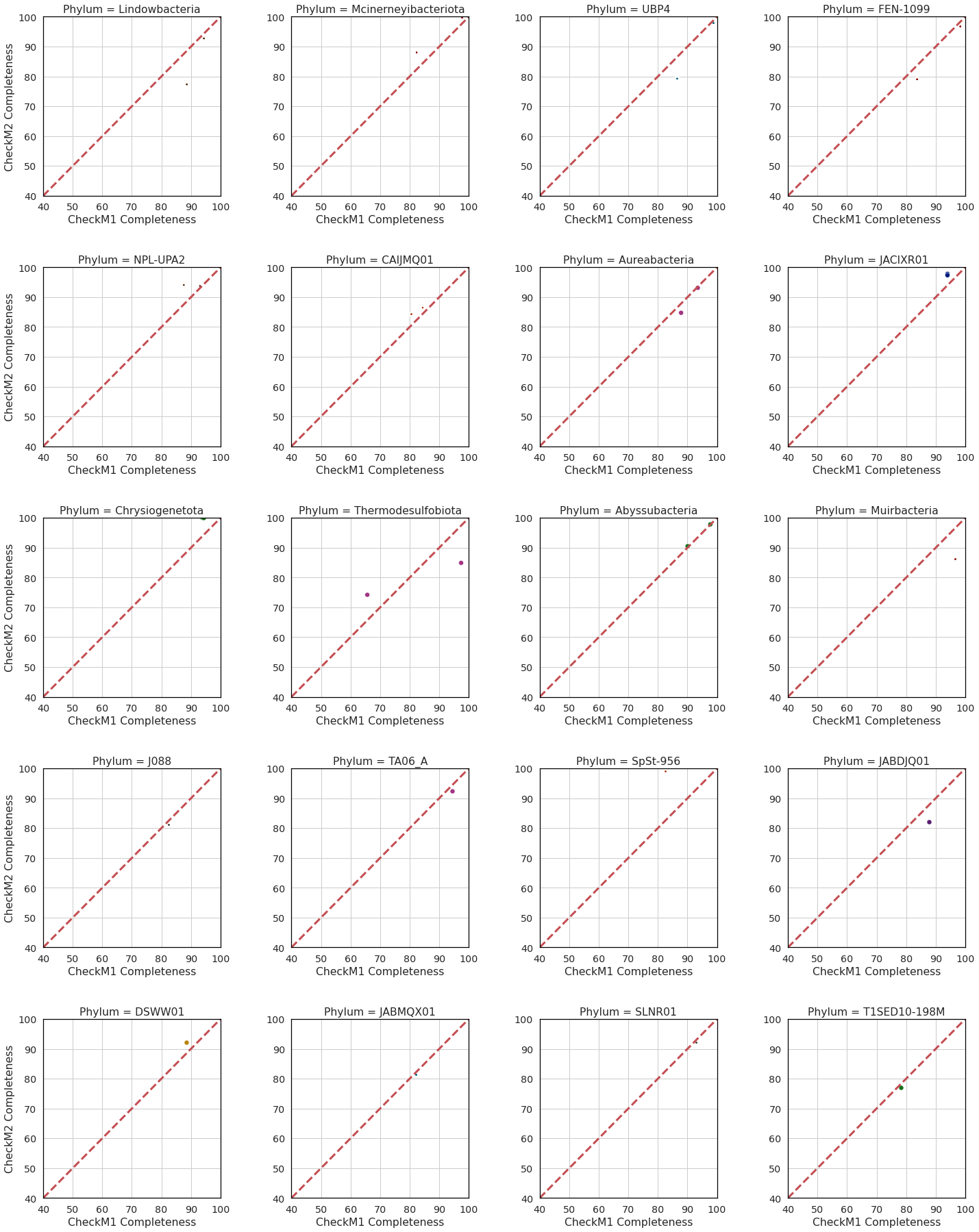

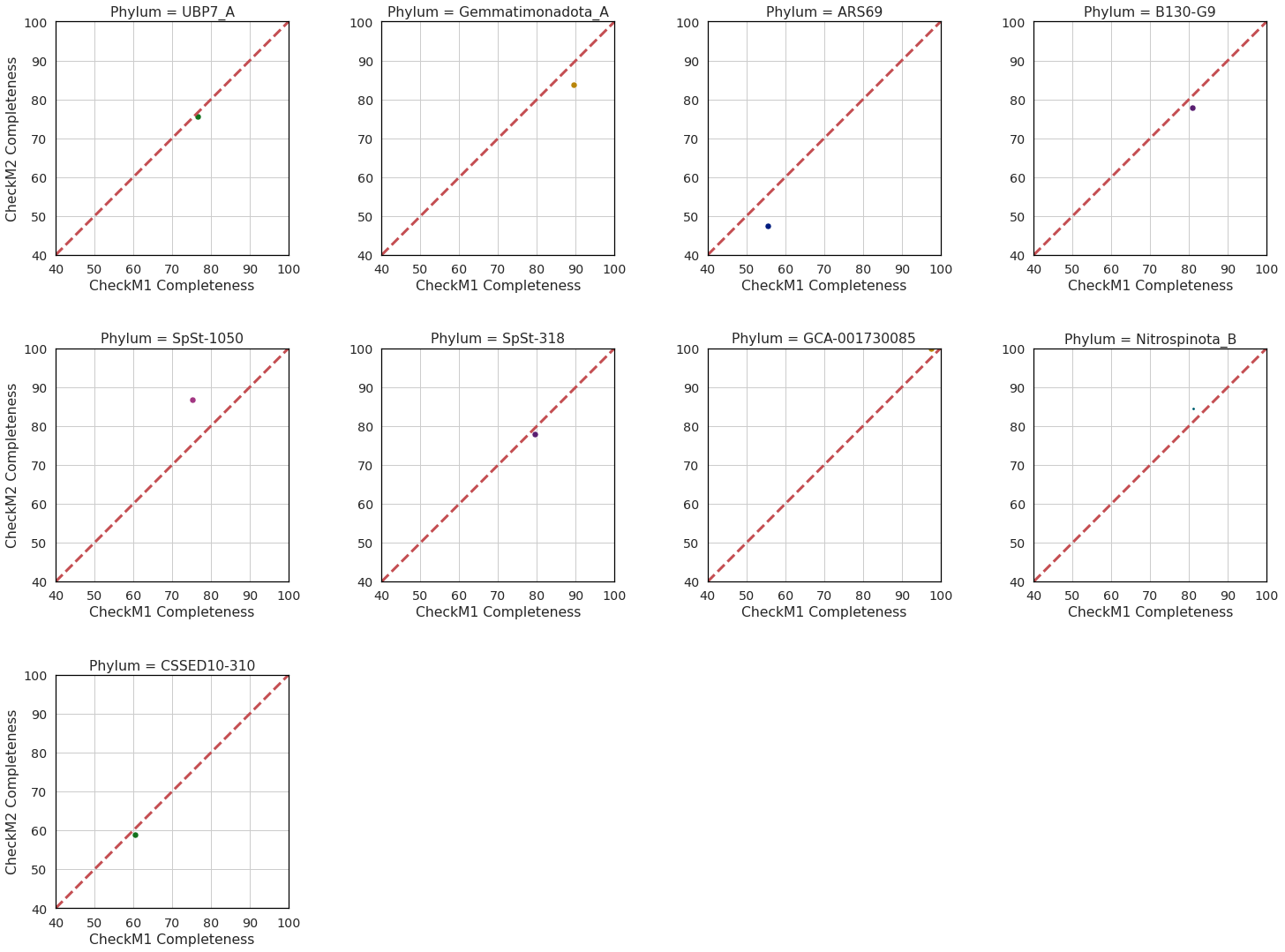

**Supplementary Figure 2**: CheckM2 vs CheckM1 completeness prediction by phylum for **a)** Archaea and **b)** Bacteria MAGs from GTDB release 202.

**a)**

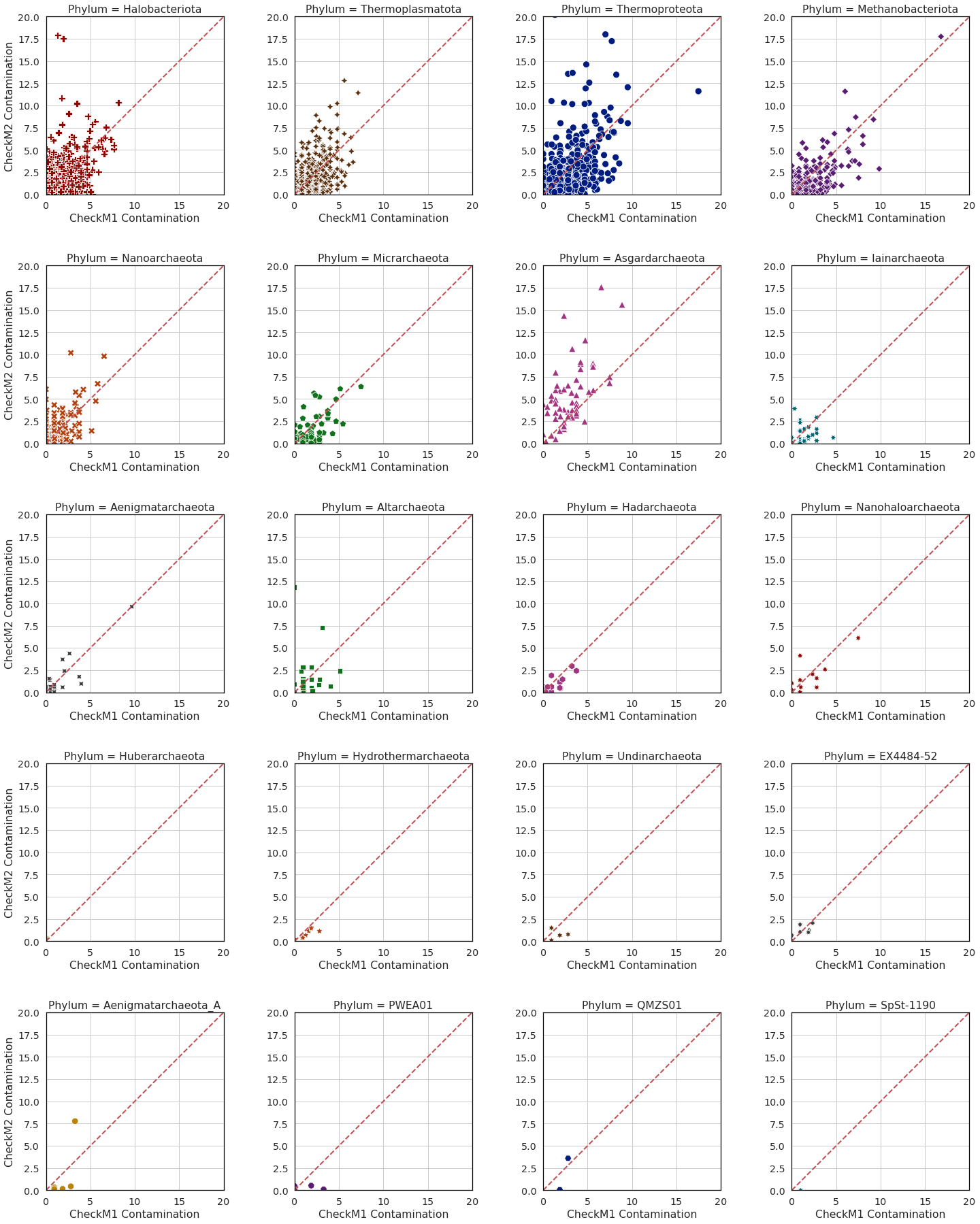

**b)**

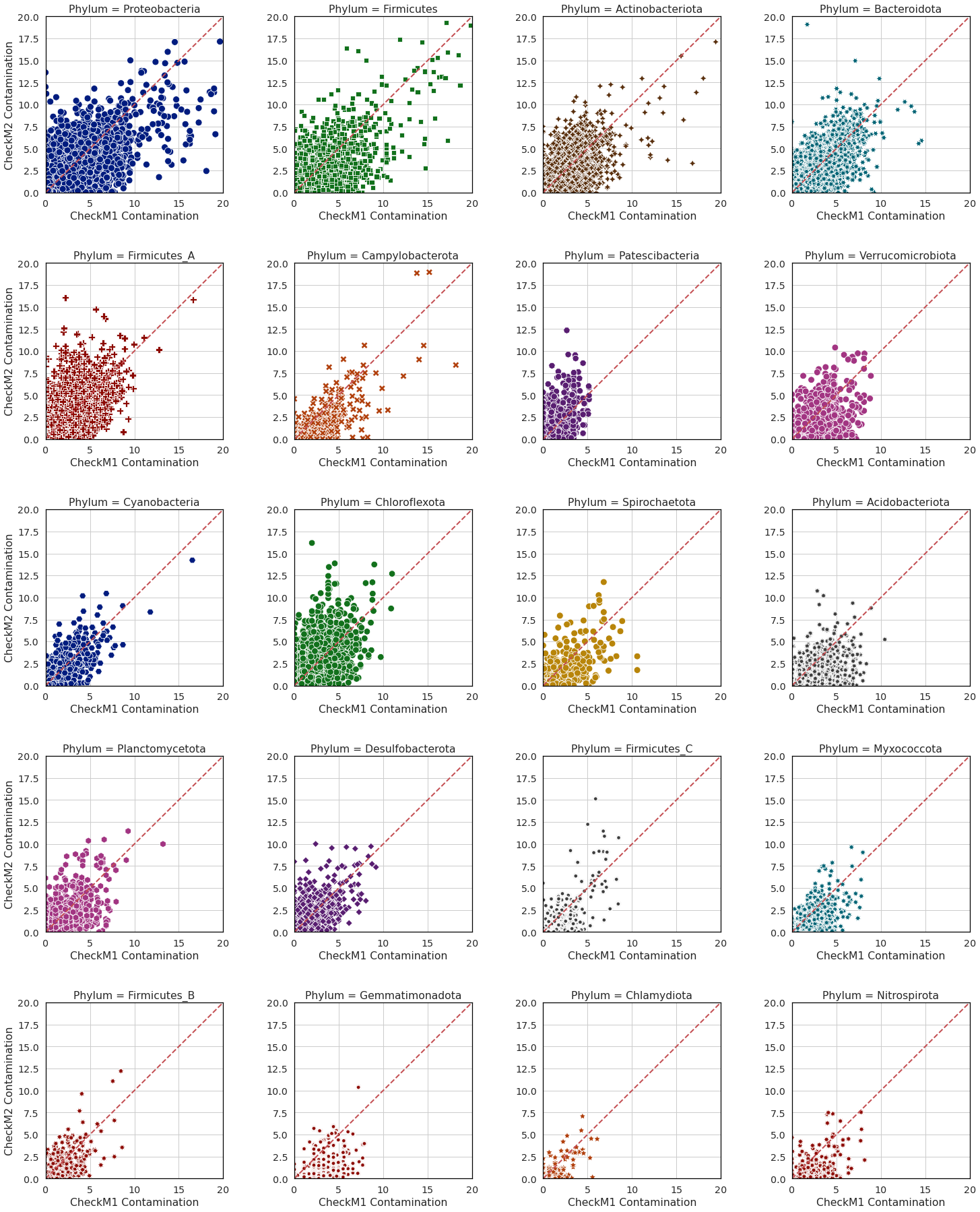

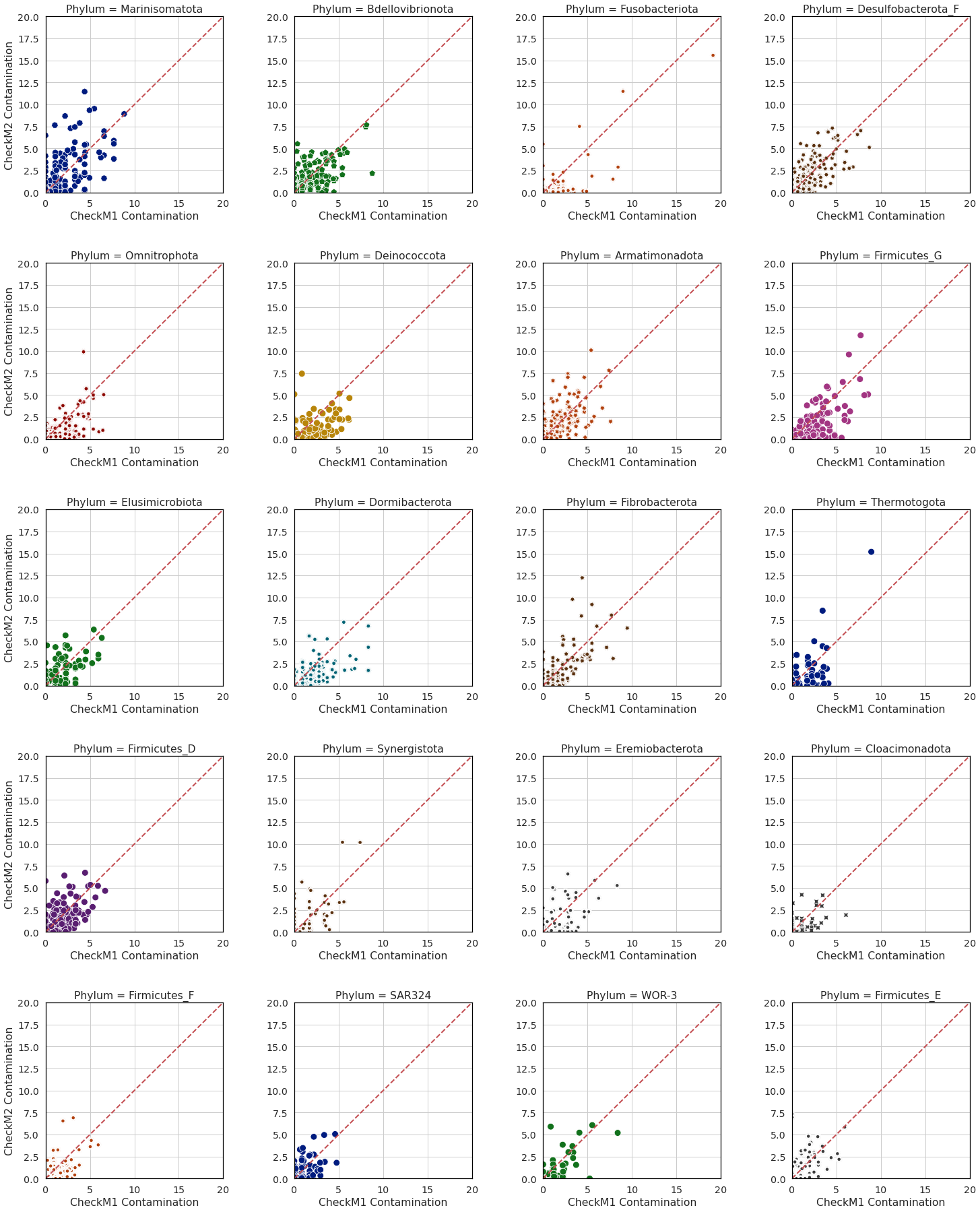

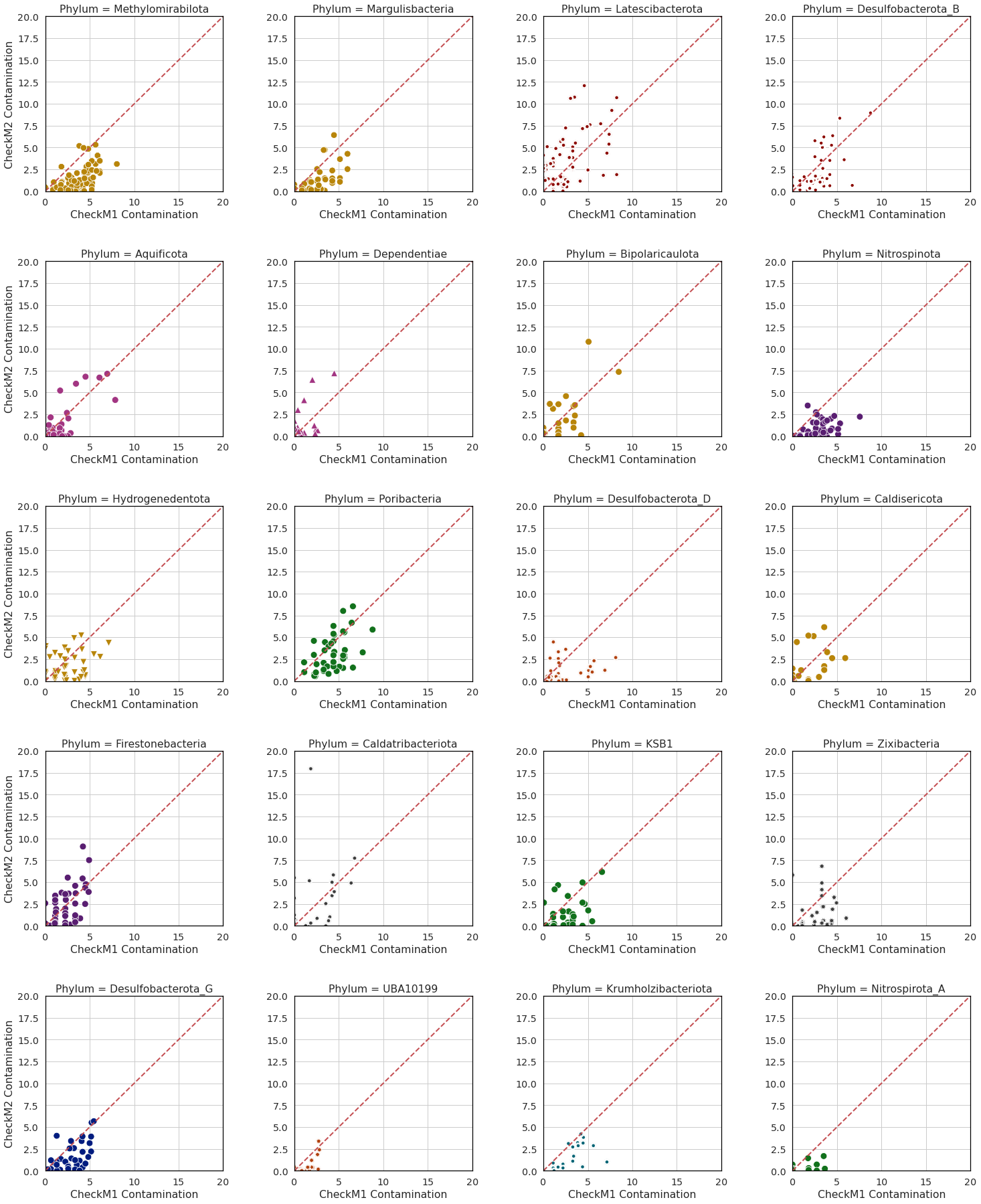

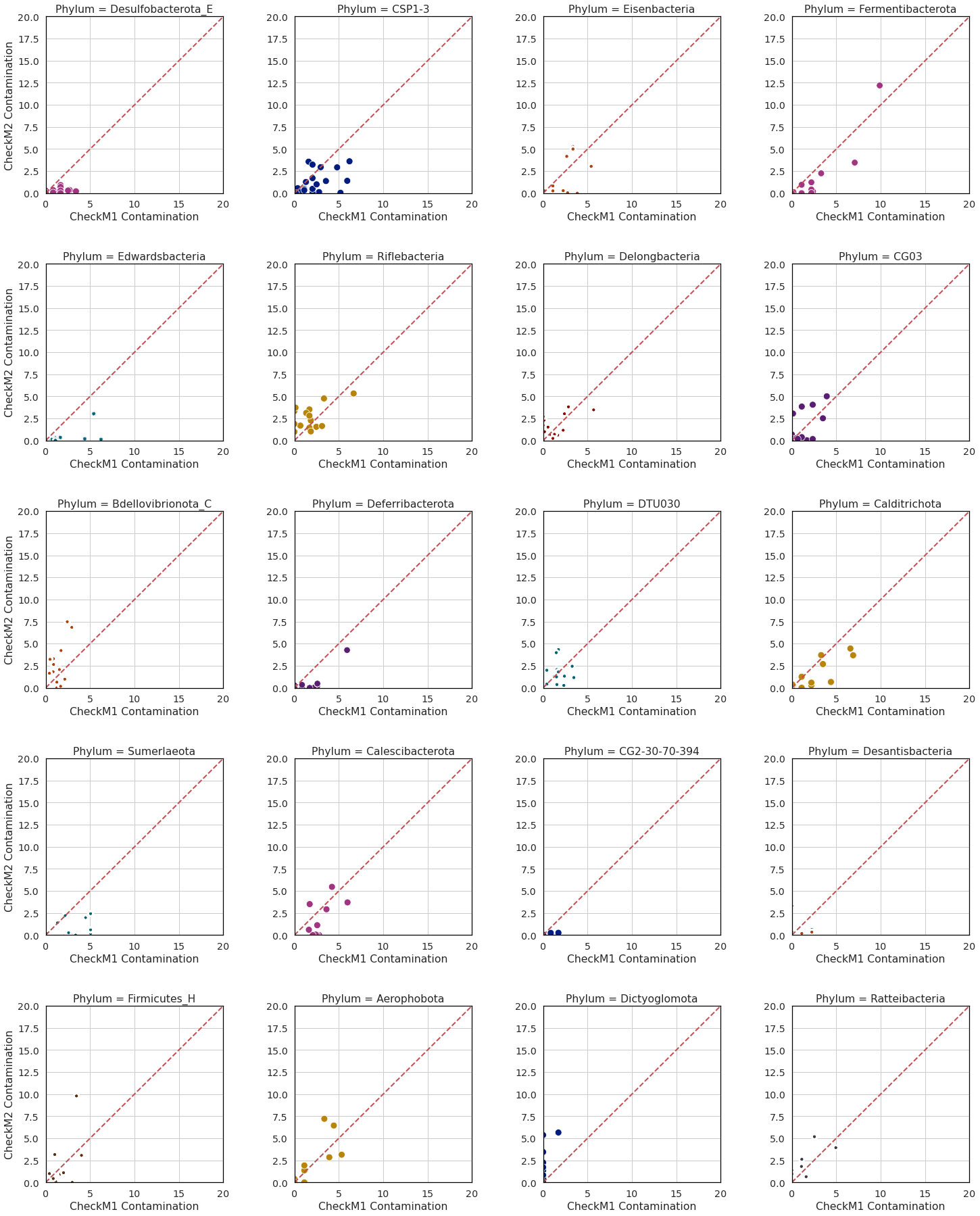

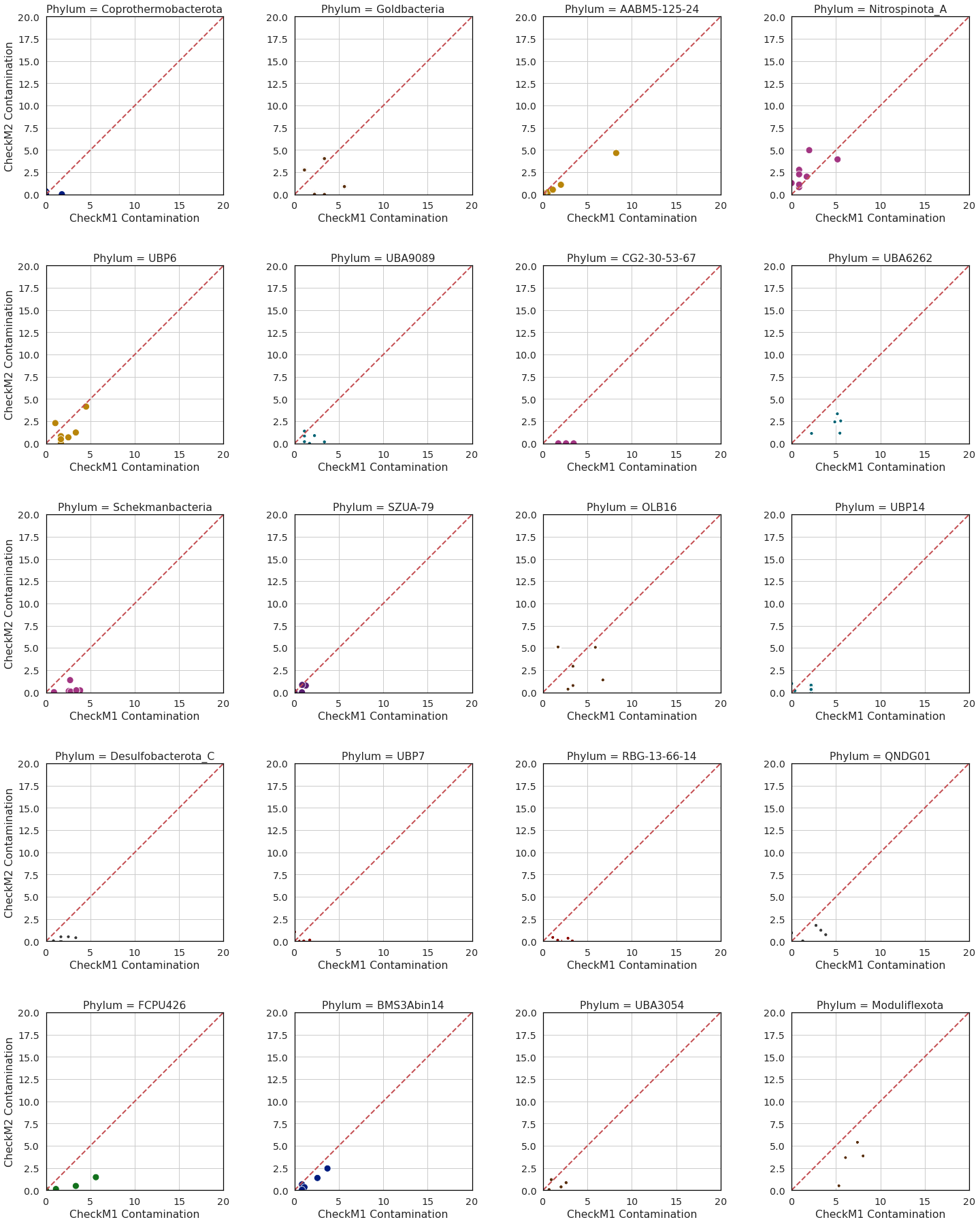

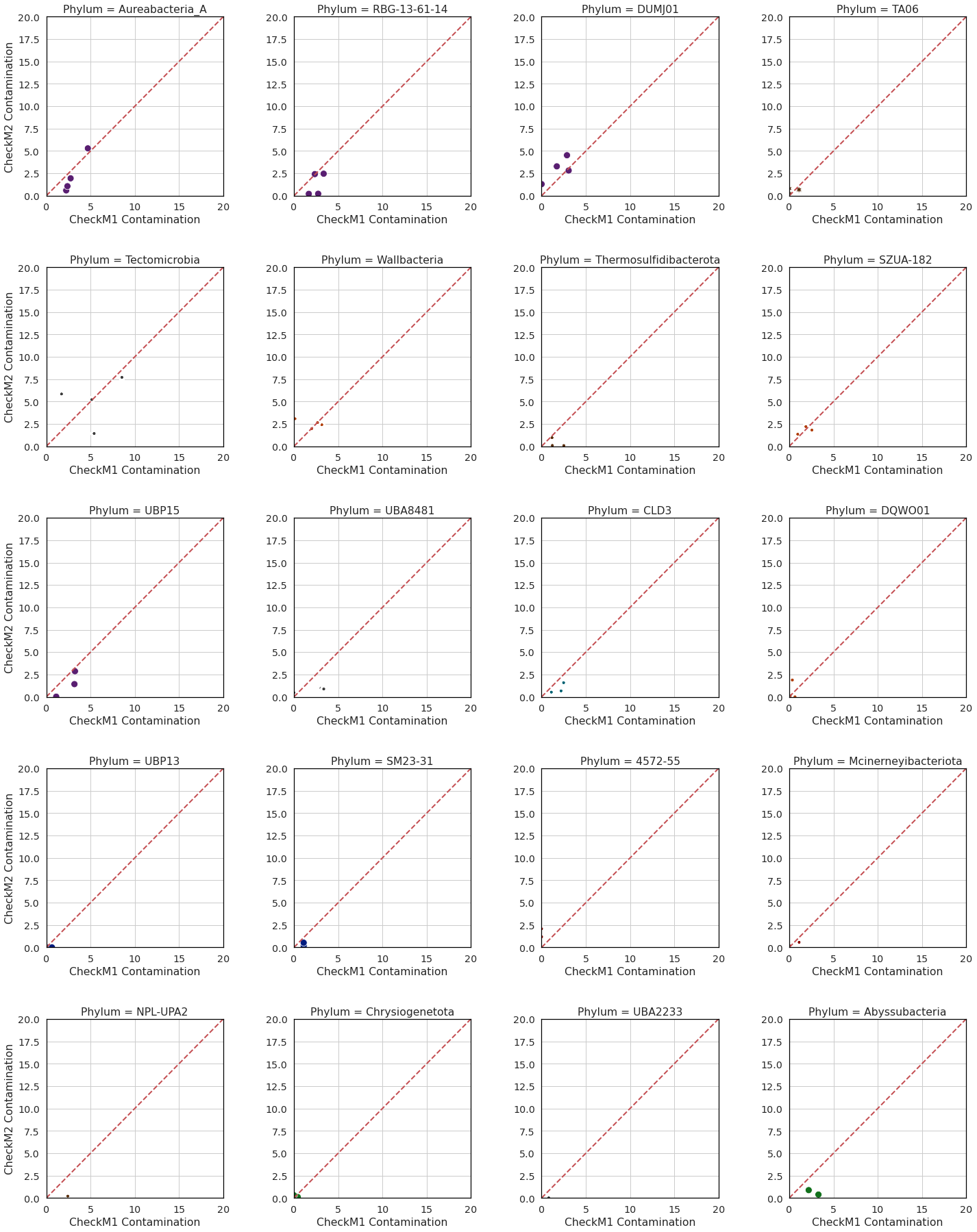

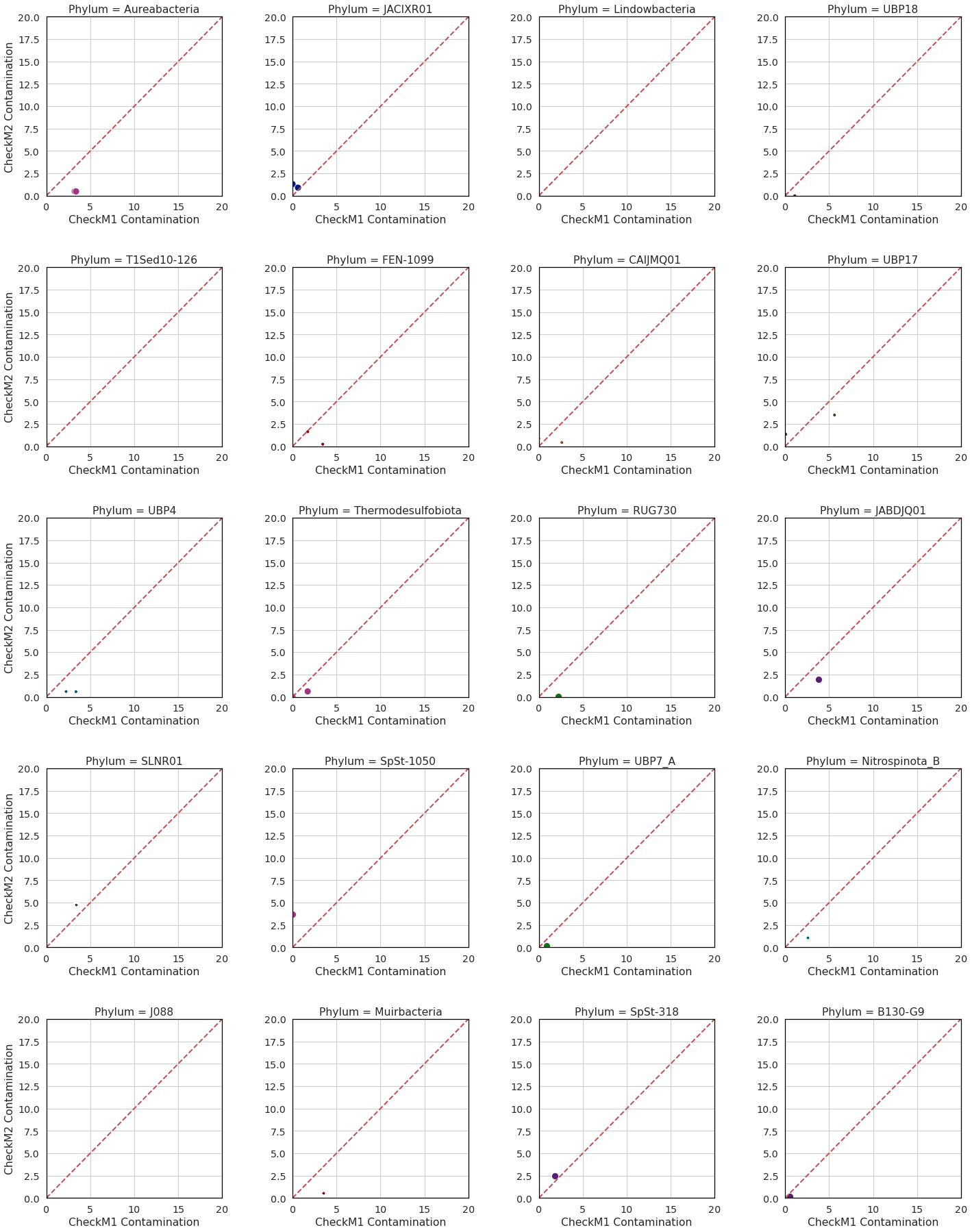

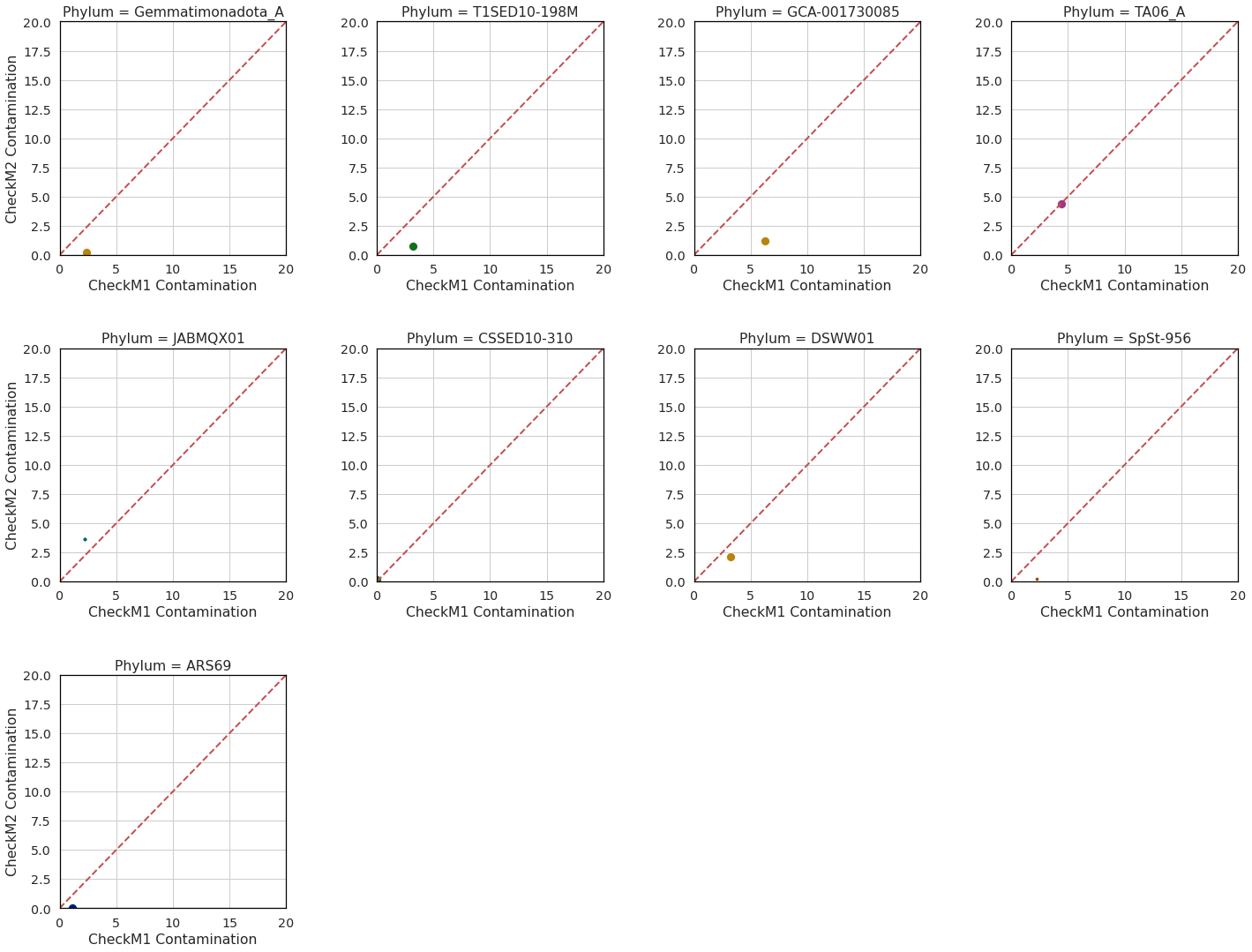

**Supplementary Figure 3**: CheckM2 vs CheckM1 contamination prediction by phylum for **a)** Archaea and **b)** Bacteria MAGs from GTDB release 202.
