## Supplementary Notes 1-8 for "CheckM2: a rapid, scalable and accurate tool for assessing microbial genome quality using machine learning"

**Supplementary Note 1:** CheckM2’s machine learning model structure

Following initial assessment of the accuracy of 11 different machine learning algorithms in predicting genome quality, the two best performing machine learning models (gradient boost and dense neural network) were further optimised. Cross-validation on 700,000 synthetic RefSeq Release 89 genomes was performed on different model architectures for the neural network and parameters for the gradient boost.

For the gradient boost models, cross-validation showed that completeness and contamination was more accurately predicted using different parameters, with the completeness model consisting of relatively shallow trees trained on smaller subsets of data (11 leaves per tree, using a feature fraction of 0.5 and a bagging fraction of 0.5) as compared to the contamination model that consisted of much deeper trees trained on more data (211 leaves per tree, using a feature fraction of 0.9 and a bagging fraction of 0.8).

For the neural network model, the initial dense and fully-connected neural network model showed signs of overfitting when given the full set of genomic inputs (20,021 feature vectors), performing worse on completeness predictions when contamination was higher. The architecture was therefore altered to perform feature extraction and reduce feature space for subsequent layers in the NN. This was achieved using 1-D convolutional layers with a windows size and stride of 10. The entire model consists of four convolutional layers before flattening to a single dense layer connected to the output neuron (**Table 1**). This architecture performed best during cross-validation, outperforming dense neural networks, as well as other convolutional neural network architectures with different number of hidden layers and nodes per layer. Increased accuracy was also obtained by adding batch normalisation layers between the convolutional layers, which substantially reduced error variance, particularly of medium and low-completeness quality predictions.

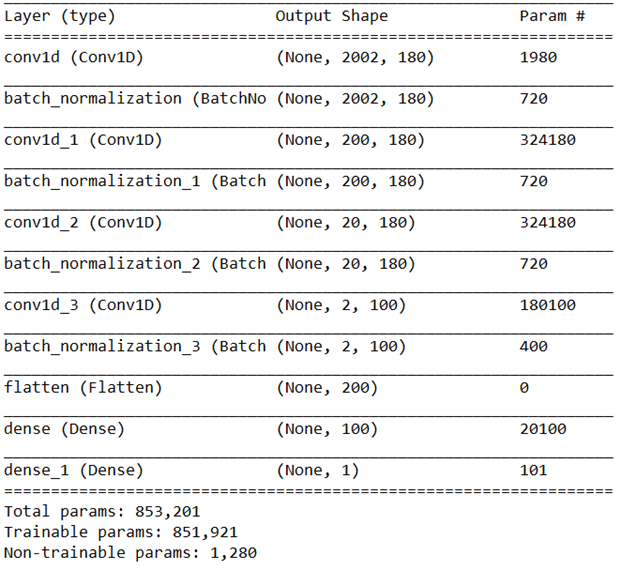

**Table 1:** Structure of optimised neural network model used to predict completeness.

**Supplementary Note 2**: Detailed comparison of CheckM2 performance on RefSeq Release 202 genomes relative to CheckM1 and BUSCO

In order to compare the performance of CheckM2 against CheckM1 and BUSCO for predicting genome quality, each tool was run on a set of synthetic genomes derived from RefSeq release 202 and assessed on a per-phyla basis for completeness (**Figure 1**) and contamination (**Figure 2**). BUSCO and CheckM1 had equivalent performance when predicting contamination, but was less accurate than CheckM1 and CheckM2 in predicting completeness for the vast majority of phyla, sometimes under-predicting completeness relative to CheckM1 and CheckM2 (e.g. **Figure 1a and 1c** in the Thermoproteota; **Figure 1b and 1d** in the Proteobacteria).

The difference in the accuracy of the three tools on the genomes simulated using the 20kb-fragmentation simulation method and the random-MAG-derived simulation method was small, with the all tools being slightly more inaccurate on genomes using the 20kb-fragmentation method (**Figure 1a, 1b** vs **Figure 1c, 1d** and **Figure 2a, 2b** vs **Figure 2c, 2d**). This result indicates that CheckM2’s machine learning models do not appear to be significantly influenced by the type of the simulation strategy. The slightly higher inaccuracy by all methods on the 20-kb-fragmentation is likely the result of a significantly higher average number of contigs in the test genomes (average of 105.6 contigs per genome for 20-kb-fragmentation compared to the average of 66.8 contigs per genome for random-MAG-derived fragmentation). The accuracy of CheckM2’s predictions across the simulated genome test sets gives increased confidence in the robustness of the ML approach for completeness and contamination predictions.

**a)**

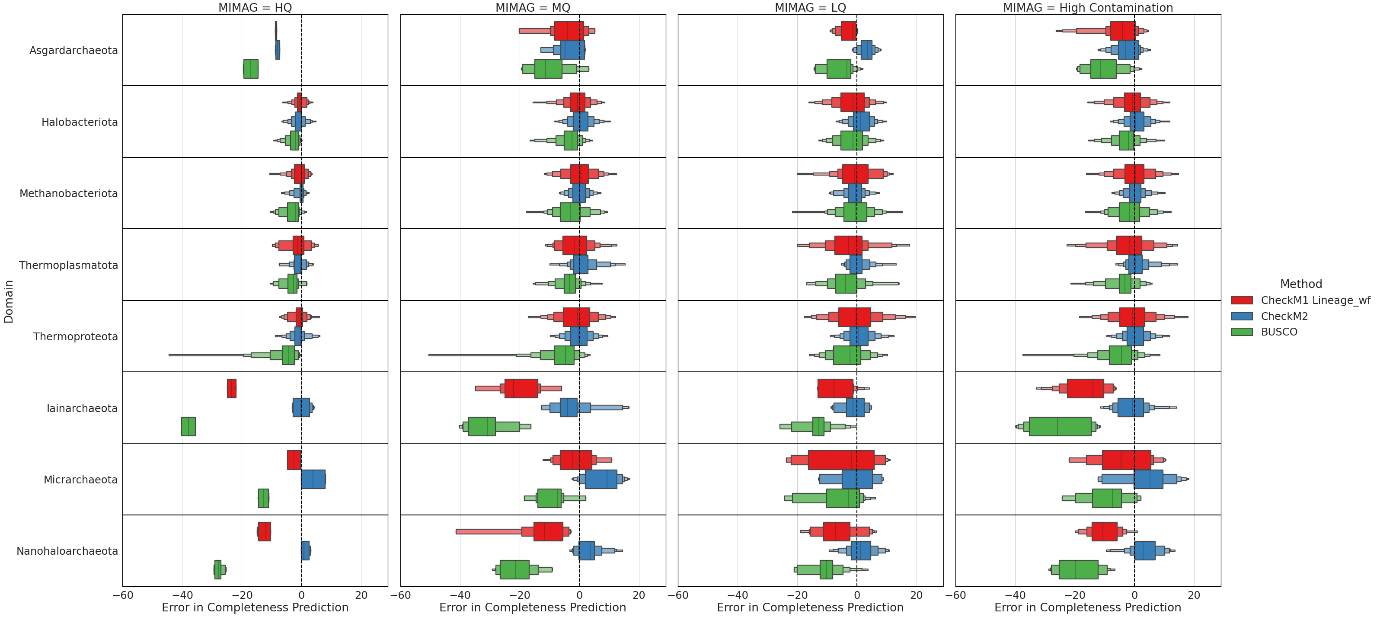

**b)**

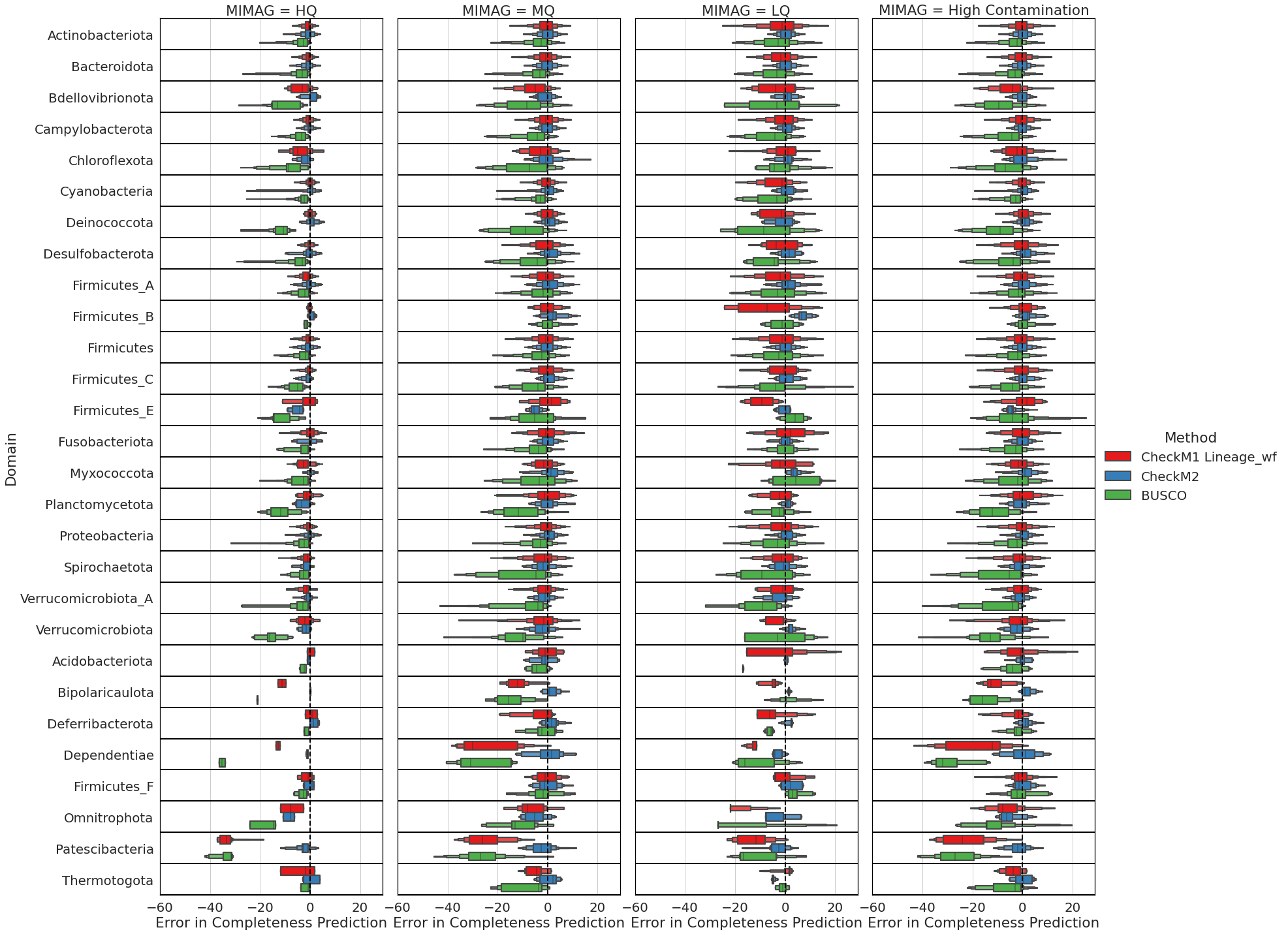

**c)**
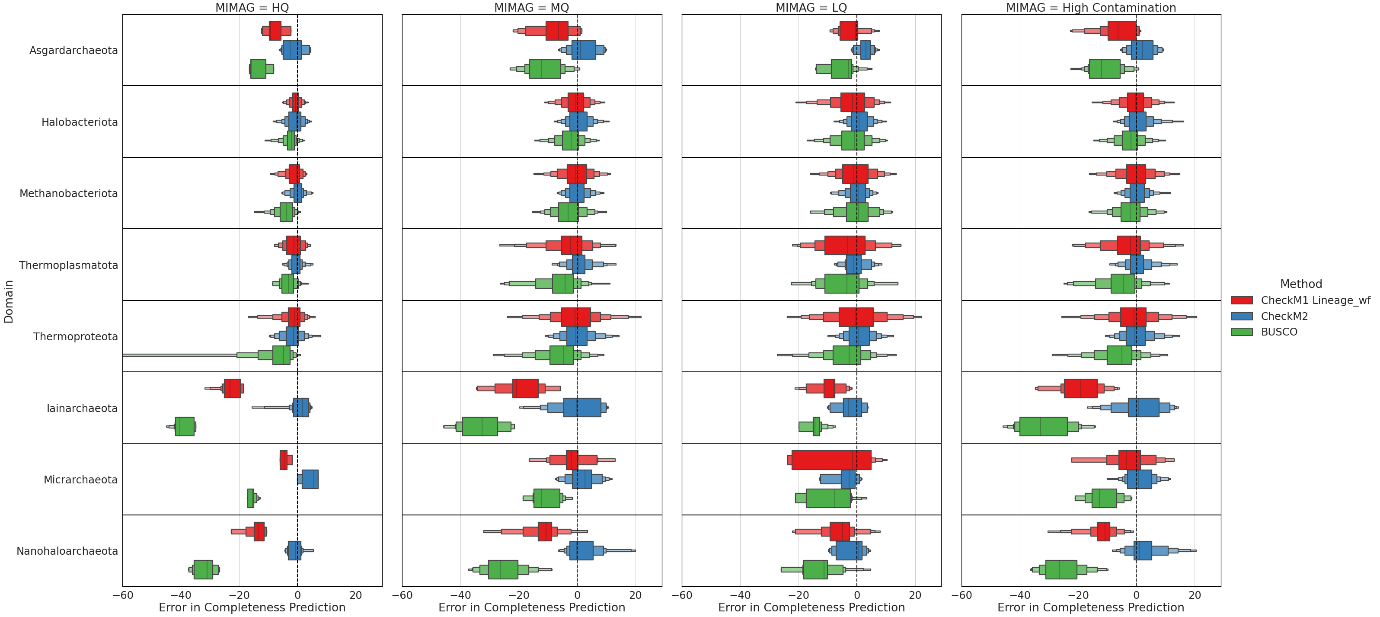

**d)**
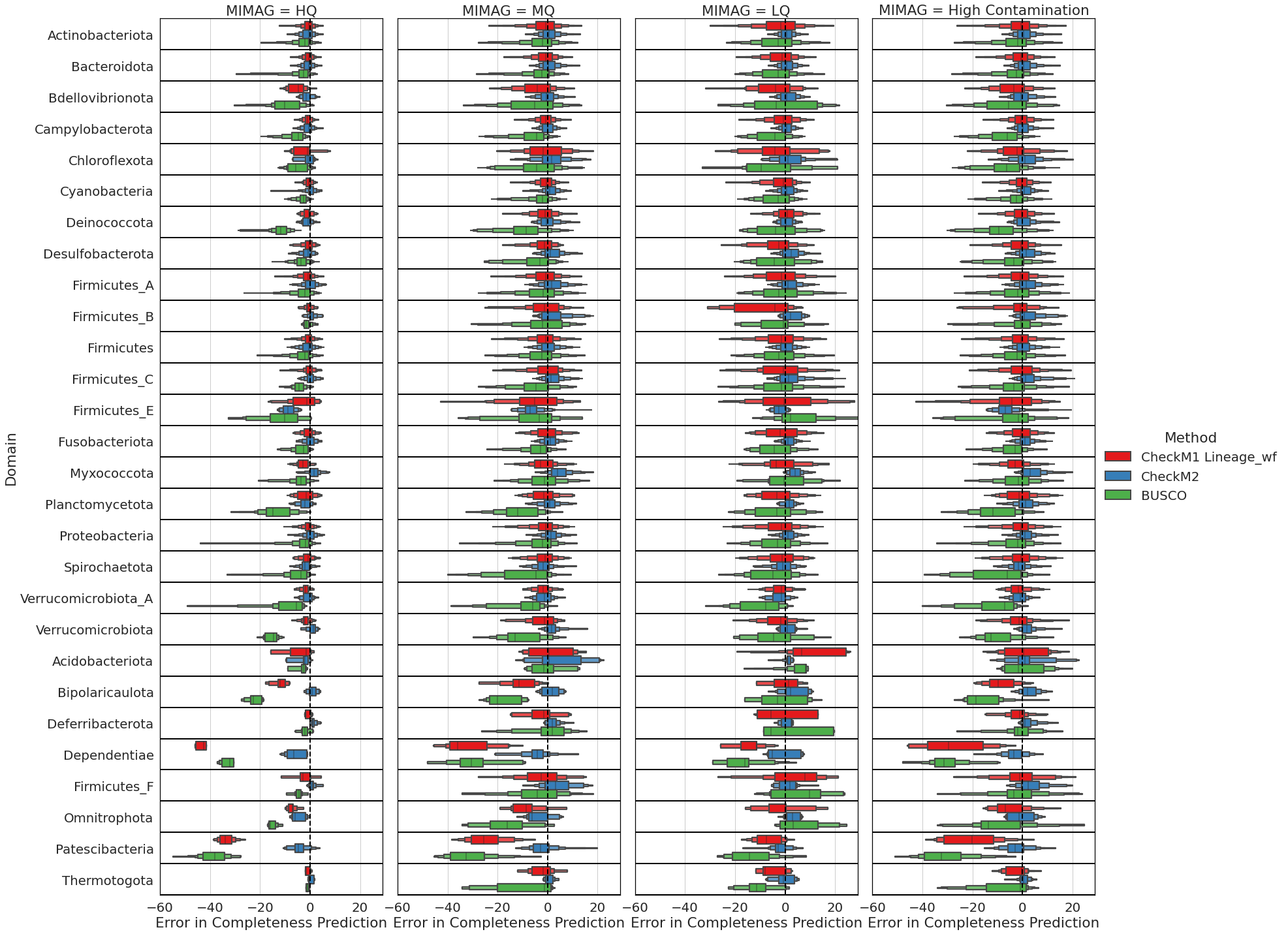

**Figure 1:** Completeness benchmarks by phylum for Archaea and Bacteria in RefSeq release 202 – colours indicate methods (CheckM1, CheckM2 and BUSCO) for **a)** 20-kb-fragmentation method for archaeal phyla, **b)** 20-kb-fragmentation method for bacterial phyla, **c)** random-MAG-fragmentation methods for archaeal phyla, and **d)** random-MAG-fragmentation methods for bacterial phyla.

**a)**
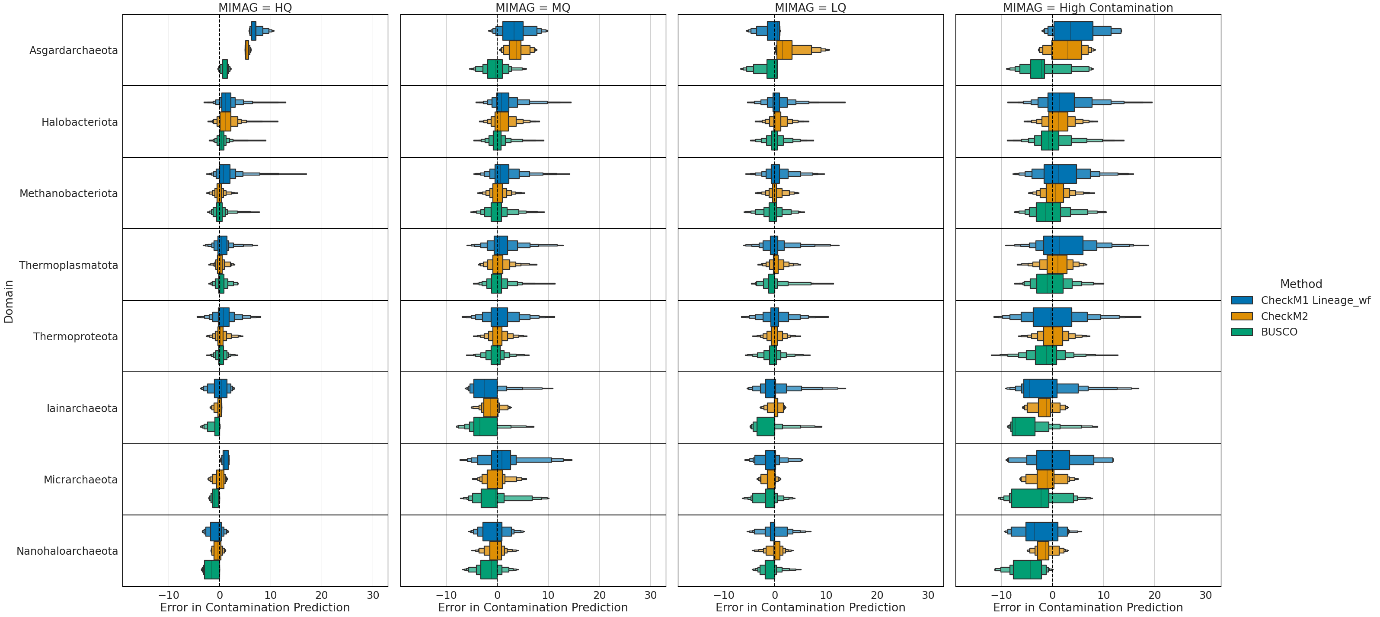

**b)**

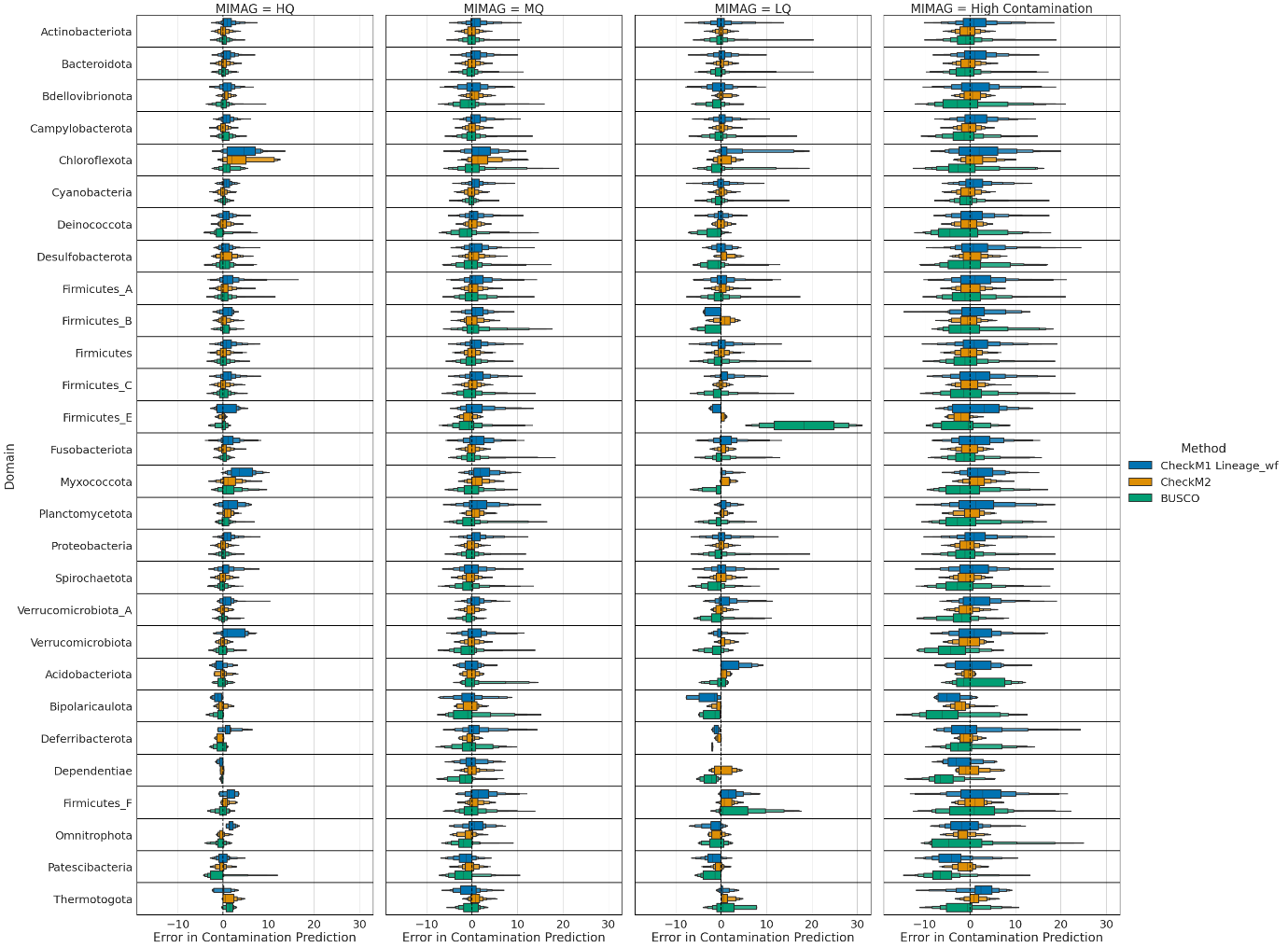

**c)**

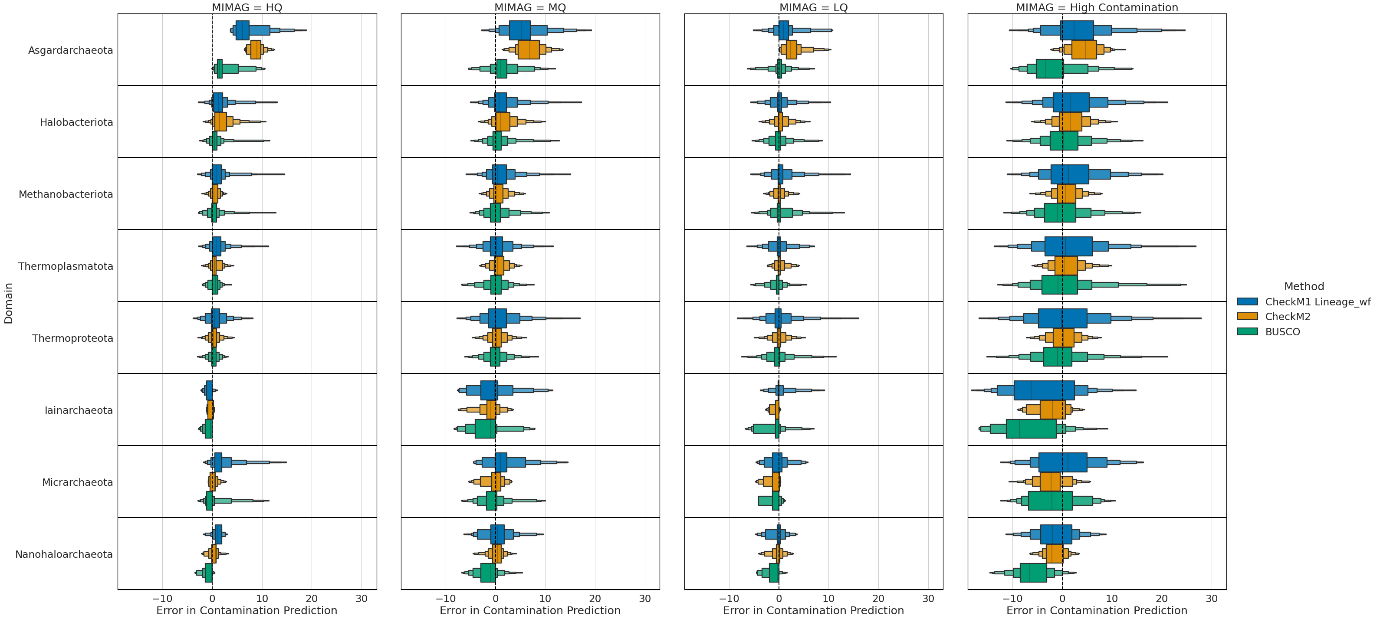

**d)**

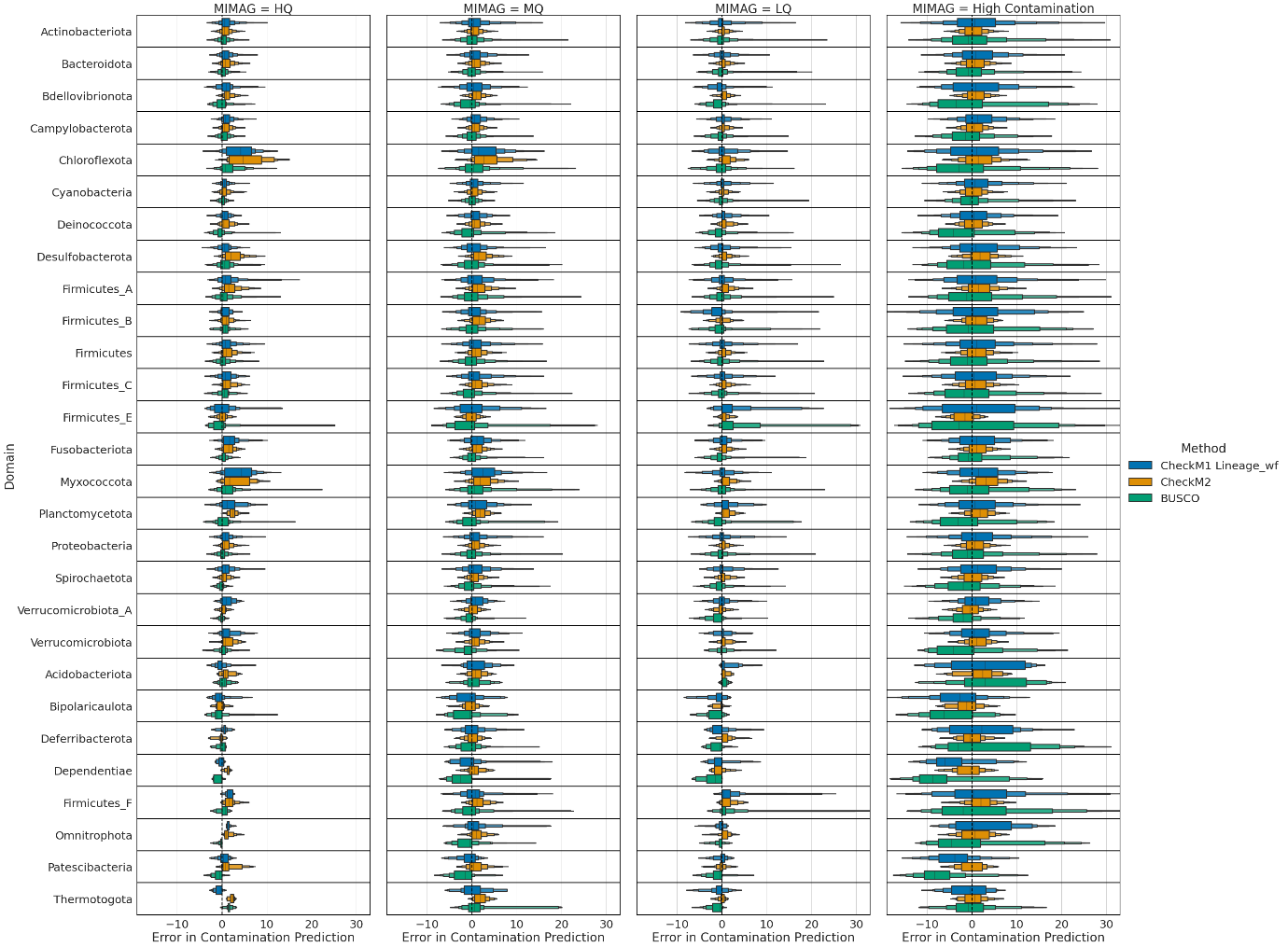

**Figure 2:** Contamination benchmarks by phylum for Archaea and Bacteria in RefSeq release 202 – colours indicate methods (CheckM1, CheckM2 and BUSCO) for **a)** 20-kb-fragmentation method for archaeal phyla, **b)** 20-kb-fragmentation method for bacterial phyla, **c)** random-MAG-fragmentation methods for archaeal phyla, and **d)** random-MAG-fragmentation methods for bacterial phyla.

**Supplementary Note 3**: Detailed comparison of CheckM2 performance on genomes from novel lineages relative to CheckM1 and BUSCO

In order to compare the performance of CheckM2 against CheckM1 and BUSCO for predicting genome quality, each tool was run on a set of synthetic genomes derived from circular MAGs assembled by Singleton et al. and Lui et al. and assessed on a per-phyla basis for completeness (**Figure 3**) and contamination (**Figure 4**). Overall, the results were consistent with prediction on synthetic genomes derived from RefSeq 202, with BUSCO underpredicting completeness across most phyla, especially for genomes from organisms with small genomes such as the Dependentiae, Iainarchaeota and Patescibacteria. CheckM1 also substantially underpredicted the Dependentiae, Proteobacteria and Iainarchaeota MAGs **(Figure 3)**.

As with the RefSeq 203 simulated genomes, CheckM2 shows no significant skews in accuracy between the two different simulation methods and overall outperforms CheckM1 and BUSCO, suggesting that it is robust and accurate on novel MAGs.

**a)**

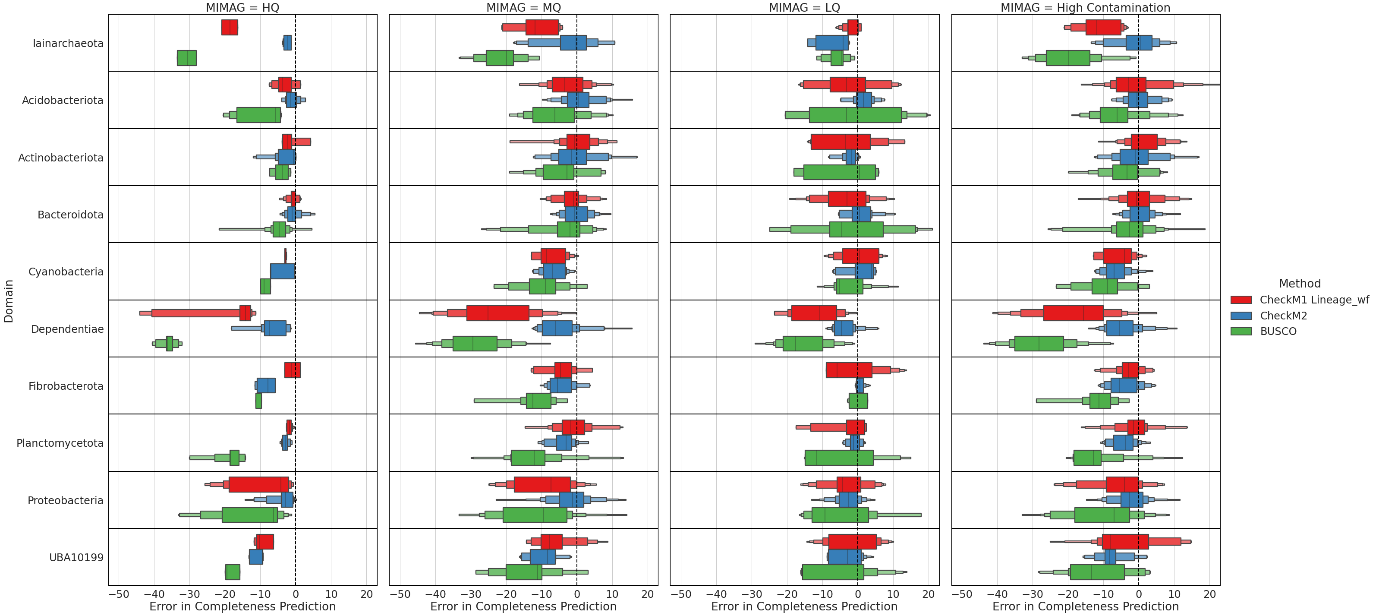

**b)**

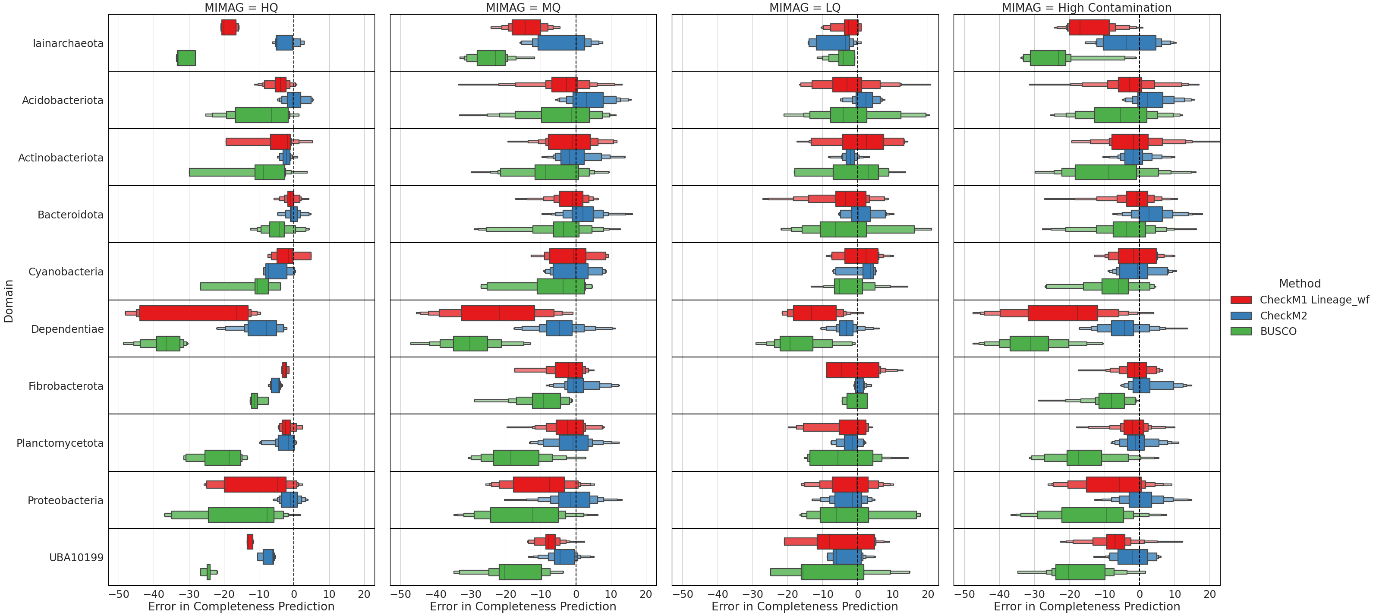

**c)**

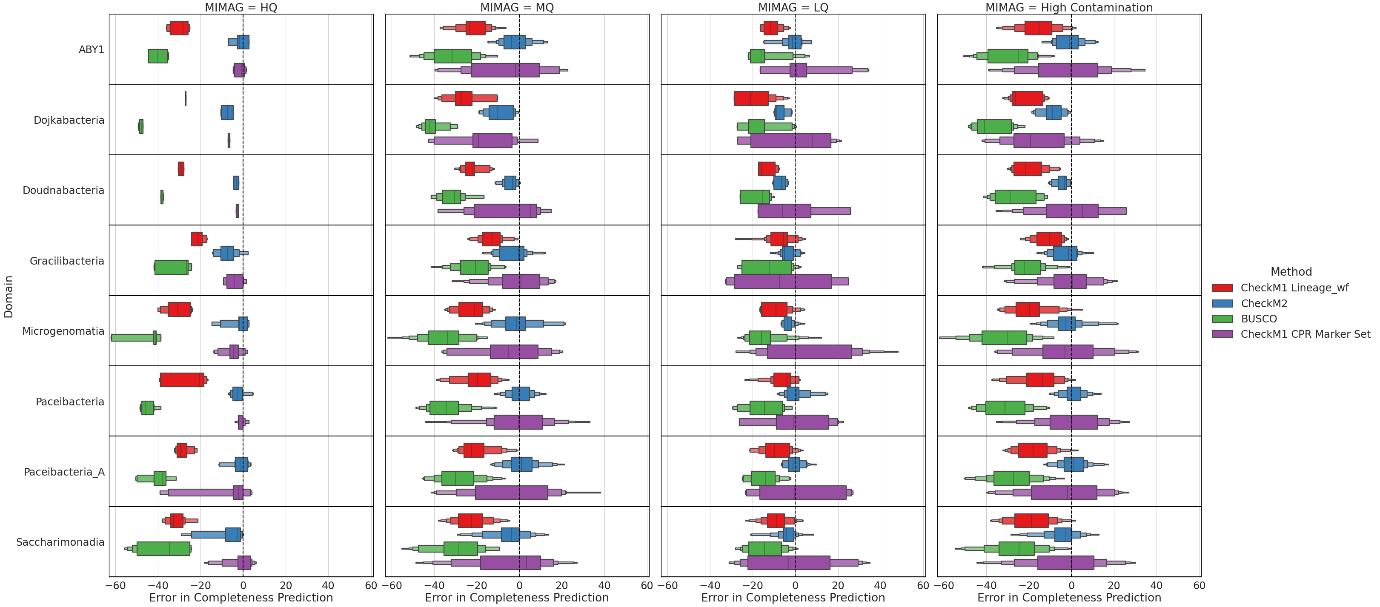

**d)**

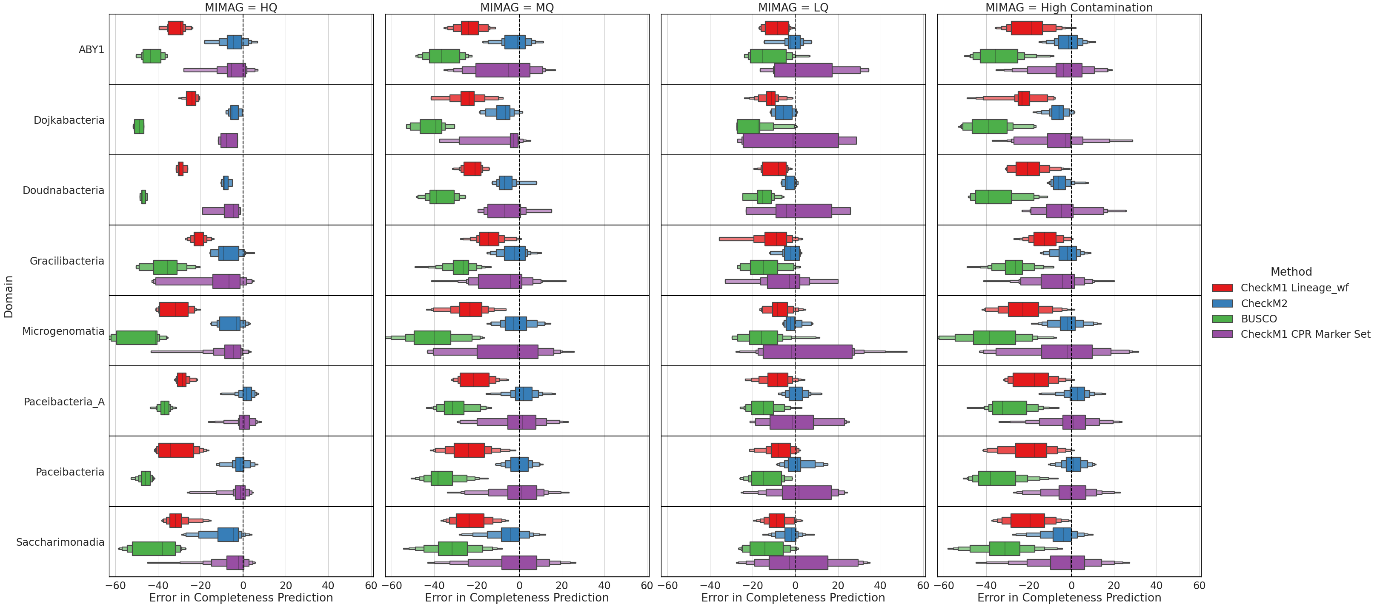

**e)**

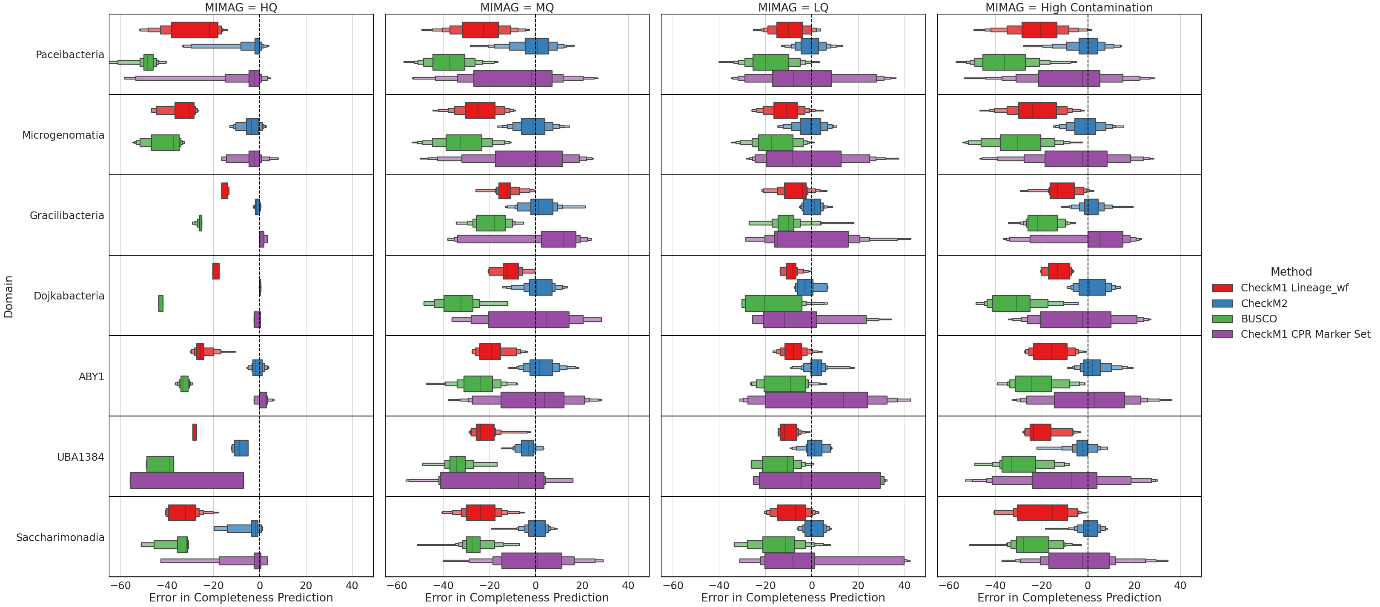

**f)**

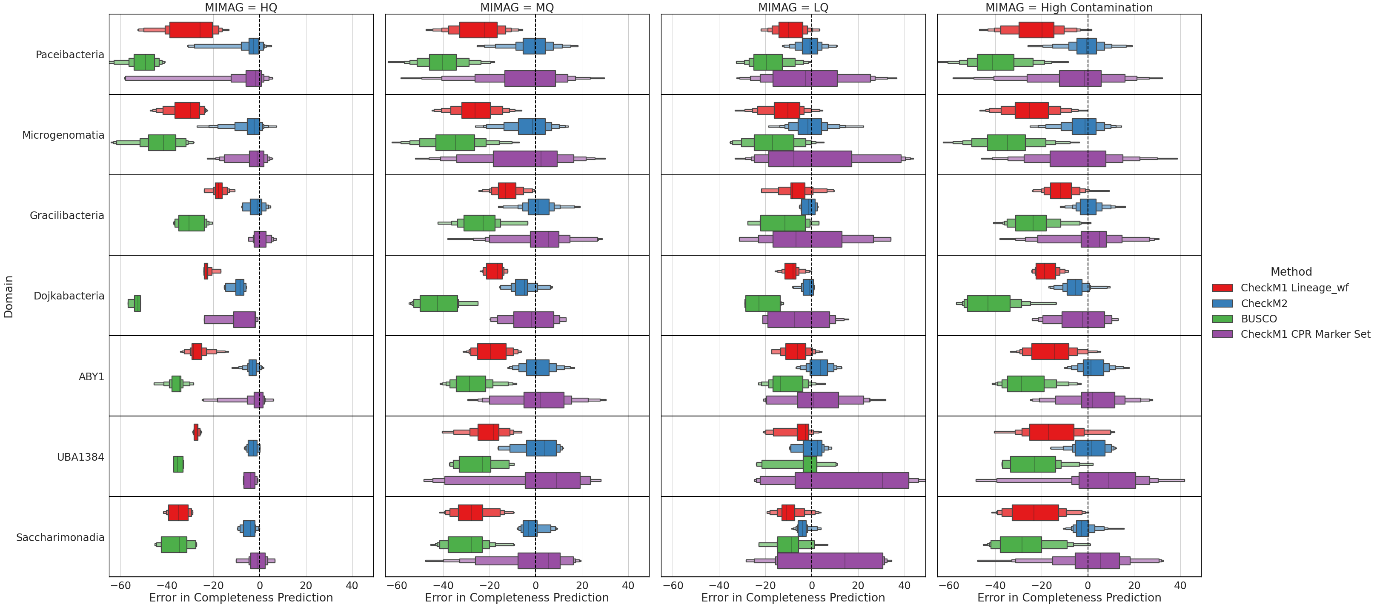

**Figure 3:** Completeness benchmarks by phylum for MAGs from Singleton et al., and Lui et al.; colours indicate methods (CheckM1 Lineage_wf, CheckM1 marker set, CheckM2 and BUSCO) for **a)** 20-kb-fragmentation method for Singleton et al., phyla; **b)** random-MAG-fragmentation for Singleton et al., phyla; **c)** 20-kb-fragmentation method for Patescibacteria genomes from Singleton et al., by class; **d)** random-MAG-fragmentation methods for Patescibacteria genomes from Singleton et al., by class; **e)** 20-kb-fragmentation methods for Patescibacteria genomes from Lui et al., by class; and **f)** random-MAG-fragmentation methods for Patescibacteria genomes from Lui et al., by class.

**a)**

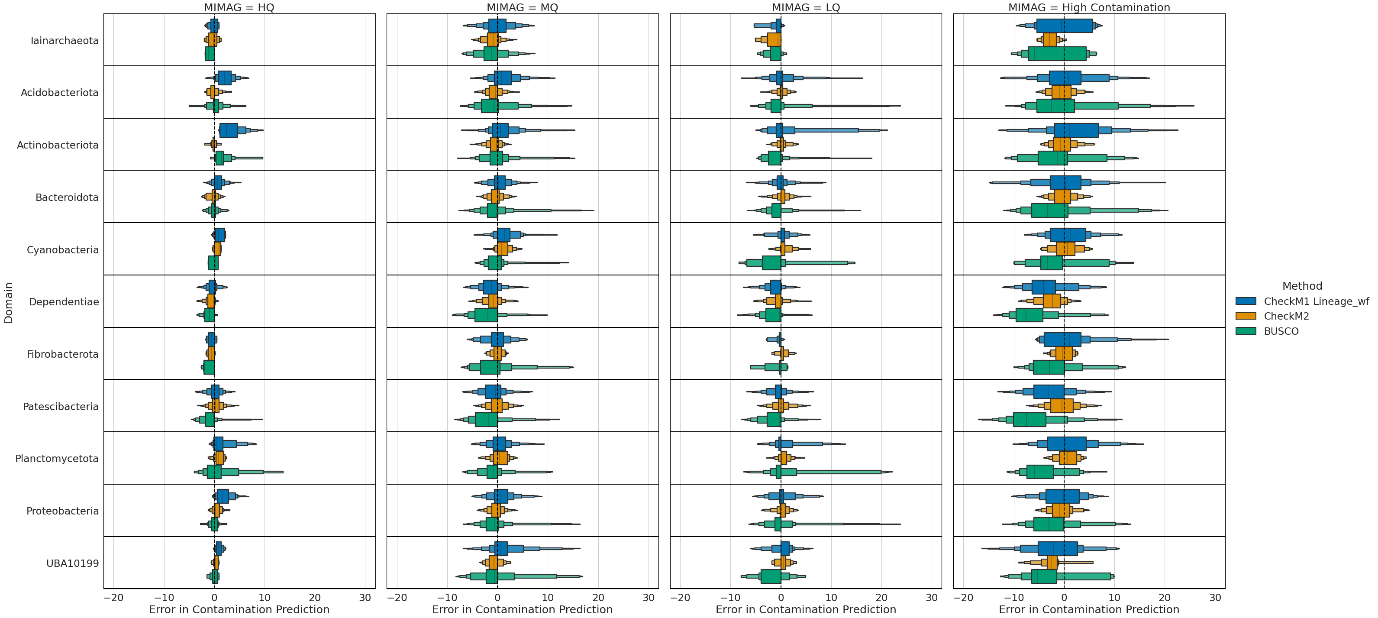

**b)**

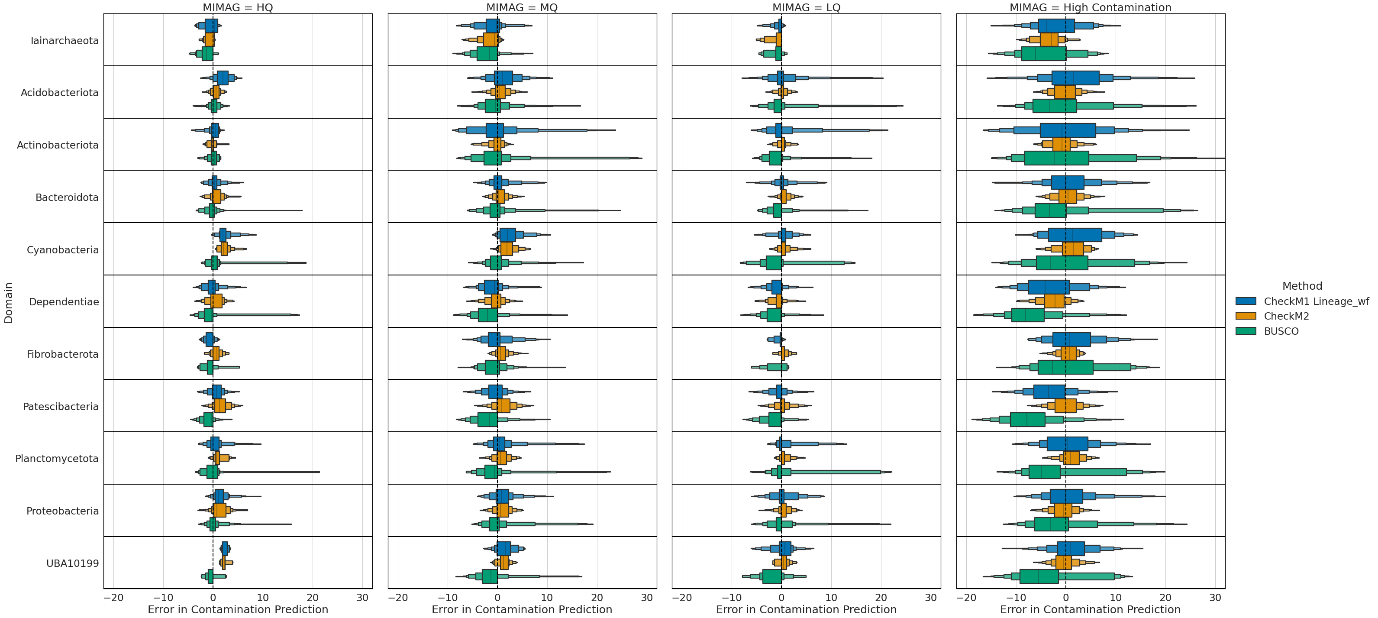

**c)**

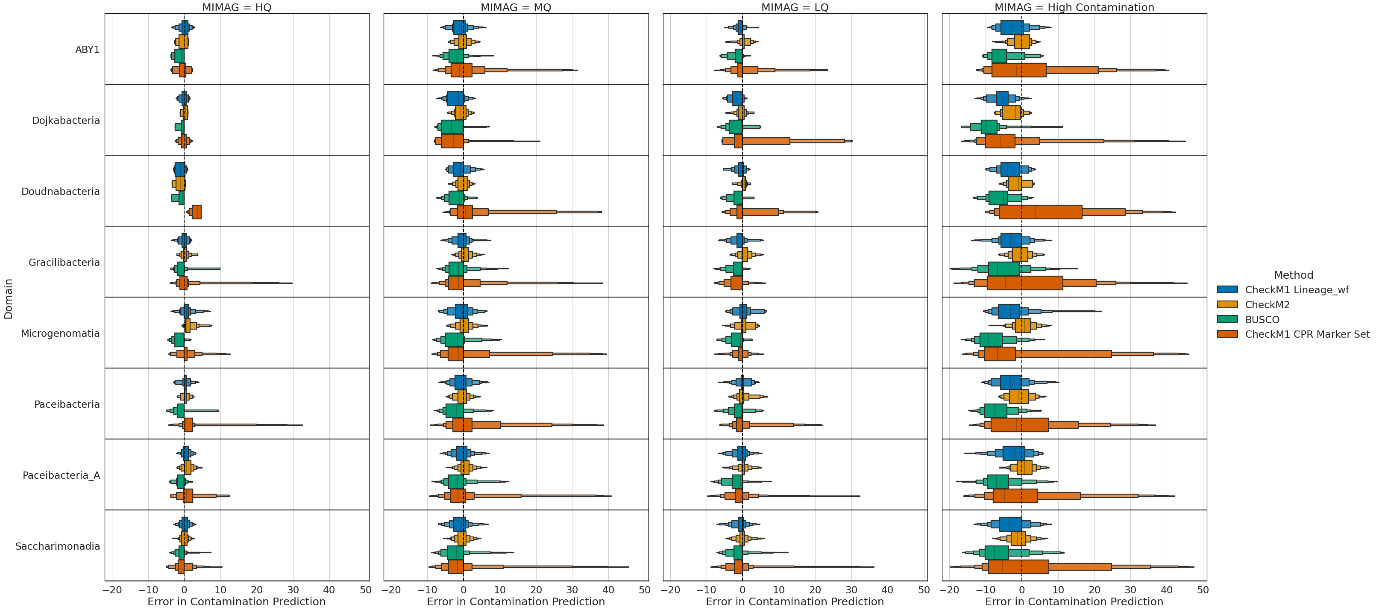

**d)**

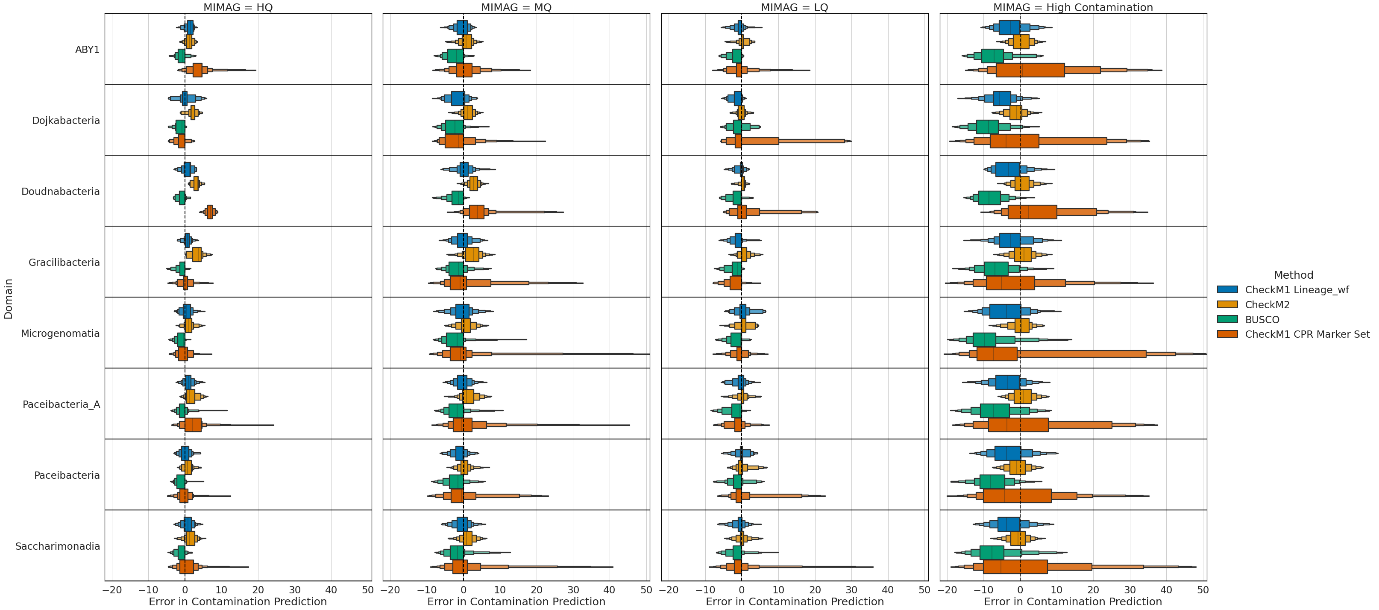

**e)**

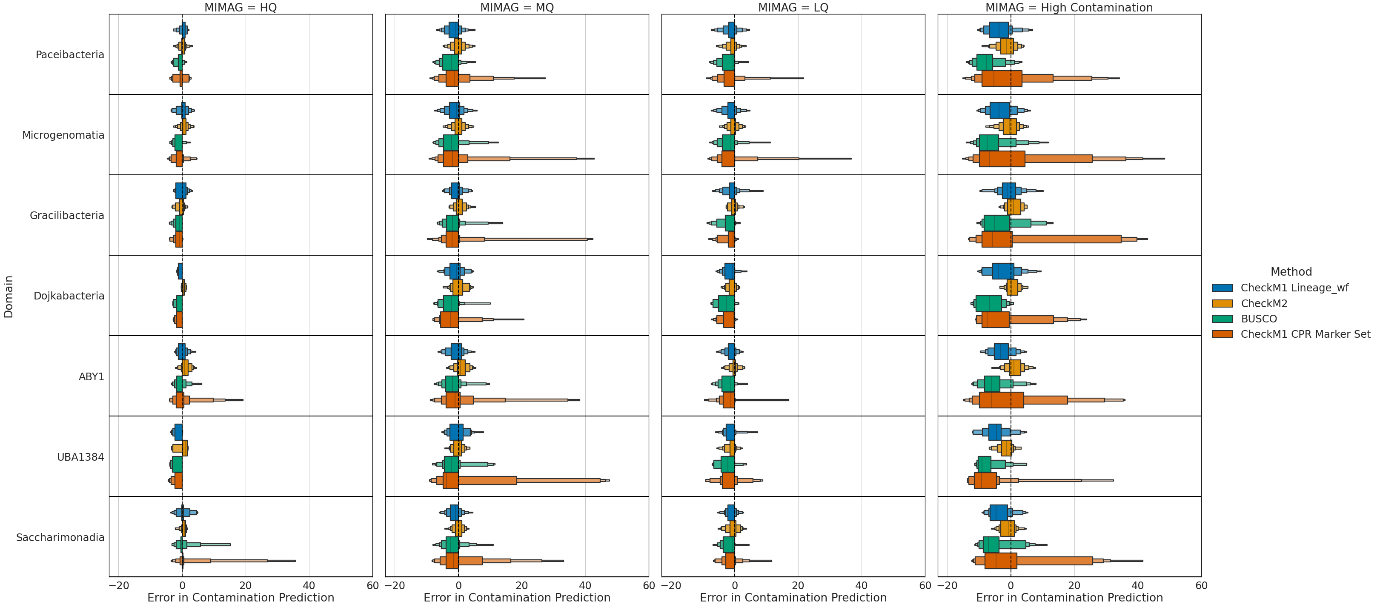

**f)**

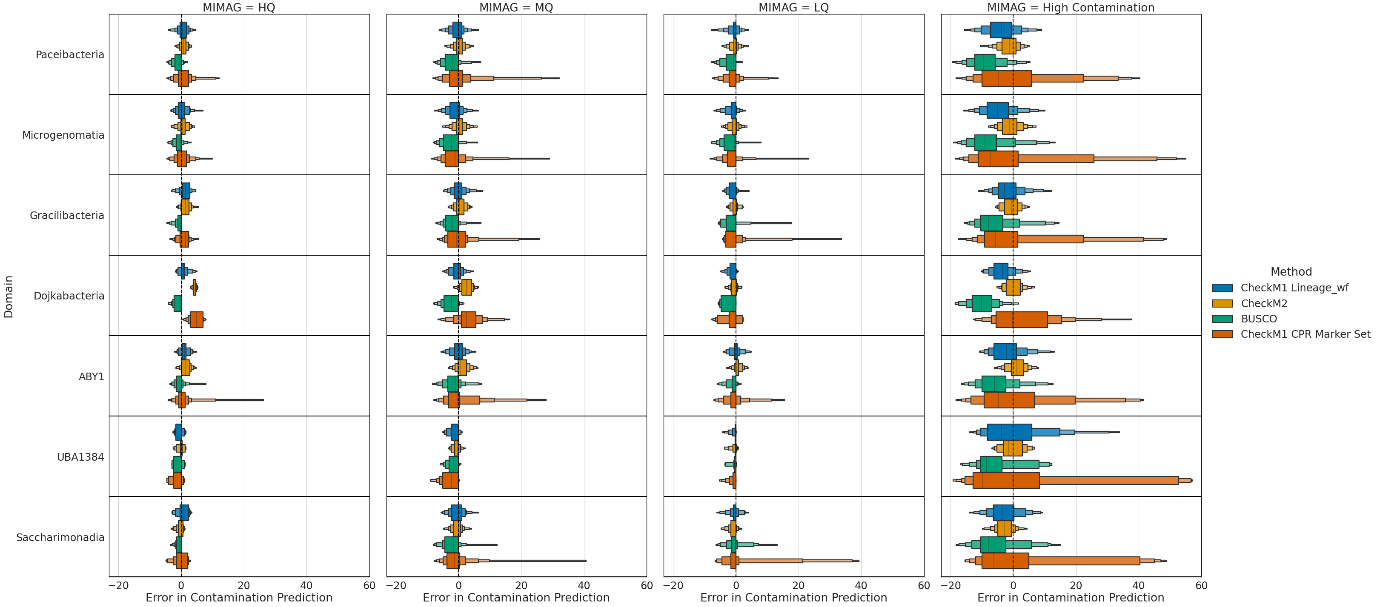

**Figure 4:** Contamination benchmarks by phylum for MAGs from Singleton et al., and Lui et al.; colours indicate methods (CheckM1 Lineage_wf, CheckM1 marker set, CheckM2 and BUSCO) for **a)** 20-kb-fragmentation method for Singleton et al., phyla; **b)** random-MAG-fragmentation for Singleton et al., phyla; **c)** 20-kb-fragmentation method for Patescibacteria genomes from Singleton et al., by class; **d)** random-MAG-fragmentation methods for Patescibacteria genomes from Singleton et al., by class; **e)** 20-kb-fragmentation methods for Patescibacteria genomes from Lui et al., by class; and **f)** random-MAG-fragmentation methods for Patescibacteria genomes from Lui et al., by class.

**Supplementary Note 4:** Investigation of cross-contamination predictions by CheckM1 and CheckM2

Examination of contamination predictions by CheckM1 and CheckM2 in the synthetically generated cross-contaminated genomes show that the less taxonomically related the contaminant contig is to the source genome, the less like it is to be detected. However, CheckM1 shows a substantial amount of overprediction, in some cases predicting as much as 40% contamination where known genome contamination never exceeds 10%. This overprediction is exhibited primarily by CheckM1, with 1954 cases of overpredictions by 10% or more compared to 64 by CheckM2 (**Table 2**).

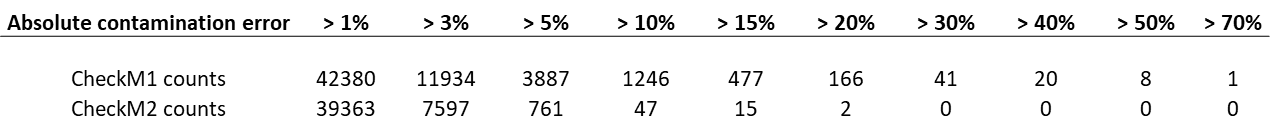

**Table 2**: Absolute counts of predictions by CheckM1 and CheckM2 by absolute contamination error for all genomes where real contamination does not exceed 5% of genome.

Notably, the overprediction by CheckM1 shifts its median prediction error artificially higher; when the data is divided into separate underprediction and overprediction distributions, CheckM1 performs roughly the same as CheckM2 when underpredicting, and is vastly outperformed by CheckM2 when overpredicting (**Figure 5** vs **Figure 6**).

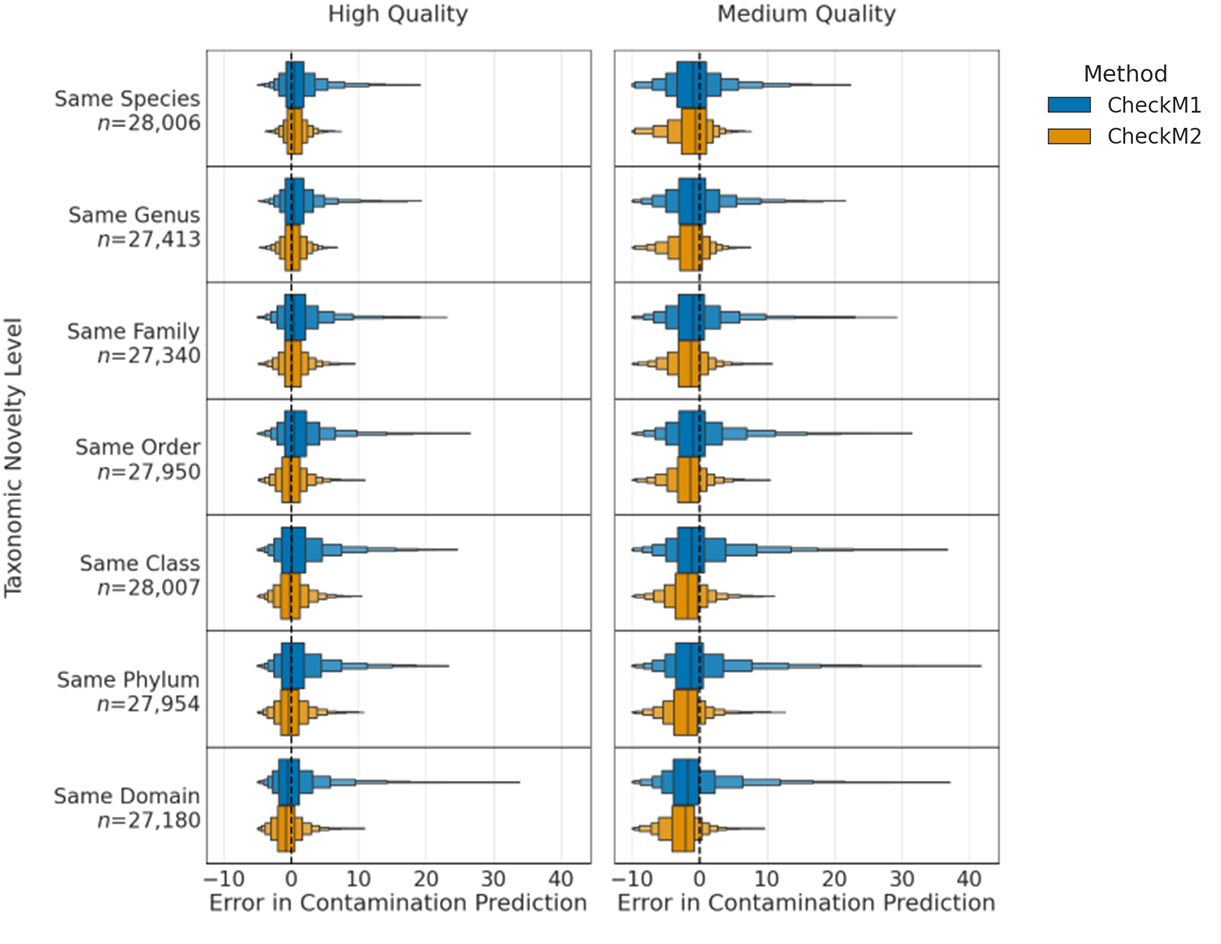

**Figure 5**: Absolute error of contamination predictions (predicted – actual) for genomes with non-self contamination across different levels of contaminant taxonomy relative to contaminated genome. Each box shows half the remaining data, starting with 50% for the first and 25% for the second. Eight boxes are shown, comprising 99.61% of all data.

**a	b**

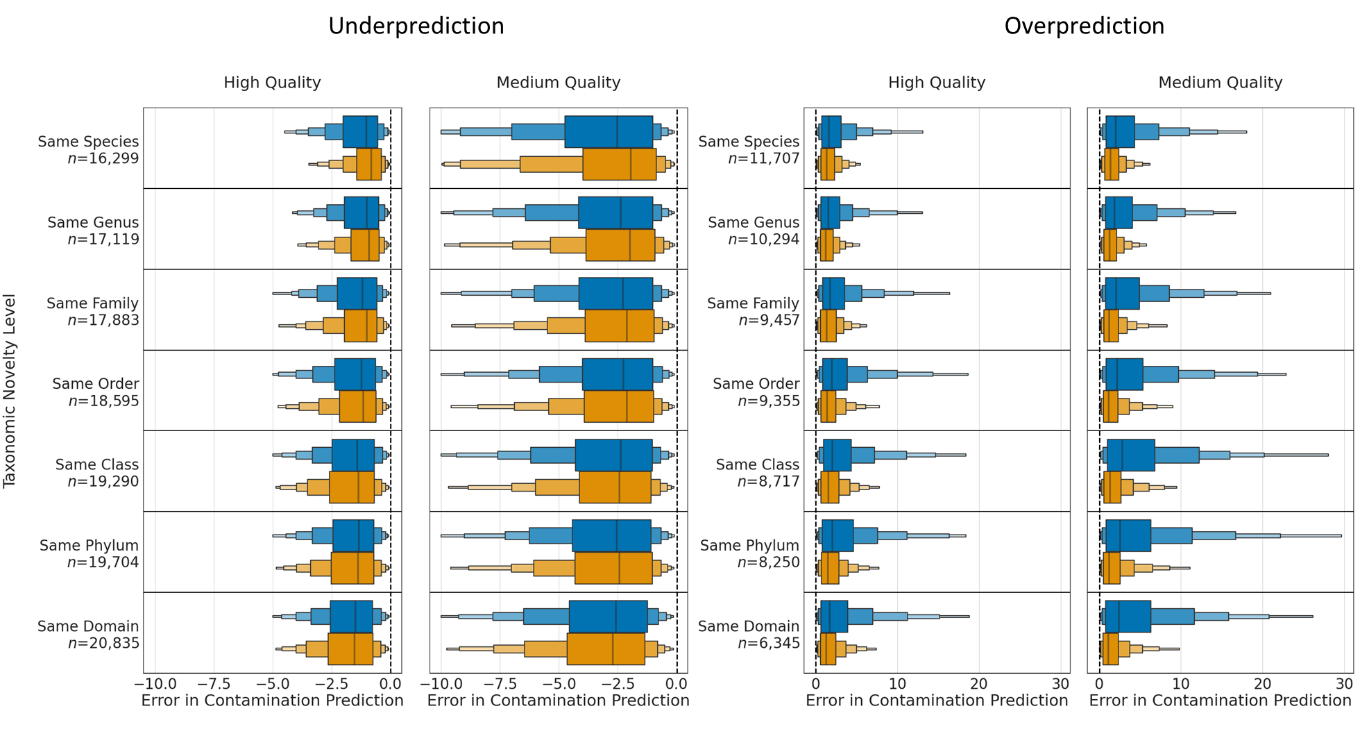

**Figure 6**: CheckM1 and CheckM2 contamination error across taxonomic levels of contamination, broken into **a)** underprediction only, and **b)** overprediction only.

These overpredictions are not confined to specific taxa and occur in all phyla represented in the synthetic genome pool, nor are they confided to any specific contaminant contig taxonomy. However, cases where CheckM1 significantly overestimates contamination are predominantly assessed using a generic marker set such as k__Bacteria (UID203). It appears that the incongruent taxonomy of the contig vs the genome pushes CheckM1’s placement of the genome into the reference tree further towards the root, and the single-copy genes present on some of the randomly sampled contaminating contigs are counted towards contamination. Because the marker set is generic and small, only a comparatively small number of single-copy marker genes found on the contaminating contig are needed to predict an inaccurately high contamination. CheckM2 on the other hand is not substantially biased by a relatively small number of duplicated single-copy marker genes and rarely overpredicts contamination at any taxa. This, combined with its superior performance on same-species, same-genus, same-family and same-order contamination relative to CheckM1 (**Figure 6b**) will lead to more accurate assessments of contamination in most cases of co-binning related contigs incorrectly, while other tools that take into account taxonomy of contamination (e.g. GUNC^1^) can more accurately identify foreign contamination from a distant taxonomic source.

**Supplementary Note 5**: An example of taxonomically divergent completeness predictions between CheckM1 and CheckM2 where CheckM1 predictions are higher

While most genomes in GTDB release 202 show robust congruence in completeness predictions, there are genomes where either CheckM2 or CheckM1 predictions are higher. Most higher CheckM2 completeness predictions are associated with genomes from microorganisms hypothesized to be undergoing genome reduction, streamlining, or have genomic features associated with symbiosis or parasitism (see Results, section ***Comparison of CheckM1 vs CheckM2 predictions across all bacteria and archaea***). Higher CheckM1 predictions do not appear to show significant correlation to taxonomy or known biological features with some exceptions.

One key exception is the phylum Dormibacterota, which shows on average 3-7% lower CheckM2 completeness scores across all genomes in this phylum **(Figure 7)**. This is the only sizable phylum to show such systematic deviation.

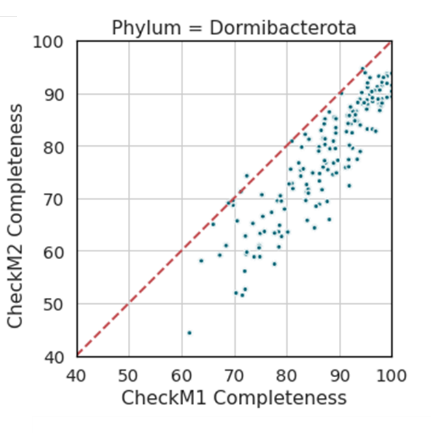

**Figure 7:** Completeness predictions of genomes in the phylum Dormibacterota by CheckM1 versus CheckM2.

Analysis of annotated KOs, modules and pathways as well as output from the SHAP^2^ package indicate that the main likely contributor to the lower completeness score of CheckM2 is the KEGG Ribosomal pathway (M00178). The Dormibacterota has one of the lowest mean percentage completeness in the Ribosomal pathway (72%) among all phyla with more than 5 representatives. This is even lower than Patescibacteria which are known to lack key genes in this pathway. Genomes in the Dormibacterota appear to have lower counts of the large subunit ribosomal proteins L15, L21, L22, L29, L32 and the small subunit ribosomal proteins S3, S16, and S20, with these genes only found in between 15-58% of genomes (average 39%). The reason that CheckM1 fails to identify this pattern is likely due to CheckM1 grouping many ribosomal marker genes into marker sets, which lowers its resolution relative to CheckM2 for this pathway.

The CheckM1 marker set used to assess most Dormibacterota (UID2307, k__Bacteria), only has S16 and S20 as separate single-copy marker genes. S3 and L22 (average mean count in Dormibacterota genomes: 0.23 and 0.53) are in a single marker set with a further 9 ribosomal genes (K02886, K02878, K02906, K02892, K02965, K02904, K02926, and K02961) for which the mean count is 0.83 ± 0.03. However, in the Dormibacterota genomes, S3 and L22 are not located next to the other 9 ribosomal genes, and therefore CheckM1’s marker set assumptions are not appropriate for this lineage. This is also the case for the ribosomal proteins L15, L21, L29 and L32, which are in CheckM1 marker sets whose synteny assumption does not hold for genomes within the Dormibacterota, and whose gene counts are substantially lower on average than other genes within these marker sets.

In summary, CheckM2’s lower completeness predictions are congruent with the underlying genomic data. CheckM1’s marker set was built on ribosomal protein synteny assumptions which do not hold for Dormibacterota, making CheckM1’s scores potentially unreliable. Sequencing high-quality isolate genomes of Dormibacterota in the future will determine if this reflects low-quality assembly or an important biological difference, while also allowing CheckM2 to incorporate this information its machine learning models.

**Supplementary Note 6**: Systematic investigation of indel effects on divergent predictions between CheckM1 and CheckM2 of MAGs from GTDB release 202

In order to investigate general patterns associated with higher CheckM1 scores, genomes from species with more than 10 genomic representatives from Archaea and Bacteria were selected where at least a third or more showed substantially higher CheckM1 values. Selecting well-represented species allows examination of genome attributes such as genome size or protein count along a gradient of predicted completeness by CheckM1 and CheckM2.

Interestingly, out of 26,000 genomes with substantial differences between CheckM1 and CheckM2 completeness predictions, only 71 (0.27%) showed consistently higher CheckM1 scores across all levels of completeness. The vast majority (25,783 or 99.3% of all genomes) showed a non-linear deviation where CheckM1 and CheckM2 scores were largely congruent for the most complete genomes (predicted to be 90-100% complete), but showed a marked deviation as predicted genome completeness declined, with CheckM1 scores being much higher than CheckM2 at lower levels of completeness (**Figure 8a, 8b**).

**a)**

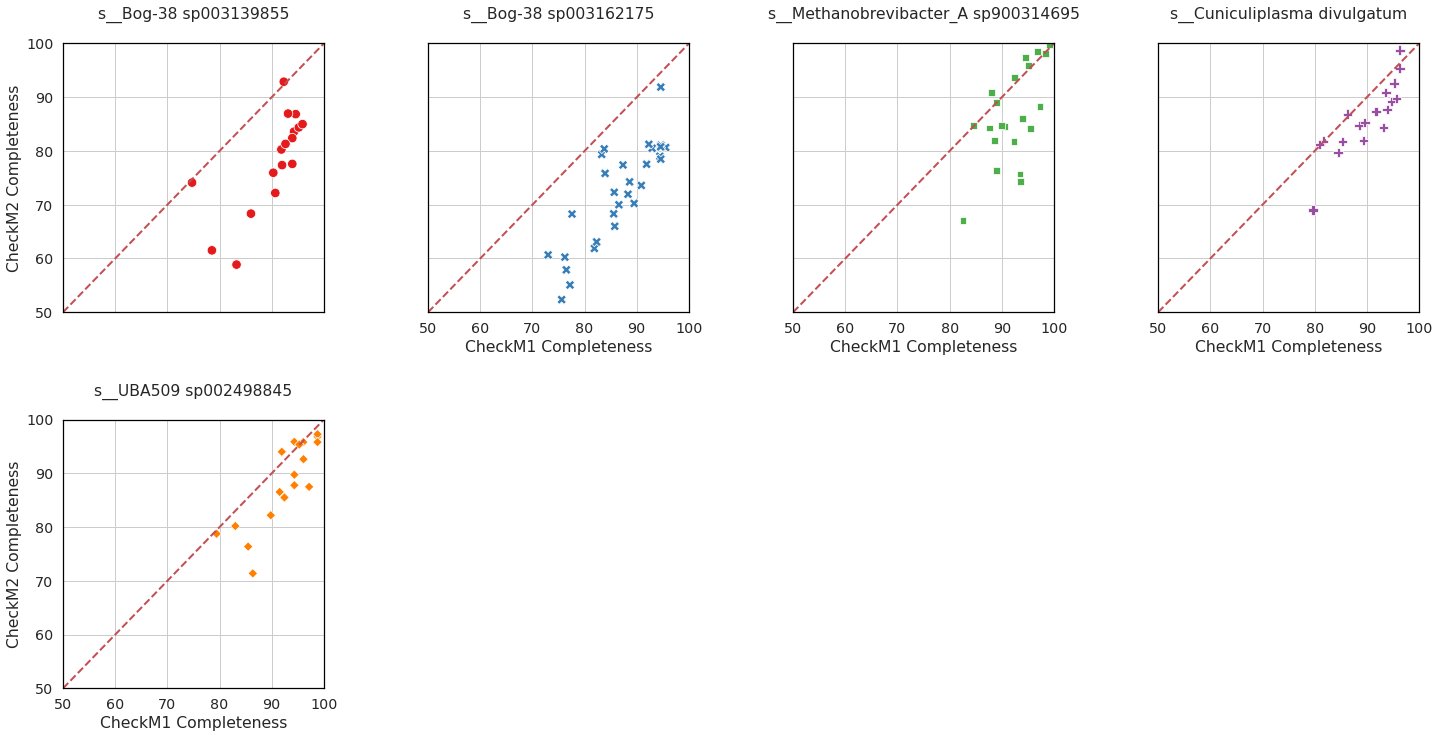

**b)**

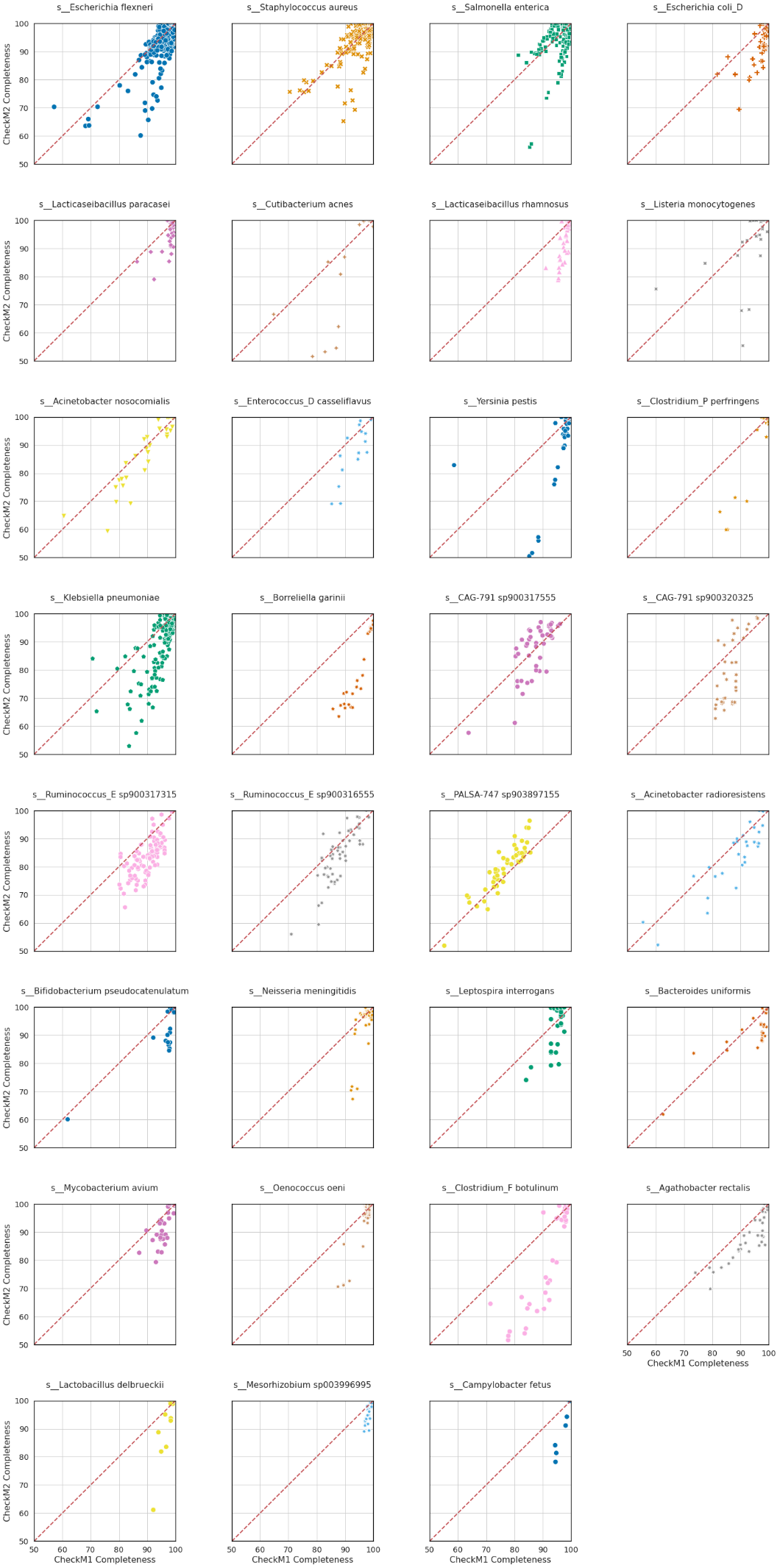

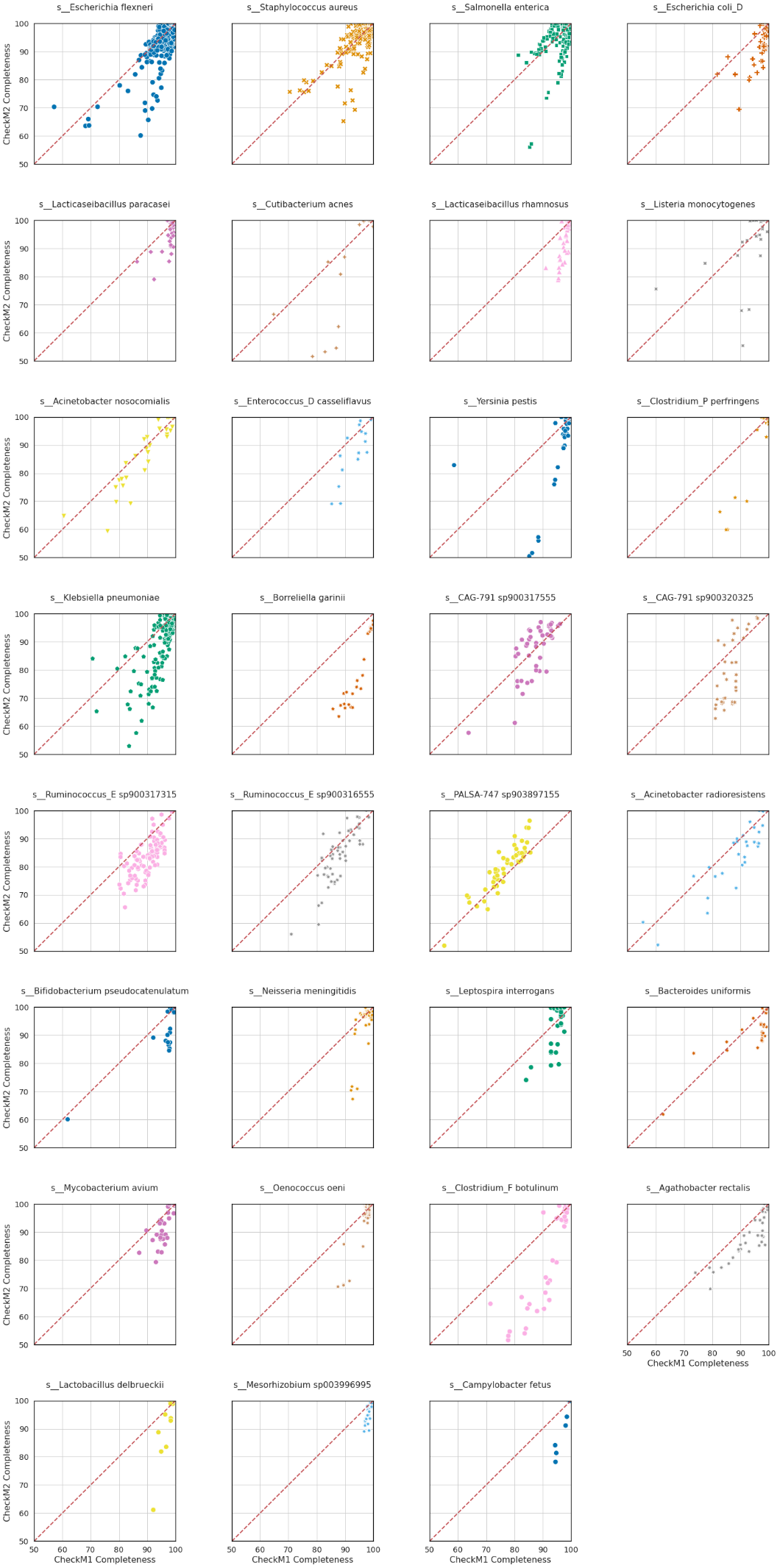

**Figure 8:** Genome completeness predictions of CheckM1 vs CheckM2 where CheckM1 scores are higher for all or some of the genomes, for a) archaeal species, and b) bacterial species.

The primary indicator of substantially lower CheckM2 scores relative to CheckM1 are genomes hypothesized to contain indels and assembled using long-read technology such as PacBio or Nanopore. NCBI assembly metadata marks suspected indel-dominated genomes as ‘containing many frame-shifted proteins’, identified through unusually high CDS count comparison to high-quality isolate genomes in its database. For those species with genomes containing suspected indels, there is a strong direct correlation between low CheckM2 scores versus CheckM1 **(Figure 9)**.

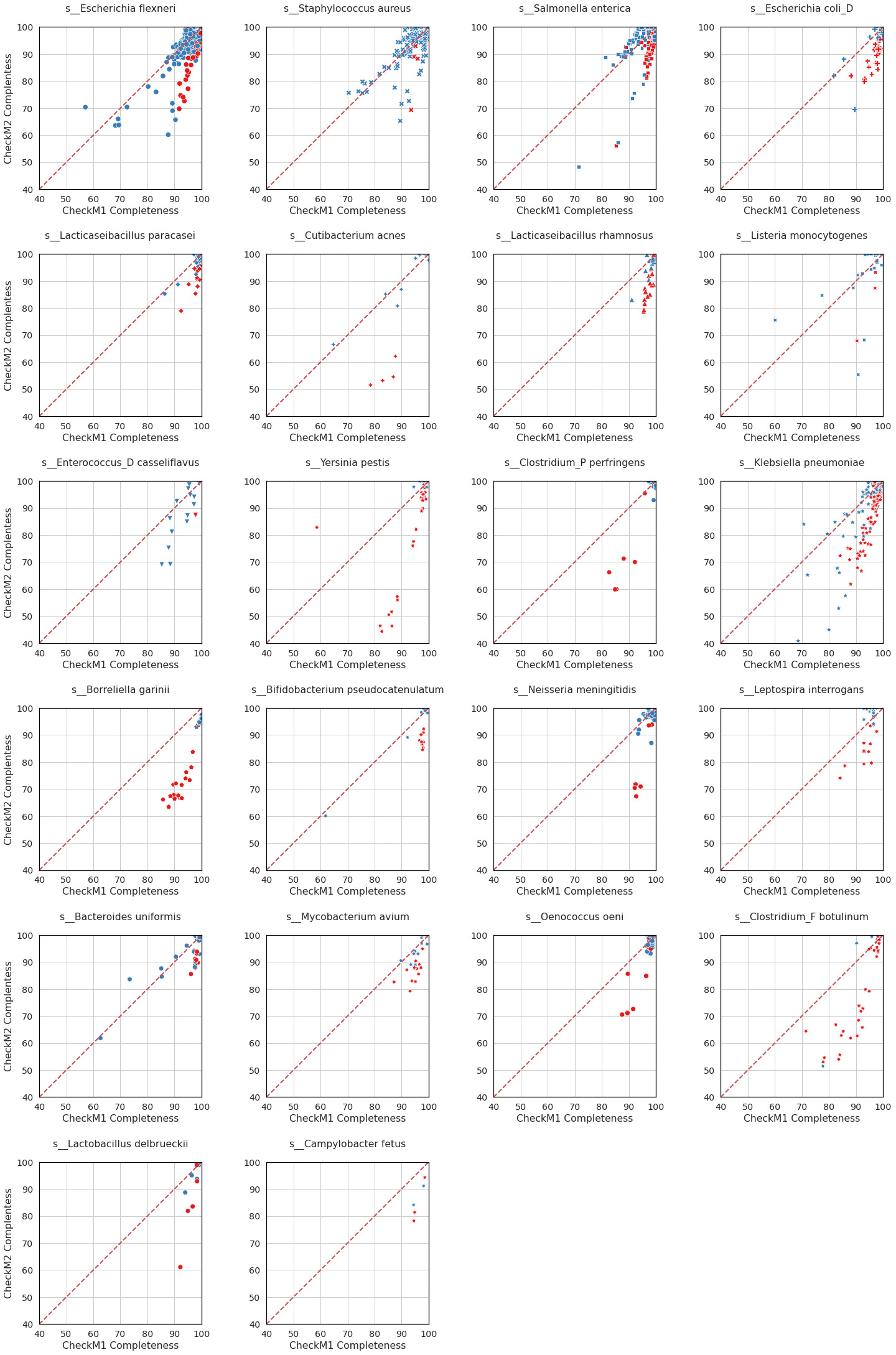

**Figure 9:** Genome completeness predictions of CheckM1 vs CheckM2 for selected bacterial species with genomes containing hypothesized indels as identified by NCBI metadata (marked in red).

In addition to genomes annotated as having indels by NCBI, there are likely further genomes containing indels not flagged by NCBI (**Figure 10**). The NCBI annotation of suspected indels relies on a simple CDS count deviation from a mean, which will likely miss some indel-dominated genomes with a lower CDS count or species with high number of representative genomes and a higher CDS count variance. Notably, for many genomes with suspected indels, there is a clear linear regression pattern of higher protein count being associated with lower coding density relative to the bulk of the genomes **(Figure 10)**. As such, there are likely genomes not identified by NCBI metadata as containing many frameshifted proteins (in blue), which nevertheless are substantially higher in protein count and lower in coding density, such as those the highly deviated genomes of *Salmonella enterica*, *Staphylococcus aureus* and *Escherichia flexneri* **(Figure 10)**. Notably, these are also associated with a substantial difference between CheckM1 and CheckM2 predictions and may likely also contain many indels.

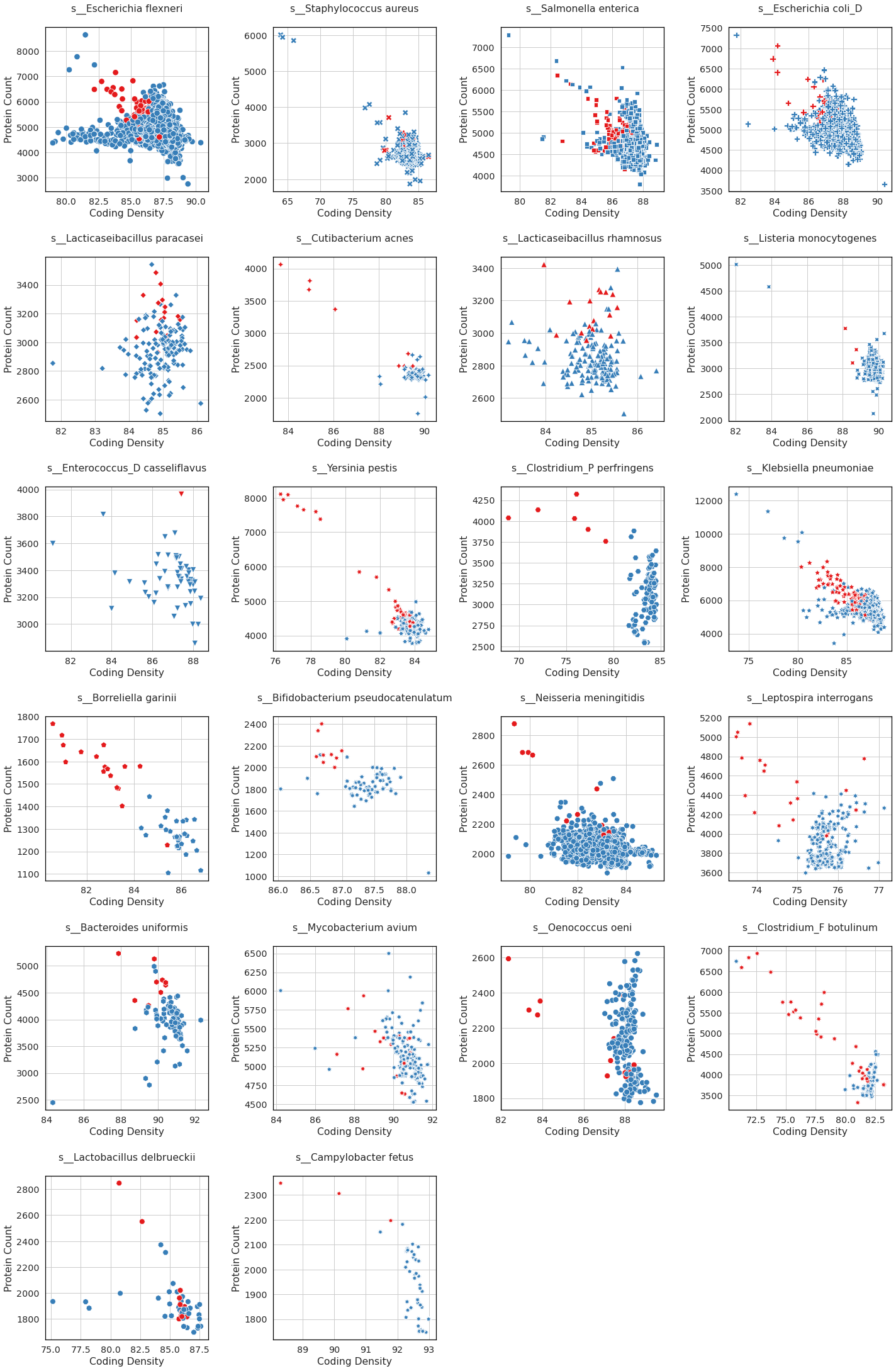

**Figure 10:** Coding density versus protein count for selected bacterial species with genomes containing hypothesized indels as identified by NCBI metadata (marked in red)

In addition to the species above, a high number of frameshifted genomes as indicated by NCBI metadata (> 50) were generally found in the phyla Spirochaetota, Bacteroidota, Firmicutes, Firmicutes_A, Actinobacteriota, Campylobacterota and Proteobacteria. Notably, these phyla show a group of genomes where CheckM1 predictions appear to be much higher than CheckM2, further supporting the hypothesis that the presence of indels significantly affects CheckM2 predictions (**Figure 11**).

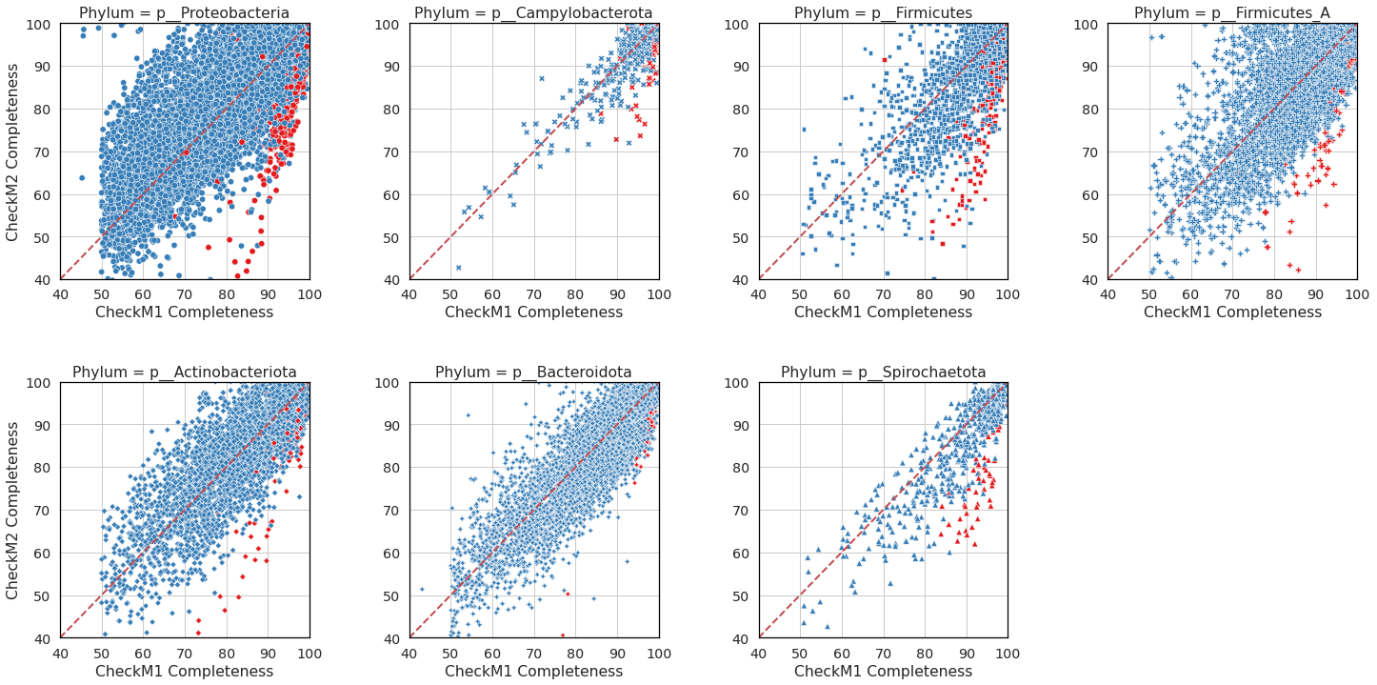

**Figure 11:** Phyla in GTDB Release 202 with > 50 genomes annotated as frameshifted by NCBI metadata. Red indicates genomes with suspected indels.

As indels affect CDS count and protein annotation, they also affect biological conclusions drawn from genomes with high number of indels. Therefore, CheckM2 scores are more reflective of the underlying quality of the genomic assembly and lead to more robust biological conclusions.

**Supplementary Note 7**: One-sided sampling effects on divergent CheckM1 and CheckM2 of MAGs from GTDB release 202 where CheckM1 predictions are higher

Curation and submission of MAGs is often guided by their CheckM1 scores. This can create a selection effect where noise error distribution is sampled asymmetrically. For example, error plots in **Figure 12a** show the distribution of CheckM1 and CheckM2 completeness predictions relative to the known true value in the phylum Spirochaeotota (based on simulated genomes created using the random MAG derived fragmentation on genomes from RefSeq release 202). CheckM2 errors are on average substantially lower than CheckM1 errors which has relatively high overprediction and underpredictions.

Applying a 50% CheckM1-based completeness cut-off (as used for GTDB) results in an asymmetrically sampled prediction distribution where substantial underpredictions are filtered out while substantial overpredictions are retained. As a result, more CheckM2 predictions appear to be lower relative to CheckM1 (**Figure 12b**). When the filtered-out genomes are added in, the linear relationship between the CheckM1 and CheckM2 completeness scores is much clearer (**Figure 12c**).

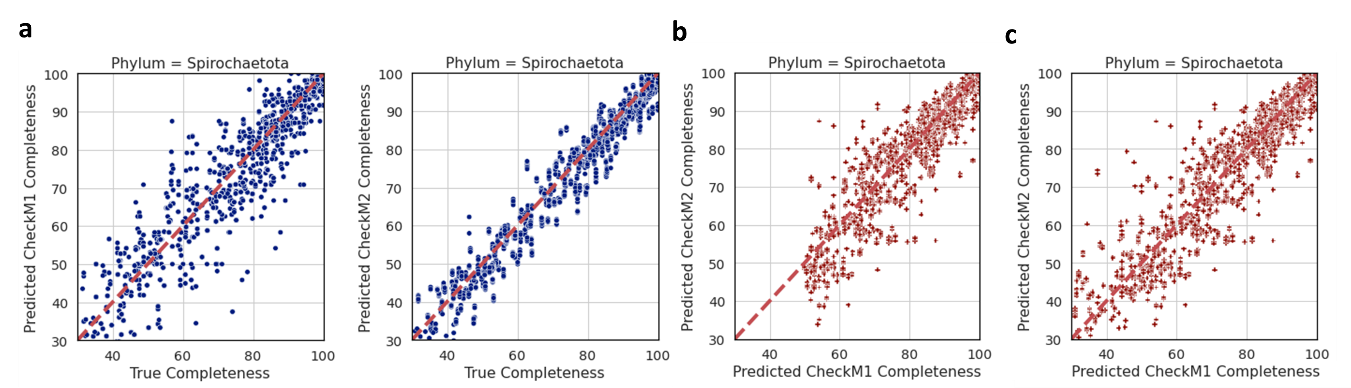

**Figure 12**: Completeness predictions by CheckM1 and CheckM2 on synthetic Spirochaeotota genomes from RefSeq release 202. **a)** CheckM1 and CheckM2 predictions plotted against the true completeness value. **b)** CheckM1 and CheckM2 predictions plotted against each other for only those genomes that pass a CheckM1-based 50% completeness cutoff. **c)** CheckM1 and CheckM2 predictions plotted against each other for all synthetic genomes.

Further evidence may be obtained by looking at all binned MAGs in a given environment instead of only those selected for publication or submission. As an example, Acidobacteriota, Actinobacteriota, Bacteroidota, Chloroflexota and Verrucomicrobiota are all phyla ubiquitous in many environments, including in the permafrost soil as outlined in Woodcroft et al., 2018^3^. Plotting the completeness predictions of all metagenomic bins recovered from samples in Woodcroft et al. belonging to these phyla without any quality cut-offs, CheckM1 and CheckM2 predictions are roughly linearly distributed **(Figure 13a)**. Nevertheless, instituting a commonly used cut-off of CheckM1 completeness > 70% and CheckM1 contamination < 10% produces asymmetrically sampled predictions where is appears that CheckM2 is underpredicting more relative to CheckM2 on average **(Figure 13b)**. It is therefore likely that at least some apparent biases where CheckM1 completeness predictions are higher than CheckM2 are a result of using CheckM1-based quality cutoffs, as these will always bias towards erroneously overpredicted scores compared to erroneously underpredicted scores.

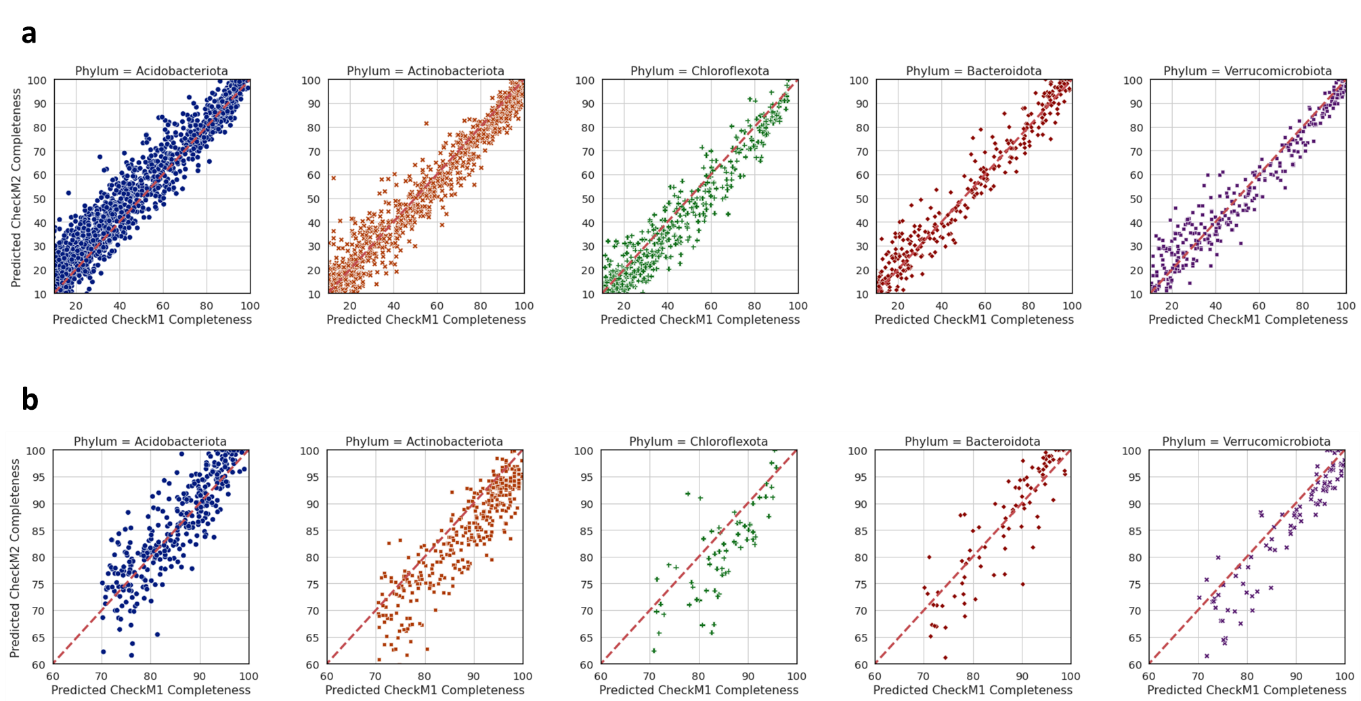

**Figure 13**: CheckM1 and CheckM2 completeness predictions for all metagenomic bins of the phyla Acidobacteriota, Actinobacteriota, Bacteroidota, Chloroflexota and Verrucomicrobiota derived from ^3^. **a)** All predictions for all bins are plotted, **b)** predictions are plotted only for those bins where CheckM1 completeness is > 70% and CheckM1 contamination is < 10%.

**Supplementary Note 8:** Examples of contamination differences between CheckM1 and CheckM2 in GTDB release 202

We assessed some example cases where contamination was either significantly higher or lower when comparing CheckM2 and CheckM1 estimates. Two such examples include the Nanoarchaeota genomes GCA_002690445.1 and GCA_002686195.1, predicted by CheckM1 to be free of contamination, but 25% contaminated by CheckM2. These genomes are unusually at large 4 and 6.6 Mbp respectively (average genome size across 88 GTDB genomes of the order Pacearchaeales is 861 Kb). CheckM1 uses the universal archaeal marker set (120 genes) on these genomes, which did not find any duplicated genes, while CheckM2’s contamination model used genes including a number of biologically important genes such as DNA polymerase I, DNA ligase I, DNA helicase and dUTP pyrophosphatase that were present in 2/6, 4/4, 2/5 and 7/5 copies respectively (K02335, K10747, K02314 and K01520; **Supplementary Table 8**) – which are present in all Nanoarchaeota genomes in GTDB release 202 at a mean count of 0.04, 1, 0.04 and 0.6 respectively, and are therefore highly elevated in copy number in the two Nanoarchaeota genomes with high suspected contamination.

In another case, CheckM2 predicts that the isolate archaeal genome of *Halogranum amylolyticum* (GCF_900110465.1; 5.18 Mbp) is 99% complete and 10.7% contamination, while CheckM1 predicted the same completeness but 1.7% contamination. Re-assembly and binning of this genomic data yielded two discrete genomic bins (3.32 Mbp and 1.28 Mbp) with distinct GC composition. The new 3.32 Mbp genome bin was predicted to be > 99% complete by CheckM1 and CheckM2, and only 0.25% contaminated by CheckM2 (1.33% contaminated by CheckM1). Analysis of the SHAP values contributing to the high initial contamination score showed that the majority of genes contributing towards a high contamination prediction are genes present in both bins (unexpected high gene copy number), or present only in the smaller bin (unexpected genes). This shows that CheckM2’s approach is able to detect contamination in a wider variety of circumstances than CheckM1.

In another case, the genome of *Omnitrophus magneticus* (GCA_000954095.1) is predicted to be 50% contaminated by CheckM1 but only 5% contaminated by CheckM2. This genome is a result of single-cell amplification (MDA amplification), which can sometimes result in a very high amplification and inflated copy number of a small number of genes relative to the overall genome^4^. Notably, CheckM2 identifies that 4 usually single-copy genes are present in more than ten copies; but does not weight those very high as most other single-copy genes are present in only one or two copies. On the other hand, CheckM1’s score is strongly influenced by the presence of a small number of single-copy marker genes in high copy number. As CheckM1 relies on the assumption that duplicate marker genes are a fair sample of the overall contamination, it is likely that CheckM2’s weighting is more accurate.

An extreme example of this bias can be seen in a simulation where a single single-copy marker gene (Ribosomal protein L14) was inserted into an *E. coli* genome 1, 5, 10, 20 and 100 times to simulate uneven genome amplification and investigate its effects on contamination prediction by CheckM1 and CheckM2 (**Table 3**). As can be seen, CheckM1 is strongly affected by this process, predicting 28% contamination for a genome where the actual contamination fraction is only 1.5%. CheckM2 on the other hand is almost insensitive to this high amplification of a single gene, yielding overall a much more accurate contamination prediction for the whole genome relative to its contamination definition (% of base pairs not belonging in the original genome sequence).

| Ribosomal protein L14 (*E. coli*) count | CheckM1 Contamination | CheckM2 Contamination | Actual Contamination (bp percentage) |
| --- | --- | --- | --- |
| 100 | 28.06 | 0.58 | 1.52 |
| 20 | 3.13 | 0.57 | 0.30 |
| 10 | 1.57 | 0.57 | 0.15 |
| 5 | 0.63 | 0.57 | 0.07 |
| 1 | 0 | 0.57 | 0.01 |

**Table 3:** Prediction of contamination by CheckM1 and CheckM2 compared to actual contamination of a gene present in 1, 5, 10, 20 and 100 copies.
